# Rapid Assessment of Chemical Complementarity of Ligands for Protein Design

**DOI:** 10.1101/2025.06.30.662286

**Authors:** Rokas Petrenas, Katarzyna Ożga, Joel J. Chubb, Andrey V. Romanyuk, Dominic Alibhai, Jennifer J. McManus, Graham J. Leggett, Nigel S. Scrutton, Thomas A. A. Oliver, Derek N. Woolfson

## Abstract

Driven by deep-learning methods, computational protein design can now rapidly generate *de novo* structures, opening new frontiers for creating proteins that bind small molecules tightly and specifically, although achieving predictable and tunable binding remains challenging. Here we address this with a rapid physics-based computational method to generate isosteric and chemically complementary binding pockets for small-molecule targets in *de novo* designed proteins. We test this experimentally by constructing and characterizing binding proteins for several synthetic and natural chromophores. By evaluating only single-digit numbers of designs, the pipeline delivers stable proteins with pre-organised binding sites in apo structures confirmed by X-ray crystallography, which bind the targets with micromolar affinities or better. To illustrate the scope and applications of this approach, we incorporate selective and coupled chromophore-binding sites in a two-domain *de novo* protein enabling controlled energy transfer between the two sites, and we develop a small *de novo* binding protein that can be used in live mammalian cells to visualise sub-cellular structures.

## Introduction

For some time, protein designers have explored the modelling and construction of small molecule-binding sites in natural and *de novo* protein scaffolds to enable functions like sensing and catalysis^1,2^. Such studies have highlighted the difficulty of achieving high-affinity and highly specific interactions. The routes taken have encompassed all approaches in *de novo* design, including minimal^3^, rational^1,4,5^, and computational strategies^6^. Many of these focus on specific ligands rather than small-molecule binding in general and the need to validate selectivity. For example, using computational design, Baker et al. have used Rosetta to computationally design binders for the steroid digoxigenin into a natural scaffold, initially achieving low-to-mid micromolar affinity, and then improving this to picomolar with library-based mutagenesis and selection^6^. Kortemme et al. have expanded the scope of computational binding-site design by fragmenting ligands and using the fragments in PDB searches to identify optimal contacts^7^. Polizzi and DeGrado combine fragment-based binding-site design with parametric scaffold generation to develop an apixaban binder with sub-micromolar affinity, and low-nanomolar binders for rucaparib^8-10^. Also, Höcker and Jürgens have engineered the *E. coli* tryptophan repressor to bind auxin as a component of a biosensor in plants^11^. Despite these and other successes, these largely physics- and knowledge-based methods are computationally slow, can have limited ligand scope, and usually require careful input and filtering from an expert user^12^.

Currently, protein design is undergoing a step-change with the application of deep-learning methods^12,13^. While AI has accelerated and simplified the sampling of designable *de novo* binders, it is not yet clear how well these tools predict which designs are most likely to succeed experimentally or how they advance mechanistic understanding of protein function. Specifically, early applications of deep-learning methods to deliver small molecule-binding proteins, whilst achieving low-micromolar to nanomolar affinities for cholic acid and methotrexate for instance, still require experimental screening of large numbers of designs per target^14^. New pipelines based on scaffold generation using RFDiffusionAA^15^ do show improvement for targets that are well represented in structural databases, such as binding pockets for haem or bilin. Whilst impressive advances, these studies highlight the considerable challenges that remain in the computational design of small molecule-binding proteins, with even the most advanced methods requiring extensive optimization and screening to achieve experimental successes^12,16^.

To address these issues, we introduce RASSCCoL, a computational pipeline for the Rapid ASSessment of Chemical Complementarity of Ligands in protein design. This is a robust and interpretable physics-based method for incorporating small molecule-binding sites into protein scaffolds. RASSCCoL’s streamlined pipeline uses the volume and functional groups of the ligand to etch complementary pockets into the proteins. Two key aspects of this approach are: (i) the use of fast volume calculations to limit the number of side-chain combinations that need to be assessed, and (ii) the leveraging of robust, highly stable *de novo* designed α-helical scaffolds. Effectively, these allow near-exhaustive sampling of sequences that are most likely to generate pre-organised, isosteric binding sites for the target ligand while bypassing the sampling of backbone flexibility. The resulting sequences are repacked onto the input backbone and are evaluated by ligand docking using AutoDock Vina^17^, and then by regenerating the protein folds and binding pockets using AlphaFold2 predictions^18^. To address the challenge of exponential scaling for larger ligands and concomitant sequence spaces, we employ machine learning to score designs at the sequence level. This restricts the computationally expensive explicit modelling of structures to the most-promising sequences.

To validate this approach, we create binding sites in single-chain coiled coil-based scaffolds generated through rationally seeded computational protein design^19,20^. These scaffolds have advantages of small size, stability, and well-understood sequence-to-structure relationships. Specifically, they have layered hydrophobic cores that facilitate ligand alignment and side-chain replacement in the computational design. Coiled-coil proteins and protein-protein interfaces are ubiquitous in nature^21^, and are amongst the most successful *de novo* designed proteins targeted to date^22,23^. In principle, however, RASSCCoL can be applied to any protein scaffold. Combined, this pipeline simplifies the design process, requires minimal user input, produces robust *in silico* models that are chemically derived and interpretable, limits the number of constructs for experimental testing, and yields high success rates of confirmed binders. Importantly, this successful rational and physical approach provides a baseline for increasingly complex design pipelines. Finally, we show that RASSCCoL can be used to introduce orthogonal but coupled binding sites to produce more-complex, functional *de novo* proteins; and to deliver small *de novo* chromophore-binding proteins for use as fluorescent reporters in cells.

## Results

### Rapid physics-based design pipeline

Initially to develop and illustrate RASSCCoL, we chose a compact and stable *de novo* designed 4-helix bundle, sc-apCC-4 (PDB id 8A3K), as the protein scaffold^19^; and two synthetic, environment-sensitive dyes—1,6-diphenyl-1,3,5-hexatriene (DPH) and Nile red—as the target ligands^24^. Our computational pipeline for generating small molecule-binding sites had three generalisable steps (Fig. 1): (i) binding-site selection; (ii) sequence generation and culling; (iii) model building, ligand docking, and evaluation.

**Fig. 1:**
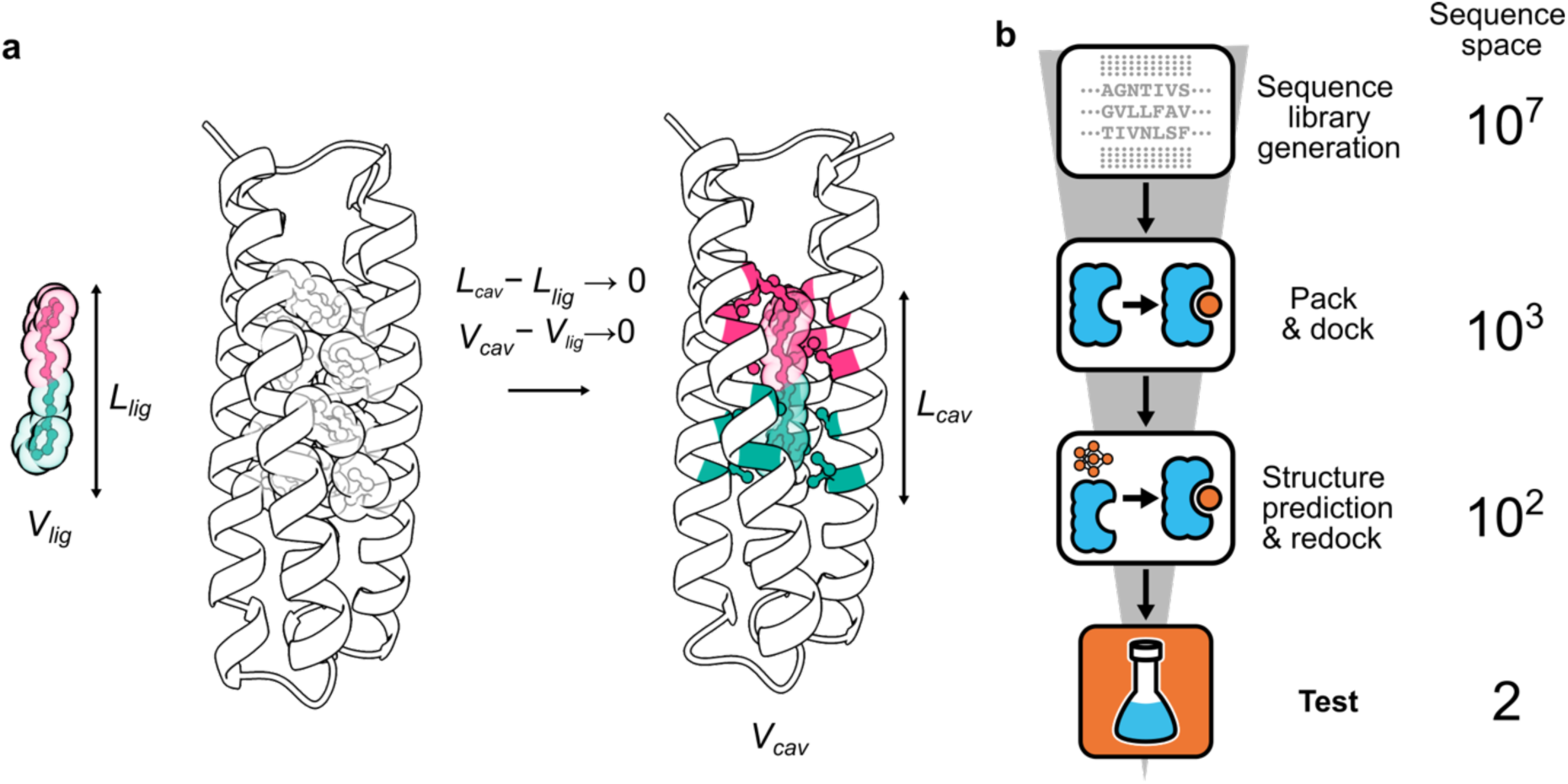
The RASSCCoL computational protein design pipeline illustrated by DPH binding. **a,** A potential site for DPH binding (*L_cav_*) is identified based on the maximum ligand length (*L_lig_*). To match the ligand’s shape, the protein scaffold is segmented into corresponding layers (magenta and green). A volume constraint is used to minimise the difference between the binding cavity (*V_cav_*) and the ligand (*V_lig_*) in each layer; effectively, replacing atoms of the side chains with those of the ligand. **b,** Design and evaluation strategy showing the numbers of sequences at each step of DPH-binder design. In this case, the sequence library allows only aliphatic amino acids in the binding pocket. Sequences passing the volume constraint are packed onto the scaffold for ligand docking. Structures with the most-negative predicted binding energies in AutoDock Vina are modelled using AlphaFold2 (AF2) and the ligands redocked. Two high-confidence designs with the best redocking scores are selected for experimental testing.

First, the positions of binding-site residues were identified based on ligand length (Supplementary Fig. 1). The core packing in sc-apCC-4 is layered with four side chains per layer and a spacing of ≈5 Å between layers. Thus, the number of layers that need to be redesigned to accommodate a ligand can be estimated as maximum ligand length in Å divided by 5 (Fig. 1a). Second, a volume calculation based on Bondi radii^25^ was used to identify side-chain permutations to generate voids that match the ligand’s volume most closely. The 2-fold symmetric and entirely hydrophobic ligand DPH (length, L ≈ 15 Å; and volume, V ≈ 274 Å^3^; Fig. 1a) spans three 4-residue layers of sc-apCC-4. Within these layers, we retained some of the Ile residues important for fold stability^19^. This left ten designable positions in total, which we restricted to combinations of the aliphatic residues, Gly, Ala, Val, Leu and Ile (Supplementary Table 1 and Supplementary Figs. 1 – 2). Thus, potentially, ≈10 million sequence combinations (5^10^) could have been searched. However, this reduced to ≈3,200 sequences by applying the ligand-volume and ligand-symmetry constraints (Fig. 1b, Supplementary Table 1). Third, using FASPR^26^ each sequence was repacked into the scaffold structure, and DPH was docked into the resulting models using AutoDock Vina (Fig. 1b)^17^, treating the receptor as rigid. AutoDock Vina has a physics-based scoring function to predict ligand-binding poses and affinities, which we used to score the models. While it does not explicitly model solvent effects, these are partially and implicitly accounted for through the parameterisation of the scoring function. We selected AutoDock Vina for its computational efficiency and suitability for high-throughput screening. Combined, these computational steps took tenths of a second for each sequence (Supplementary Fig. 3), enabling the entire design protocol to finish within 30 minutes on a single CPU core (Supplementary Table 1). Finally, AlphaFold2 (AF2) was used to predict structures for a small number of the top-scoring sequences, followed by docking to evaluate ligand-pose recovery (Fig. 1b, Supplementary Table 2). These provided assessment metrics to choose only a small number of designs for experimental production and characterisation (Supplementary Fig. 4, Supplementary Table 3). To evaluate how RASSCCoL extended to asymmetric ligands, we targeted Nile red (L ≈ 13 Å; V ≈ 324 Å³; Supplementary Fig. 2). In this case, the sequence space was significantly larger at ≈400,000 possible sequences after applying the volume constraint (Supplementary Table 1, Supplementary Fig. 5). By parallelizing this approach to utilise multiple CPU threads, the computations for Nile red were completed in less than 2 hours.

Thus, RASSCCoL enables *in silico* screening of hundreds of thousands of designed sequences within hours even with modest computational resources (Supplementary Table 1, Supplementary Fig. 2). This allows users to embellish the RASSCCoL pipeline as needed, for instance to include short molecular dynamics (MD) simulations to explore wider backbone and side-chain conformational space (Supplementary Figs. 9 – 10).

### Robust design of small-molecule binders

For experimental testing, we targeted five ligands of varying volume, polarity, and heteroatom content (Fig. 2a, Supplementary Fig. 2): the aforementioned hydrophobic fluorophores DPH (L ≈ 15 Å; V ≈ 274 Å³) and Nile red (L ≈ 13 Å; V ≈ 324 Å³); more-hydrophilic and biologically relevant lumiflavin (L ≈ 11 Å; V ≈ 257 Å³); and biologically active arcyriaflavin A (L ≈ 12 Å; V ≈ 310 Å³) and SN38 (L ≈ 14 Å; V ≈ 396 Å³). For each ligand, we introduced up to ten mutations into three layers of the sc-apCC-4 scaffold (Fig. 2b and Supplementary Table 1). For lumiflavin, arcyriaflavin A and SN38, the side-chain libraries were diversified to include small polar residues, Ser, Thr, Asn, and Asp (Supplementary Table 1). In this way, we aimed to accommodate the polar functional groups of these ligands, hypothesizing that these would be captured by the AutoDock scores in the RASSCCoL pipeline (Supplementary Figs. 6 – 9). Indeed, binding-site sequences with polar amino acids were returned for these ligands (Fig. 2b, Supplementary Fig. 11). RASSCCoL calculations and full-length sequences of designs made as synthetic genes are summarised in Supplementary Tables 1 and 3.

**Fig. 2:**
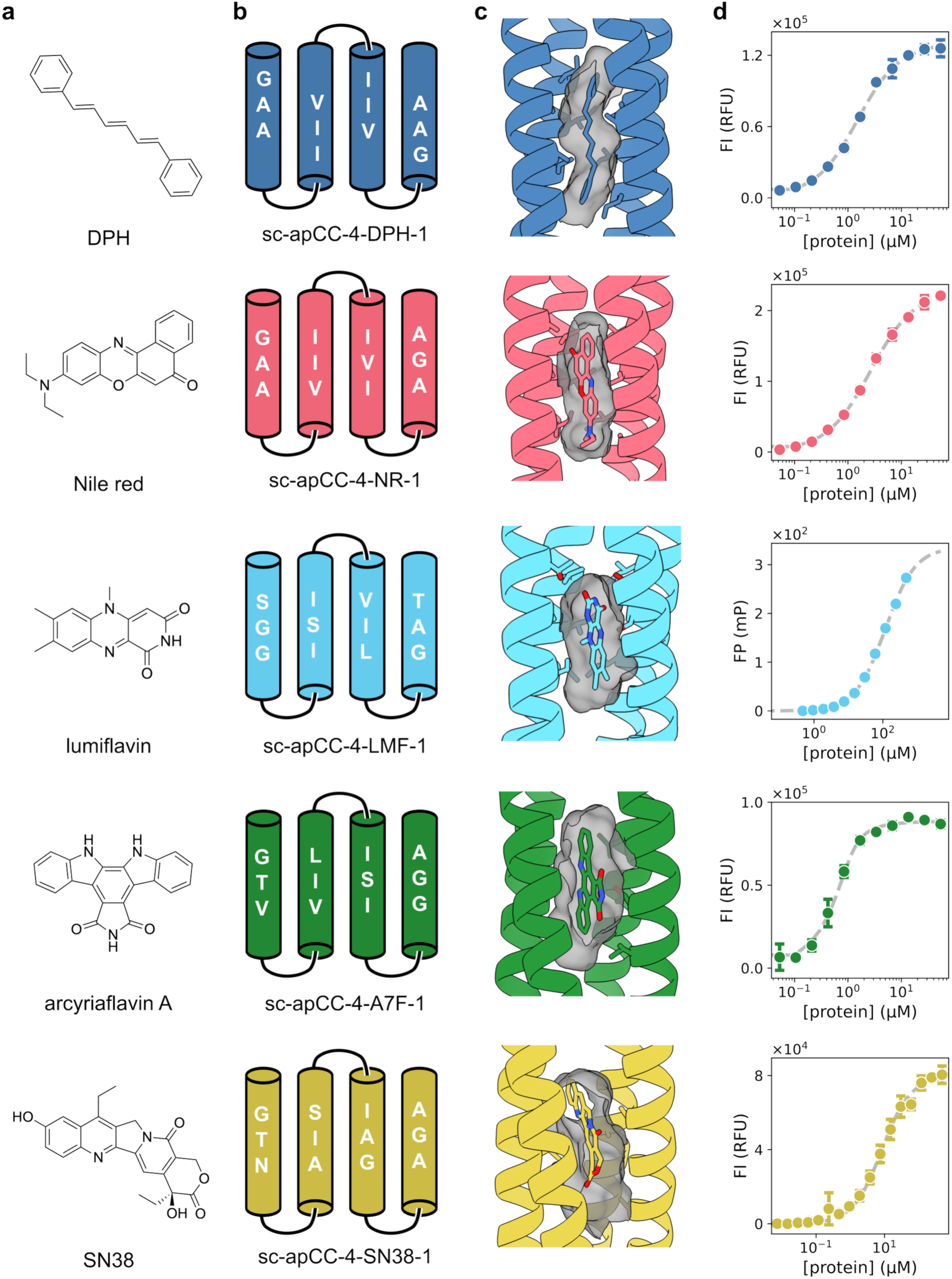
RASSCCoL generates low-μM binders to a range of ligands in a starting scaffold devoid of cavities. **a,** Ligands tested for binding, top to bottom: DPH (blue-coloured panels), Nile red (red), lumiflavin (cyan), arcyriaflavin A (green), and SN38 (yellow). **b,** Binding-site residues within the hydrophobic core of the 4-helix bundle, sc-apCC-4 (PDB id 8A3K). **c,** Structure predictions of the binders from AF2 with ligands docked using Autodock Vina. **d,** Saturation-binding data measured by fluorescence intensity (FI) or fluorescence polarization (FP). Fitted dissociation constants, top to bottom: K_D(DPH)_ = 1.2 ± 0.1 μM; K_D(Nile red)_ = 2.4 ± 0.1 μM; K_D(lumiflavin)_ > 50 μM; K_D(arcyriaflavin A)_ = 0.11 ± 0.02 μM; K_D(SN38)_ = 7.8 ± 0.6 μM. Conditions: 0.5 μM DPH/Nile red or 1 μM lumiflavin/SN38/arcyriaflavin A, with either 10% v/v MeCN and 0 – 60 μM protein (DPH/Nile red/arcyriaflavin A) or 10% v/v DMSO and 0 – 480 μM protein (lumiflavin/SN38). DPH/Nile red/arcyriaflavin A/lumiflavin were measured in PBS, pH 7.4. SN38 was measured in ABS, pH 4. The data shown are the mean of n=3 independent repeats, with error bars representing the standard deviations from the means. For simplicity, only the highest-affinity protein for each ligand is shown, see Supplementary Figs. 17 – 21 for the other designs. Source data are provided as a Source Data file.

To test the robustness of both RASSCCoL and the sc-apCC-4 scaffold, we selected just thirteen sequences for characterization based on the docking scores: two for DPH, with clockwise and anticlockwise arrangements of the helices predicted by AF2 (Supplementary Fig. 12) but with the same binding-site solution; two each for Nile red and arcyriaflavin A; three for lumiflavin; and four for SN38 (Supplementary Table 3 and Supplementary Figs. 4 – 9). All proteins expressed well, were monomeric, α helical, and thermally stable (Supplementary Figs. 13 – 16). Binding assays using fluorescence intensity (FI) or fluorescence polarization (FP) measurements returned sub-micromolar affinities for DPH and arcyriaflavin A, low-micromolar affinities for Nile red and SN38, and an order of magnitude weaker > 50 μM affinity for lumiflavin (Figure 2d, Supplementary Figs. 17 – 21). In retrospect, our best explanation for the lower affinity of lumiflavin is that it is least lipophilic ligand in our set (Supplementary Table 4) and has the smallest hydrophobic surface area.

Interestingly, and although ligand selectivity was not explicitly included at this stage in the RASSCCoL pipeline, we found that several of the designed proteins discriminated between the different small molecules (Supplementary Table 5, Supplementary Fig. 22). Notably, the sc-apCC-4-DPH-1 and sc-apCC-4-NR-1 proteins, with designed hydrophobic binding pockets, did not bind the more hydrophilic lumiflavin—despite its smaller molecular volume compared to DPH and Nile red—consistent with the lack of chemical complementarity (Supplementary Table 5, Supplementary Fig. 22).

To explore the design space further, we tested if the top RASSCCoL hits could be optimised through sequence redesign using LigandMPNN^27^ and protein:ligand complex prediction with Boltz2^28^. After two design rounds, Boltz2 ipTM scores plateaued at 0.97 – 0.99, and binding-site sequences converged (Supplementary Table 6, Supplementary Figs. 24 – 28). Experimentally, we tested two designs each for DPH, Nile red, lumiflavin and arcyriaflavin A, and four for SN38 (Supplementary Table 7). These had up to 9 mutations at the ***a*** and ***d*** core positions within the binding site relative to the original RASSCCoL designs (Supplementary Figs. 24 – 28). All proteins expressed, and 10/12 were monomeric, α helical, and thermally stable (Supplementary Figs. 29 – 30). Similarly to the original RASSCCoL designs, one each of the DPH- and Nile red-binding variants bound the target with sub-micromolar affinity, and the lumiflavin binders showed >50 µM binding affinity (Supplementary Figs. 31–33). However, no detectable binding to SN38 was observed (Supplementary Fig. 34). Notably, arcyriaflavin A was the only targeted ligand for which the revised procedure returned a design with a marked improvement, binding with <2 nM affinity (Supplementary Fig. 35). Thus, these results indicate that additional sequence optimisation does not always improve affinity and can, in some cases, be detrimental, despite substantially higher computational cost. Therefore, for the remainder of the study, we evaluated and used the RASSCCoL-only designs.

As controls to test the robustness of the scaffold and any ligand binding through non-specific interactions, we mutated the targeted binding site residues to either all Gly or all Ala to give sc-apCC-4-Gly and sc-apCC-4-Ala, respectively. Despite high AF2 confidence (Supplementary Table 2), the sc-apCC-4-Gly was completely unfolded (Supplementary Fig. 12, top). However, the sc-apCC-4-Ala mutant was monomeric, α helical, and thermally stable (Supplementary Figs. 13 and 16). It bound target ligands at least one order of magnitude weaker than the best designed binders (Supplementary Table 5, Supplementary Figs. 17 – 21). Interestingly, arcyriaflavin A was the most promiscuous ligand in our panel, and it was the only ligand to bind the Ala-based control with µM affinity.

We determined high-resolution crystal structures of the ligand-free forms for all five *de novo* binders. (Fig. 3, Supplementary Tables 8 – 9). The DPH binder crystalised as a domain-swapped dimer despite being monomeric in solution (Supplementary Figs. 15 and 36 – 37)^29,30^. Nonetheless, both domains aligned well with the AF2 model, with the designed binding site geometry maintained (Fig. 3a). The Nile red, lumiflavin, arcyriaflavin A and SN38 binders matched the respective AF2 models closely with overall root-mean square differences between the C_a_ atoms (C_α_-RMSDs) of < 0.5 Å (Fig. 3a). All five apo-protein structures retained their overall backbone conformations (Supplementary Fig. 38) and revealed solvent-accessible pockets. The side-chain geometries matched those predicted by AF2 (Fig. 3a) or initial repacking with FASPR (Supplementary Figs. 39 – 44), with binding-site heavy-atom RMSDs of <0.5 Å (Supplementary Fig. 45). Thus, the observed cavity shapes and volumes matched the computational designs closely. Moreover, we found that the most computationally efficient approach of side-chain repacking on a rigid backbone achieved comparable accuracy to more computationally intensive methods (Supplementary Figs. 39 – 44).

**Fig. 3.**
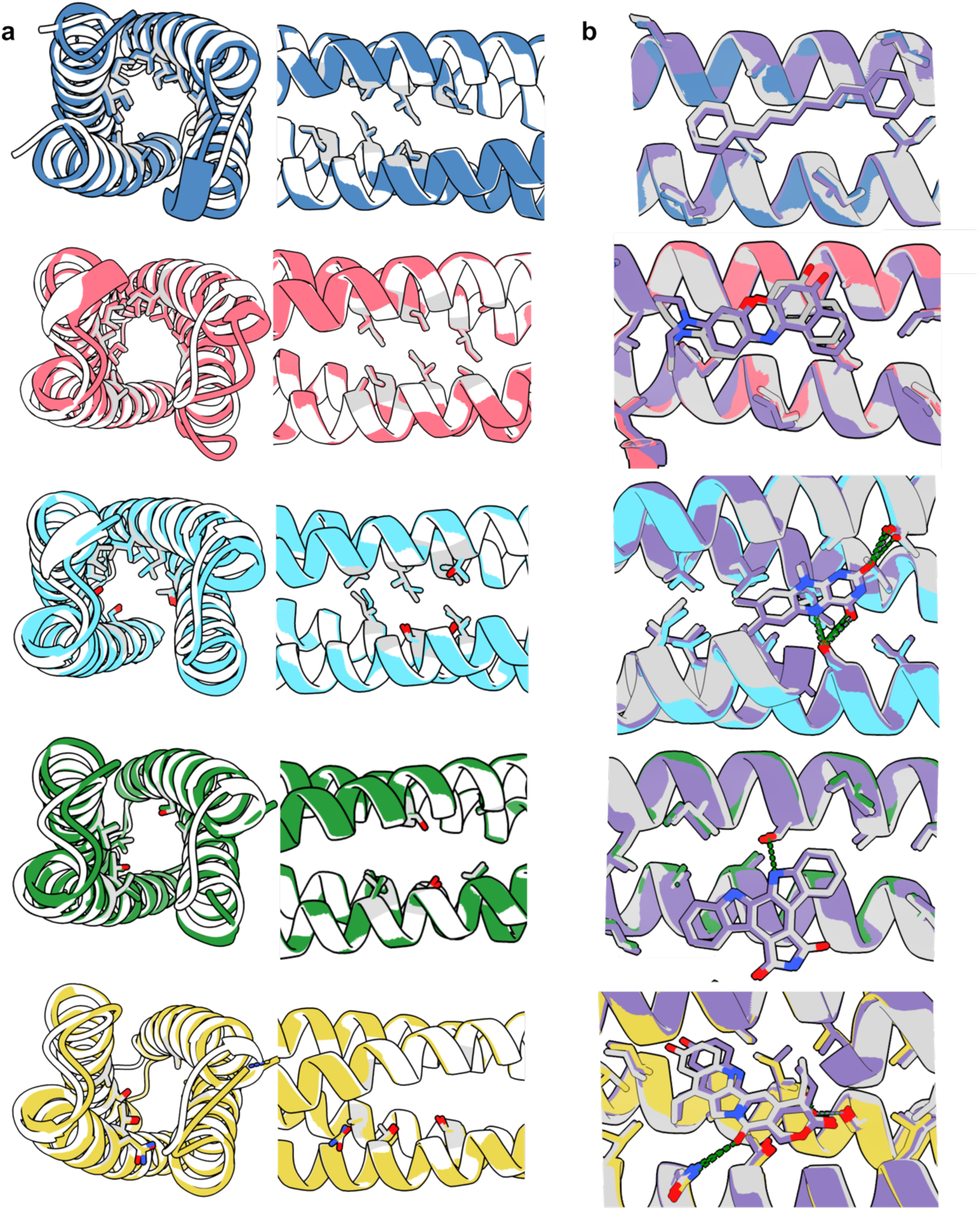
Binding-site models generated by RASSCCoL closely match ligand-free crystal structures. **a,** Binding-site heavy-atom alignments comparing AlphaFold2 models (grey), with the experimental X-ray crystal structures (coloured as in Fig. 2). The DPH binder, sc-apCC-4-DPH-2, at 2.4 Å resolution (with a root-mean square difference between the side-chain heavy atoms (sc-RMSD_AF2_) = 0.45 Å, PDB id 9R1N). Only one domain of the domain-swapped dimer is shown, see Supplementary Fig. 37. The Nile red binder, sc-apCC-4-NR-1, at 1.7 Å resolution (sc-RMSD_AF2_ = 0.33 Å, PDB id 9R1J). The lumiflavin binder, sc-apCC-4-LMF-1, at 2.4 Å resolution (sc-RMSD_AF2_ = 0.30 Å, PDB id 9R1M). The arcyriaflavin A binder, sc-apCC-4-A7F-1, at 2.3 Å resolution (sc-RMSD_AF2_ = 0.35 Å, PDB id 9R1K). The SN38 binder, sc-apCC-4-SN38-1, at 1.85 Å resolution (sc-RMSD_AF2_ = 0.38 Å, PDB id 9R1L). **b,** Slices through the bundles showing binding-site heavy-atom alignments comparing Boltz2 models with ligands (grey), and RASSCCol repacked and redocked models (pale purple) with the experimental X-ray crystal structures. Binding-site residues are highlighted and shown as sticks, with oxygen atoms in red and nitrogen atoms in blue. Hydrogen bonds are shown as dashed green lines. For sc-RMSD to Boltz2 and RASSCCoL repacked and redocked models, see Supplementary Fig. 45.

Although we were unable to obtain crystals of the ligand:protein complexes, modelling with Boltz2^28^ produced high-confidence binding poses consistent with the design models and binding pockets observed in the ligand-free crystal structures, including predicted hydrogen bonds between the ligands and the introduced polar side chains (Fig. 3b).

### RASSCCoL scales to larger sequence spaces

Traditional physics-based protein design methods that explore combinations of side chains do not scale well with increasing number of designable positions and large amino-acid palettes^31-33^. For our *de novo* scaffolds, significant reduction of the sequence space is possible by exploiting the symmetry, layered structures, and well-understood sequence-to-structure relationships of α-helical bundles. However, in more general cases using RASSCCoL, even with its optimised and relatively fast model-building and docking steps, the process would become computationally expensive once sequence space exceeded a few million. To mitigate this, we implemented an active sequence sampling strategy, prioritising the most-informative data points rather than exhaustively docking all possible sequences, and significantly reducing computational cost and time (Supplementary Figs. 46 – 51).

We benchmarked several computationally efficient machine learning models used in active learning, including gradient-boosted trees (GBT), random forests (RF), and linear models, on two datasets containing 26 and 21 million sequences, respectively (Supplementary Figs. 46 – 47). Briefly, the models were trained initially on a small (50,000) randomly selected subset of sequences and used to predict docking scores for the remaining sequences. The training set was then expanded iteratively by selecting sequences based on the highest prediction uncertainty (i.e., knowledge gaps) and the lowest predicted docking scores (i.e., likely binders), which were evaluated subsequently by explicit docking. This process was repeated until no further improvement in the model’s ability to retrieve the top 1% of sequences from an unseen test set was observed. All models required training on <5% of the sequence space to converge, demonstrating rapid active learning on these large datasets (Supplementary Figs. 48 – 51). At convergence, the remaining sequences were docked to evaluate model performance. Among the models tested, GBT consistently performed best, recovering 92% and 99% of the top 2,500 sequences in the two datasets with over 10-fold reduction in computational cost compared to brute-force docking (Supplementary Figs. 49 and 51). This enables exploration of tens of millions of sequences within hours.

To test this approach experimentally, we used a recently designed antiparallel 6-α-helix barrel, sc-apCC-6 (PDB id 8QAD)^20^. Unlike sc-apCC-4, this has a central hydrophobic cavity that binds DPH with low-micromolar affinity but has a weaker affinity for Nile red (Supplementary Fig. 52). We aimed to reverse this selectively and design a variant that bound Nile red but not DPH by exploiting only shape complementarity using RASSCCoL. The smaller 4-helix bundle required redesigning both ***a*** and ***d*** core positions of the ***abcdefg*** coiled-coil sequence repeats to generate binding sites (Supplementary Fig. 53). However, due to the increased radius and changes in core packing of the 6-helix barrel^20^, we reasoned that redesigning only ***a*** positions should accommodate the ligands (Supplementary Fig. 53). In addition, the larger scaffold allowed the amino-acid pallet to be expanded to include Phe, which the 4-helix bundle cannot accommodate. By restricting designs to just the ***a*** sites of three layers and a subset of hydrophobic residues (Gly, Ala, Val, Leu, Ile, and Phe), ≈1.3 million sequences passed the cavity volume filter (Supplementary Table 1). Using active sampling with GBT, only 130,000 sequences were evaluated using AutoDock Vina, thus reducing the computation 10-fold (Supplementary Figs. 54 – 55).

Again, we tested just two of the resulting designs experimentally: one intended to bind both ligands equally well (sc-apCC-6-DPH-NR, labelled P for promiscuous), and another specific for Nile red (sc-apCC-6-NR, labelled S for specific) (Fig. 4a). Both proteins expressed well, were monomeric, α helical, and thermally stable (Supplementary Figs. 56 – 57). As predicted, we observed low-μM affinities for DPH and Nile red in the promiscuous binder (Fig. 4b, panel P). By contrast, the selective binder exhibited 170-fold tighter binding for Nile red compared to DPH, with a sub-µM dissociation constant for the former (Fig. 4b, panel S). Inspection of the docked AF2 models suggests that selectivity against DPH—which is ≈2 Å longer than Nile red—arises from subtle shortening of the binding pocket, causing steric clashes between DPH and residues at the extremes of the binding pocket (Fig. 4c).

**Fig. 4.**
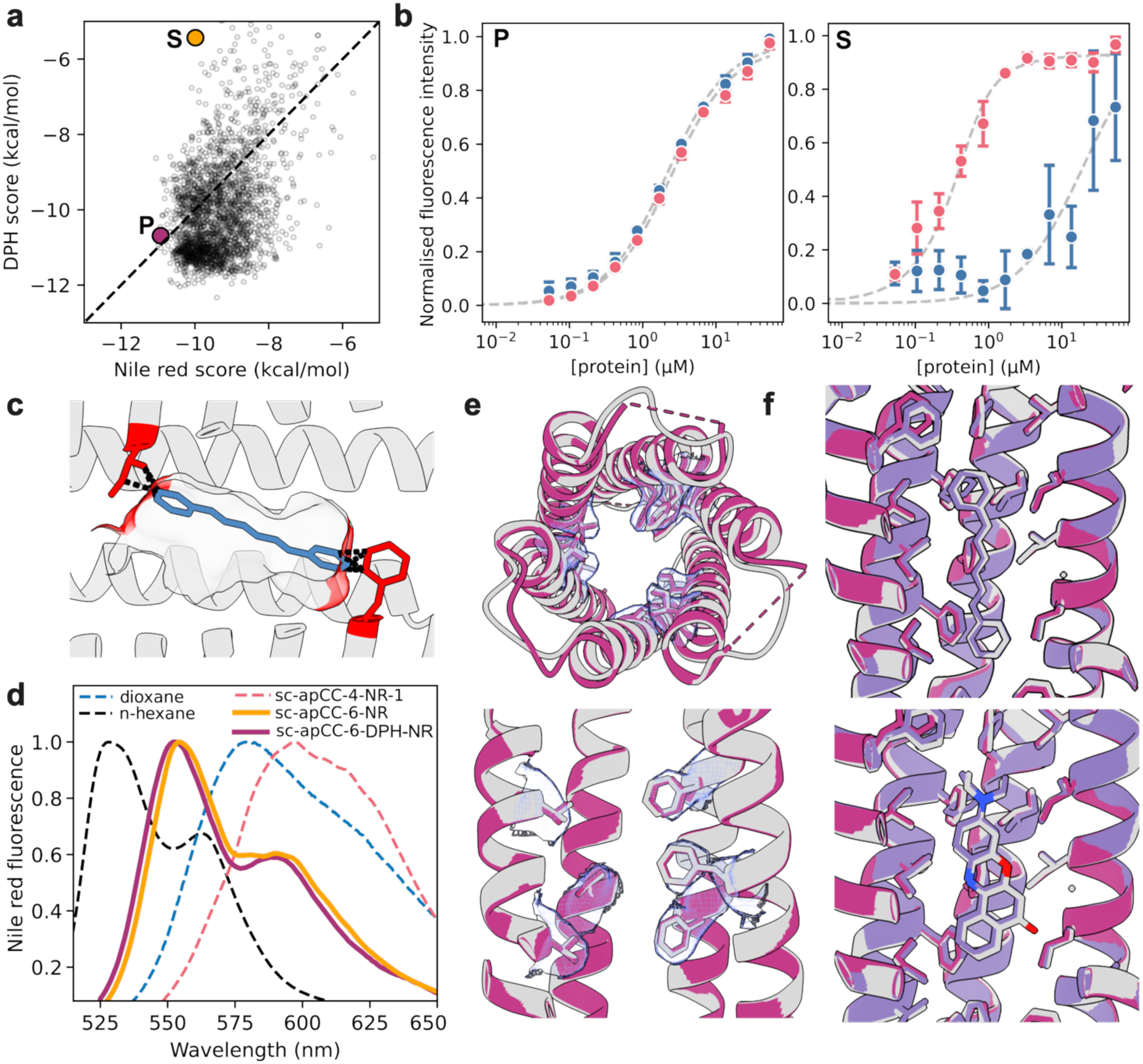
RASSCCoL-generated promiscuous and selective binders for DPH and Nile red. **a,** AutoDock Vina scores for DPH and Nile red docked into AF2 models for the top 1,000 generated designs. The dashed line is the 1:1 diagonal, and the Nile red binders are labelled P (promiscuous) and S (selective). **b,** Saturation binding curves and fits for Nile red and DPH to the proteins. **Left**, promiscuous binder, K_D(DPH)_ = 1.9 ± 0.2 μM, K_D(Nile red)_ = 2.2 ± 0.2 μM. **Right**, the specific binder, K_D(DPH)_ > 20 μM, K_D(Nile red)_ = 0.11 ± 0.04 μM. **c,** AF2 model for the specific binder with DPH docked. Selectivity against DPH is achieved by Leu and Phe residues (red) that shorten the binding site. Severe clashes between DPH and the side chains (overlap > 1 Å) are shown in black. **d,** Normalised emission spectrum for Nile red after excitation at 500 nm in aqueous solution with the specific and promiscuous 6-helix binders, compared to the 4-helix Nile red binder and organic solvents. **e,** Structural alignment between the AlphaFold2 model (grey) and the experimental X-ray crystal structure of the promiscuous binder P (sc-apCC-6-DPH-NR) at 3.2 Å resolution (Ca RMSD_AF2_ = 0.294 Å, PDB id 9R1O). NCS-averaged electron density for the designed residues is shown at 1σ level in blue. **f,** Boltz2 poses of DPH and Nile red in the promiscuous binder P (grey), overlaid with the design target (RASSCCoL repacked and redocked model; pale purple) and the ligand-free crystal structure (magenta). Oxygen atoms shown in red and nitrogen atoms in blue. Binding conditions: 0.5 μM ligand, 0 – 60 μM protein, PBS, 10% v/v MeCN, pH 7.4. Data are the mean of n=3 independent repeats, error bars represent the standard deviation from the mean. Source data are provided as a Source Data file.

Nile red is an environment-sensitive probe^34^. This is apparent in changes in its emission spectrum upon binding to the P and S designs (Fig. 4d): *i.e.*, the resolved vibronic bands similar to those seen in n-hexane; and the blue-shifted fluorescence peak relative to that observed in dioxane and the more-solvent-exposed binding site of sc-apCC-4-NR-1^24,35^. This infers binding to tight hydrophobic pockets of the P and S designs. Finally, we solved a structure of P without ligand to 3.2 Å. This matched the AF2 model with Cα-RMSD of 0.3 Å (Fig. 4e, PDB id 9R1O). Despite the medium resolution, electron density for the designed residues— particularly the Phe side chains—was well-resolved (Fig. 4e). Protein:ligand complex modelling with Boltz2 returned ligand poses that closely aligned with the ligand-free binding pocket observed in the X-ray crystal structure (Fig. 4f).

### Binders function *in vitro* and in cells

Building beyond single-domain, small molecule-binding proteins will require generating binding sites (or binding modules) that are orthogonal and can be co-assembled in modular manner into more-complex *de novo* frameworks. To explore this, we used two of the designed 4-helix bundles that bind DPH and Nile red selectively (sc-apCC-4-DPH-1 and sc-apCC4-NR-1, Supplementary Table 5, Supplementary Fig. 22) to construct a two-domain *de novo* protein. For this, the domains were fused *via* a rigid, read-through helical linker (Fig. 5a). Since DPH and Nile red are a known Förster resonance energy transfer (FRET) pair^24,36^ (Supplementary Fig. 58), we designed this construct to position the two dyes strategically at the Förster distance of ≈50 Å. The resulting protein was expressed in *E. coli*, and was monomeric, α-helical and thermally stable (Supplementary Figs. 12 and 15). Small-angle X-ray scattering (SAXS) indicated an extended overall conformation consistent with the model (Fig. 5a,b, Supplementary Table 10). The protein retained its affinities for both ligands (Fig. 5c and Supplementary Fig. 59). Moreover, with both ligands present, selective excitation of DPH resulted in both DPH and Nile red fluorescence (Fig. 5d and Supplementary Figs. 60 – 62). The latter must arise from FRET, indicating a population of the protein with both DPH and Nile red bound. The FRET lifetime of 2 ± 0.2 ns gave a predicted distance of 51 ± 1 Å, consistent with the computational design model and MD simulations of it (Supplementary Fig. 64). This experimentally confirmed the design of the dual binder (Fig. 5e).

**Fig. 5:**
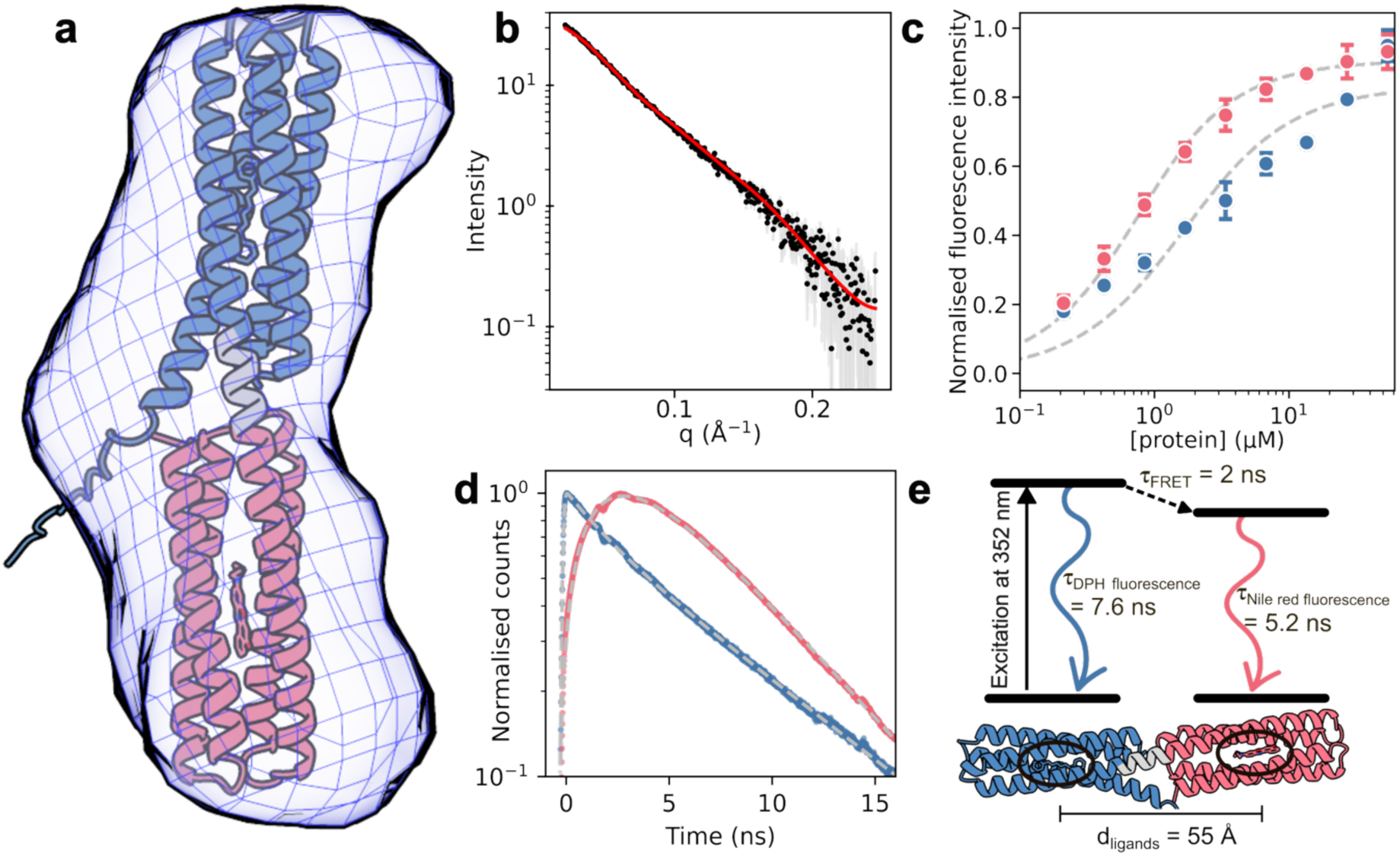
RASSCCoL-derived double-binder that transfers energy from DPH (blue) to Nile red (red). **a,** AF2 model of the double-binder that combines sc-apCC-4-DPH-1 and sc-apCC-4-NR-1, superposed to the averaged *ab initio* model from small angle X-ray scattering data. DPH and Nile red are docked with AutoDock Vina. **b,** Small angle X-ray scattering data and FoXS ^37^ fit to the AF2 model (red, χ2 = 1.44). **c,** Saturation binding curves and fits: blue, K_D(DPH)_ = 1.3 ± 0.4 μM; red, K_D(Nile red)_ = 0.53 ± 0.05 μM. **d,** Fluorescence lifetime decay measurements for DPH (blue, < 450 nm) and Nile red (red, >600 nm) after selective excitation of DPH at 352 nm. The delayed rise in Nile red emission over the first few nanoseconds originates from Förster resonance energy transfer between DPH and Nile red. In contrast, control measurements that excited only Nile red (Supplementary Fig. 62) showed no delayed emission. Bi-exponential fit of the data returned t_FRET_ = 2.0 ± 0.2 ns and population size of 0.23, yielding an inter-ligand distance of 51 ± 1 Å. The expected distance from MD simulations of the model from panel **a** was 55 ± 1 Å, see Supplementary Fig. 64 and Supplementary Table 11. **e,** Kinetic model for energy transfer between DPH and Nile red, derived from panel **d**, and direct excitation of DPH and Nile red in isolation (Supplementary Figs. 61 and 62). TCSPC conditions: 20 μM of each ligand, 30 μM protein, PBS, 10% v/v MeCN, pH 7.4; the non-FRET population arises from partial co-occupancy of the protein by both ligands, which results from using sub-stoichiometric ligand concentrations to minimise scattering. Binding conditions: 0.5 μM ligand, 0 – 60 μM protein in PBS, 10% v/v MeCN, pH 7.4. Binding data are the mean of n=3 independent repeats, error bars represent the standard deviation from the mean. SAXS conditions: 5 mg/mL protein in PBS, pH 7.4; black points are the mean from n=10 successive measurements, grey error bars indicate the standard deviation from the mean. Source data are provided as a Source Data file.

Finally, we explored the potential for using the *de novo* synthetic chromophore-binding proteins to target and visualise subcellular structures in mammalian cells. There are advantages of using *de novo* fluorescent proteins for such applications. For instance, at ≈130 residues and ≈15 kDa, sc-apCC-4 variants are approximately half the size of commonly used GFP variants^38^ and on par with some of the smallest FPs, such as: iLOV (13 kDa)^39^; SNAP-tag FP mimics (20 kDa)^40^; *de novo* β barrels (14 kDa)^41^; and transmembrane 4-helix bundles (21 kDa)^42^. Moreover, bright fluorophore encapsulation within proteins generally leads to improved and tuneable spectral properties compared to conventional FPs. To examine this, we found that Nile red bound to sc-apCC-4-SN38-1, had a high fluorescence quantum yield and thus brightness (e_552_ = 52 400 cm^-1^ M^-1^; Φ = 0.9), and a long fluorescence lifetime (τ = 4.8 ns) (Supplementary Table 12 and Supplementary Fig. 65). This is brighter and longer lived than conventional fluorescent proteins in the same spectral region, such as mCherry (e_587_ = 72 000 cm^-1^ M^-1^; Φ = 0.2; τ = 1.4 ns) ^43^. Furthermore, emission wavelengths can be tuned toward the near-infrared (NIR) by substituting Nile red for derivatives with similarly high quantum yields, brightness and long fluorescence lifetimes; e.g., Nile blue (e_627_ = 34 400 cm^-1^ M^-1^; Φ = 0.74; τ = 4.2 ns) (Supplementary Table 12 and Supplementary Figs. 66 – 68)^44^.

On this basis, we tested several of our *de novo* protein modules in HeLa cells as fusions to mEmerald, and with or without additional peptide tags for directing the constructs to different subcellular regions (Supplementary Table 13, Supplementary Fig. 69). For the main text these experiments are illustrated for constructs based on sc-apCC-4 and sc-apCC-4-SN38-1 and with Lifeact peptide tags for targeting actin filaments (Fig. 6a). The sc-apCC-4 constructs are non-binding controls, whereas its SN38 variant binds both Nile red and Nile blue; K_D,Nile Red_ ≈ 0.2 µM and K_D_,_Nile blue_ ≈ 0.4 μM (Supplementary Table 5, Supplementary Figs. 22, 66, 67).

**Fig 6:**
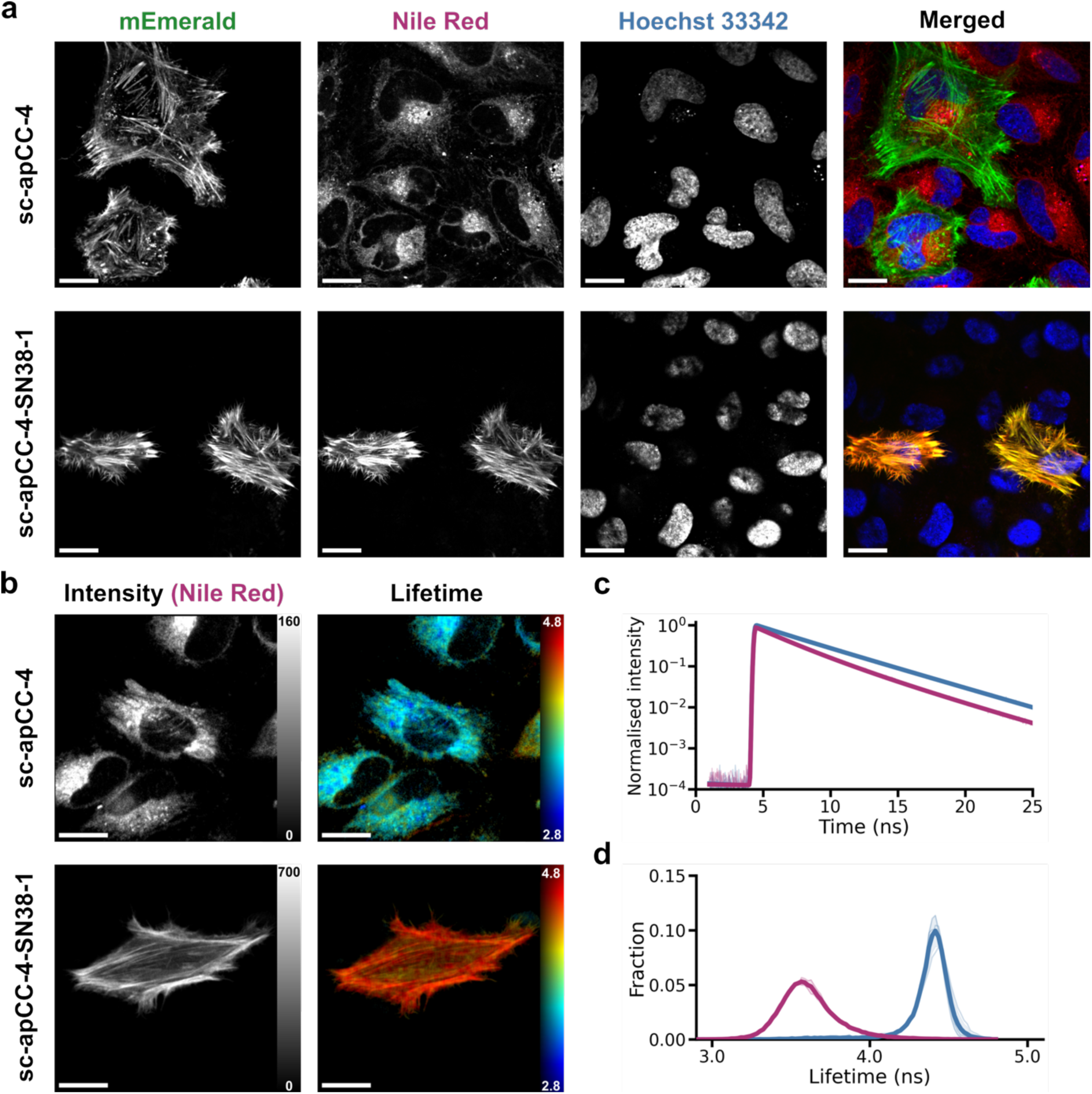
*De novo* designed fluorescent proteins for subcellular imaging, illustrated by targeting actin filaments. **a**, Fluorescence confocal microphotographs of live HeLa cells transiently expressing fusions for either the non-binding parent sc-apCC-4 control (top row) and sc-apCC-4-SN38-1 (bottom row). Both constructs have *N*-terminal Lifeact and *C-*terminal mEmerald modules. Cells were stained by adding 0.1 µM Nile red and Hoechst 33342 to the cell media 30 minutes before imaging. Merge (right-hand column): mEmerald (green), NR (red), and Hoechst 33342 (blue). The images are representative of n=3 independent experiments, with 3–5 fields of view acquired per condition in each experiment. **b**, Fluorescence lifetime images of Nile red in live HeLa prepared as above. Fluorescence intensity (left) scaled up to 160 units for sc-apCC-4 and 700 units for sc-apCC-4-SN38-1, and intensity-weighted mean lifetime (right) scaled from 2.8 to 4.8 ns. The images are representative of n=1 independent experiment per condition, with 3 fields of view acquired per condition. **c&d**, Fluorescence decays measured for Nile red (**c**), and distributions of Nile red fluorescence lifetimes (**d**) across pixels above threshold for sc-apCC-4 (deep pink lines) and sc-apCC4-SN38-1 (blue lines). For panels **a** & **b**, all scalebars = 20 μm. Source data are provided as a Source Data file.

For the control, we fused the parent sc-apCC-4 to the *N* terminus of mEmerald variant of the green fluorescent protein and transformed the construct into HeLa cells (Supplementary Table 13, Supplementary Fig. 69). The transiently expressed fusion protein was dispersed throughout the cell. Next, we added short, *N-*terminal peptide sequences to the fusion to target: the nucleus with a nuclear-localization signal (NLS)^45^; the plasma membrane with a Fyn-tag^46^; and the actin network with Lifeact^47^. In all cases, the mEmerald signal was redistributed and located as expected (Supplementary Fig. 69). These experiments were performed with 0.1 µM Nile red added to the cell media 30 minutes before imaging. However, with these non-Nile-red-binding controls, none of the images showed any coincidence of the mEmerald and Nile red signals. By contrast, with sc-apCC-4 substituted by sc-apCC-4-NR-1 or sc-apCC-4-SN38-1, the mEmerald and Nile red signals were coincident as evident in the merged images (Supplementary Figs. 69). These indicate that the dye had been taken up by cells and sequestered by the designed Nile red-binding components of the fusion proteins. Interestingly, the images were sharper and clearer with the sc-apCC-4-SN38-1 design (shown in more detail in Supplementary Fig. 70). We attribute this to its sub-µM affinity for Nile red, which is 10-fold tighter than for sc-apCC-4-NR-1 (Supplementary Table 5 and Supplementary Fig. 22).

As the fluorescence lifetime of Nile red is sensitive to the polarity of its environment^48,49^, we used fluorescence lifetime imaging microscopy (FLIM) to confirm Nile red binding by sc-apCC-4-SN38-1 in cells. For all tagged sc-apCC-4-SN38-1:mEmerald constructs, Nile Red fluorescence accumulated at the expected cell structures, and intensity-weighted FLIM images resolved these compartments due to the long-lifetime Nile Red signal at these sites (≈4.4 – 4.5 ns) (Supplementary Fig. 71). Again, this is illustrated here in the main text for constructs with *N*-terminal Lifeact tags (Fig. 6b-d). These values closely matched the lifetime of Nile red bound to sc-apCC-4-SN38-1 *in vitro* (4.8 ± 0.2 ns, Supplementary Table 12, Supplementary Fig. 65). By contrast, for the control constructs with the non-binding parent, sc-apCC-4, the fluorescence lifetimes (≈3.6 – 3.7 ns) were consistently and significantly reduced (Fig. 6b-d, and Supplementary Fig. 71). Furthermore, the fluorescence of Nile red in the cells expressing the binding proteins exhibited a uniform mono-exponential decay, indicating a well-defined environment in the designed cavity (Fig. 6c, Supplementary Fig. 72). In contrast, in cells expressing the parent sc-apCC-4, Nile red fluorescence decay was more complex characteristic of multiple environments and consistent with Nile red in non-transfected HeLa cells (Supplementary Fig. 73).

Finally, to demonstrate that sc-apCC-4 alone is sufficient as a *de novo* fluorescent protein chassis, we repeated the experiments without the mEmerald reporter. These worked similarly to the fusion constructs showing robust staining of targeted subcellular structures (Supplementary Fig. 74).

## Discussion

Designing proteins that bind small molecules tightly and specifically is a key challenge for protein design and synthetic biology, with potential applications including bioimaging, *in vivo* sensing, catalysis and therapeutics^12^. Here we present RASSCCoL, a fast, efficient, and intuitive computational strategy for introducing small molecule-binding sites into *de novo* designed proteins. This works by replacing side chains of the *de novo* scaffold to create cavities that are isosteric and chemically complementary with the target ligand. The calculations are vastly accelerated by: i) using a volume filter to enrich for sequences most likely to accommodate the ligand; ii) rapid modelling with FASPR^26^ and AutoDock^17^; (iii) evaluation of the models using statistical and physical metrics; and iv), where needed for large targets, an active-learning procedure to reduce the number of explicit modelling steps. To put this approach into practice, we leverage well-understood, mutable, and thermostable *de novo* proteins as the scaffolds; namely, a growing family of single-chain α-helical bundles and barrels^19,20^. This streamlines the design process even further by bypassing backbone optimization and focusing on sequence design and refinement. Although only a subset of these scaffolds was explored in the present study, the framework is readily extendable to a broader range of architectures within this family.

Specifically, we have designed low-to-sub-micromolar protein binders for two hydrophobic fluorophores, DPH and Nile red; for lumiflavin, a proxy for flavin-based cofactors; and for two cytotoxic compounds, arcyriaflavin A and SN38. The last three feature polar functional groups that RASSCCoL accommodates by placing complementary polar side chains in the designed binding sites. In each case, successful designs are achieved by testing just 1 – 4 expressed genes in *E. coli*. The approach scales well to larger scaffolds, or with additional design constraints such as ligand selectivity, which we also demonstrate experimentally. To test the modularity of some of the resulting designs, we combine two different binding modules for DPH and Nile red (a known FRET pair) in a two-domain protein that holds the chromophores in a specified orientation and at a set distance to enable energy transfer between the two sites. Finally, we demonstrate the potential of our designs as small (≈15 kDa), stable, mutable, and adaptable fluorescent protein tags for visualizing subcellular structures and processes in mammalian cells.

Together with advances by others (Supplementary Fig. 75)^9,10,42^, our findings reinforce how the use of straightforward, well-understood, and fully characterised *de novo* scaffolds can accelerate functional protein design for small-molecule binding. In this way, RASSCCoL offers a robust and interpretable platform for creating next-generation protein-based tools and modules for broader applications in cell and synthetic biology, diagnostics, and therapeutics.

## Methods

### General

All solvents, chemicals, and reagents were purchased from commercial sources and used without further purification. Luria-Broth (LB), antibiotics, IPTG, and L-rhamnose were purchased from Thermo Fisher. All other chemicals were reagent grade and purchased from Sigma-Aldrich. Protein biophysical characterization were recorded in a PBS buffer (50 mM sodium phosphate, 150 mM NaCl, pH 7.4).

### Computational design

Van der Waals volumes for ligands, side chains, and cavities were calculated from Bondi radii^25^. Binding-site sequences were generated and filtered for cavity volumes matching ligand size within a ± 30 % cutoff (see Supplementary Table 1 for all input parameters). For SN38, the closed lactone form representing the pharmacologically active tautomer was used during the computational design process, as only this form engages in canonical binding interactions in vivo^50^. Passing sequences were packed with FASPR^26^. Ligands were docked with AutoDock Vina^17^ using parameters optimised for speed (exhaustiveness of 1, box size of 1.5 × ligand length, spacing of 0.5 Å, and 5,000 evaluations). Sequences below a user-defined energy cutoff were refolded with AF2, see Supplementary Figs. 3 – 8. For Nile red, lumiflavin, arcyriaflavin A and SN38, only AF2 model 4 was used. Earlier DPH designs used all five AF2 models. Candidates for experimental validation were chosen based on AutoDock Vina^17^ scores after redocking with default parameters (exhaustiveness of 16, spacing of 0.375 Å, and 10,000 evaluations). Throughout the pipeline, no backbone or side-chain minimisation or relaxation was performed.

For design refinement, 1000 RASSCCoL models with the lowest AutoDock Vina scores after repacking were redesigned using LigandMPNN (default settings; 5 sequences per input) and subsequent protein:ligand complex prediction with Boltz2. For DPH and Nile Red, 270 and 600 sequences were used, respectively, reflecting the smaller sequence space of RASSCCoL binders for these ligands. The top 1000 sequences ranked by ipTM were then subjected to a second round of redesign and refolding. After two rounds, Boltz2 confidence metrics and the binding-site sequences had converged (Supplementary Figs. 24 – 28).

For active sampling benchmarks, sequences were one-hot encoded for linear, ridge, and MCP models, and integer encoded for RF and GBT models. Models were initialized with 50,000 random sequences and iteratively updated in steps of 10,000. Uncertainty was estimated via bootstrap ensembles (linear models, 4×, 80% data) or tree subsampling (RF/GBT). Sampling used a min–max normalized acquisition function (μ + σ). The validation set matched the initialization size, and the test set comprised 10% of the data. In pool-based learning, 100× the step size was randomly sampled per iteration. Early stopping based on Spearman correlation was used to control the number of trees in RF/GBT. Ranking performance was evaluated using recall@1% and recall@5%, defined as recovery of the true top k% (lowest AutoDock Vina scores) within the top k% of model predictions. Linear models from scikit-learn were implemented with default parameters; tree models from LightGBM used 80% row and 90% feature subsampling, 100 leaves, and a decaying learning rate (0.1 to 0.01).

For the production run on sc-apCC-6 dataset (1.3 million sequences), a GBT model was initialized on 20,000 sequences and trained via pool-based active learning (step size 1,000) until recall@1% plateaued on a 20,000-sequence validation set (40 iterations). The trained model was then used to predict AutoDock Vina scores for the remaining sequences; 50,000 best predictions were then taken further.

Additional details of RASSCCoL pipeline are available on Woolfson Lab GitHub (https://github.com/woolfson-group/RASSCCoL).

### Molecular dynamics simulations

OpenMM 7.7 was used for running MD simulations^51^. Simulations were run with the Amber ff14SB force field, Langevin Middle integrator with an integration step of 4 fs, 1.5 au H-mass repartitioning and collision rate of 1 ps^-1^.

For sc-apCC-4-DPH-1-sc-apCC-4-NR-1, the system (Supplementary Table 14) was solvated with tip3p waters and 100 mM NaCl in a rhombic dodecahedron with a padding of 1 nm, particle mesh Ewald with a cut-off of 1 nm was used for long-range electrostatics, and the pressure was kept constant at 1.01325 bar with a MonteCarlo barostat. The AlphaFold2 model with the ligands docked was initialised at 10 K with 5 kcal/mol restraints on the protein backbone and ligand heavy atoms and minimised. The system was then heated to 300 K over 600 ps. Heavy atom restraints were relaxed over 3.5 ns and the unrestrained system was further equilibrated for 10 ns. A production run was then performed for 100 ns. Periodic boundary conditions were removed using GROMACS trajconv^52^, trajectories were analysed using MDAnalysis^53^.

Short MD simulations to evaluate SN38 binding were run with implicit water (obc2) using a repacked 4-helix bundle (PDB id 8A3K) with SN38 docked within the binding pocket. The structure was initialised at 0 K, minimised, heated to 400 K over 400 ps, and then equilibrated for 1.6 ns. A production run was performed for 5 ns. Trajectories were analysed using MDAnalysis^53^, and ligand binding affinities were analysed using AutoDock Vina. The simulations were run with and without 20 cal mol^-1^ Å^-2^ backbone heavy atom restraints to minimize protein unfolding at elevated temperatures.

### Protein expression and purification

Codon-optimized DNA encoding the protein sequences was purchased from IDT or GeneArt, cloned into pET28a vectors using NdeI and XhoI, transformed, and expressed in *E. coli* Lemo21(DE3) (New England Biolabs). DNA sequences are provided in Supplementary Data. Flasks containing 1 L of Miller’s Luria Broth–kanamycin–chloramphenicol and 0.5 mM L-rhamnose were inoculated with 5 ml of overnight cultures and incubated to an optical density at 600 nm of ∼0.6 at 37 °C with 200 r.p.m. shaking. Expression was induced with 0.5 mM isopropyl-β-D-thiogalactoside, and cultures were incubated at 37 °C overnight with 200 r.p.m. shaking. Following expression, cultures were pelleted and resuspended in 20 ml lysis buffer (50 mM Tris, pH 7.4, 500 mM NaCl, 30 mM imidazole, 1 mg ml^-1^ lysozyme) for 30 min at 37 °C. Resuspended pellets were sonicated using a Biologics Model 3000 Ultrasonic homogenizer with settings at 50% power and 90% pulser (1 pulse per second) for 5 min and then clarified at 25,500 *g* for 30 min. The clarified lysate was heat shocked at 75 °C for 10 min and then cooled on ice for 10 min before reclarifying at 25,500 *g* for 10 min. The expressed proteins were first purified with Ni-based immobilised metal affinity chromatography at room temperature. Filtered lysate was loaded onto an ÄKTAprime plus (GE, PrimeView 5.31) equipped with a HisTrap HP 5-ml column (Cytiva). His-tagged proteins were eluted using a single step gradient from 0 to 55% buffer B (buffer A consisted of 50 mM Tris, 500 mM NaCl and 30 mM imidazole at pH 7.4; buffer B consisted of 50 mM Tris, 500 mM NaCl and 300 mM imidazole at pH 7.4). Fractions were combined and further purified by SEC using a HiLoad 16/600 Superdex 200-pg or 75-pg size exclusion column (Cytiva) equilibrated in buffer containing 50 mM sodium phosphate and 150 mM NaCl (pH 7.4) at room temperature. Eluted fractions were pooled, concentrated and separated using SDS–PAGE to confirm protein identities.

### Circular dichroism spectroscopy

Circular dichroism data were collected on a JASCO J-810 or J-815 spectropolarimeter fitted with a Peltier temperature controller in the far UV region. Spectra Manager (1.55) was used for data collection. Protein samples were prepared as 5-15 μM solutions in 50 mM sodium phosphate and 150 mM NaCl (pH 7.4) at 5 °C. Data were collected in a 1-mm quartz cuvette between wavelengths of 190 nm and 260 nm with the instrument set as follows: band width, 1 nm; data pitch, 1 nm; scanning speed, 100 nm min^-1^; response time, 1 s. Each circular dichroism spectrum was obtained by averaging eight scans and subtracting the background signal of the buffer and cuvette. For thermal response experiments, the circular dichroism signal at a 222 nm wavelength was monitored over the temperature range 5 – 95 °C at a ramping rate of 60 °C per hour with the same settings and protein concentrations given above. The spectra were converted from ellipticities (mdeg) to mean residue ellipticities (deg·cm^2^·dmol^-1^·res^-1^) by normalizing for concentration of peptide bonds and the cell path length using the equation:

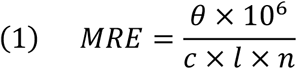

where the variable *θ* is the measured difference in absorbed circularly polarized light in millidegrees, *c* is the micromolar concentration of the compound, *l* is the path length of the cuvette in millimeters, and *n* is the number of amide bonds in the polypeptide.

### Analytical ultracentrifugation

AUC was performed on a Beckman Optima X-LA or X-LI analytical ultracentrifuge with an An-50-Ti or An-60-Ti rotor (Beckman-Coulter) equipped with ProteomeLab XL-A (5.5) software. Buffer densities, viscosities, and peptide and protein partial specific volumes (*v̄*) were calculated using SEDNTERP (http://rasmb.org/sednterp). For sedimentation velocity, protein samples were prepared in PBS at a 30 μM protein concentration and placed in a sedimentation velocity cell with a two-channel centerpiece and quartz windows. The samples were centrifuged at 40 k.r.p.m (129,000 × g at 7.20 cm) at 20 °C, with a total of 120 absorbance scans taken over a radial range of 5.8 – 7.3 cm at 5 min intervals. Data from a single run were fitted to a continuous *c*(*s*) distribution model using SEDFIT (v15.2b) at a 95% confidence level ^54^. Residuals for sedimentation velocity experiments are shown as a bitmap in which the grayscale shade indicates the difference between the fit and raw data (residuals, <−0.05 black and >0.05 white). Good fits are uniformly grey without major dark or light streaks.

### Ligand binding

Ligand-binding experiments were performed in triplicate. The total concentration of ligand was kept constant and below reported solubility limits in aqueous solution^55-58^: 0.5 μM DPH/Nile red or 1 μM arcyriaflavin A in 10% v/v MeCN; 1 μM lumiflavin/SN38 in 10% v/v DMSO. At these concentrations, neither solvent measurably affected protein folding or stability. In addition, binding was evaluated relative to the corresponding non-specific control (sc-apCC-4-Ala). Hence, we do not expect the solvent choice to introduce a meaningful confounding effect on the interpretation of the results. The concentration of the *de novo* protein design was varied from 0 – 54 μM. Fluorescence intensity and polarisation data were collected on a Clariostar plate reader (BMG Labtech) using the following excitation wavelength: λ_DPH_ = 352 nm, λ_Nile red_ = 520 nm, λ_arcyriaflavin A_ = 320 nm, λ_lumiflavin_ = 450 nm, λ_SN38_ = 350 nm. Fluorescence anisotropy was employed for ligands that did not exhibit significant changes in fluorescence intensity upon binding (e.g., lumiflavin), whereas intensity-based measurements were used when binding produced a measurable fluorescence change, providing improved sensitivity and signal-to-noise. Binding constants were extracted by fitting the data to the following equation:

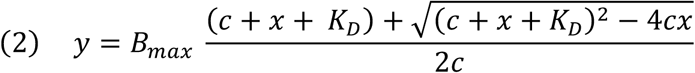

where c is the total concentration of the constant component (ligand), x is the concentration of variable component (protein), Bmax is the fluorescence intensity/polarisation signal when all the constant component is bound, and y is the fraction of bound component being monitored via fluorescence signal.

### Small-angle X-ray scattering

Data for sc-apCC-4-DPH-1-sc-apCC-4-NR were collected at The European Synchrotron Radiation Facility beamline BM29. Batch mode samples were prepared at 0.5 - 5 mg/mL in 50 mM sodium phosphate pH 7.4 and 150 mM NaCl. The His-tag was not removed as we observed non-specific cleavage at the domain linker after incubation with TEV protease.

Data for sc-apCC-4-DPH-2 were obtained at the Diamond Light Source beamline B21. The His-tag was removed using thrombin protease. Size-exclusion chromatography mode samples were prepared at 20 mg/mL in 20 mM Tris pH 8.0, 50 mM NaCl, 1% v/v glycerol. Shodex KW403 column was equilibrated in the same buffer at 4 °C.

For both designs, buffer subtraction and data merging was performed with ScÅtter^59^. q_min_ was taken as the first point of the linear Guinier region, q_max_ was calculated using ShaNum through ATSAS interface^60^. MultiFoxS software (Sali Lab) was used to compare experimental scattering profiles to design models and assess quality of fit by calculating χ^2^ ^37,61^. Sc-apCC-4-DPH-1-sc-apCC-4-NR data were fit to a His-tagged AlphaFold2 model. Sc-apCC-4-DPH-2 data were fit to a dimer observed in the crystal structure (PDB id 9R1N) and a monomeric AlphaFold2 model.

### X-ray crystallography

Crystals were grown using a sitting-drop vapour-diffusion method. Proteins in 20 mM Tris pH 8.0, 50 mM NaCl were concentrated to 20 mg/mL. Commercially available sparse matrix screens were used (Morpheus®, JCSG-plus^TM^, Structure Screen 1 and 2, Pact Premier^TM^, ProPlex^TM^; Molecular Dimensions), and the drops were dispensed using a robot (Oryx8; Douglas Instruments). For each well of an MRC 2 drop plate, 0.3 μL of peptide or protein solution and 0.3 μL of reservoir solution in parallel with 0.4 μL of the peptide or protein solution and 0.2 μL of reservoir solution were mixed and the plate was incubated at 20 °C. Diffraction-quality crystals were grown through multiple rounds of seeding. Crystals generally formed within a month, and after looping were soaked in reservoir solution containing 25% glycerol as a cryoprotectant. Final crystallization conditions for are provided in Supplementary Table 8.

Diffraction data for the crystals were obtained at the Diamond Light Source (Didcot, UK) on beamlines I04 or I24. Sc-apCC-4-DPH-2, sc-apCC-4-NR-1, sc-apCC-4-SN38-1 and sc-apCC-4-A7F-1 data were processed using the automated Xia2 pipeline^62^, which ports data through DIALS^63^ to POINTLESS and AIMLESS^64^ as implemented in the CCP4 suite^65^. Sc-apCC-6-DPH-NR and sc-apCC-4-LMF-1 were processed using the AUTOPROC pipeline, which use the same integrating and data reduction software in addition to STARANISO^66^. All structures were phased by molecular replacement using the AlphaFold2 model for PHASER^67^. Final structures were obtained after iterative rounds of model building with COOT^68^ and refinement with Phenix Refine^69^. Solvent-exposed atoms lacking map density were deleted. Data merging and refinement statistics for are provided in Supplementary Table 9.

For co-crystallisation trials, protein (50 μM) was incubated with ligand (100 μM) in 20 mM Tris (pH 8.0), 50 mM NaCl, 10% DMSO for 24 h prior to concentration to 20 mg/mL using centrifugal concentrators. Crystallisation screens were set up as for the apo protein. For soaking experiments, crystals were transferred to drops containing saturated ligand solution containing crystallization condition supplemented with 20% DMSO, and incubated for 1 min, 5 min, 10 min, 1 h, 2 h, and 24 h prior to harvesting. Co-crystallisation produced crystals under the same conditions as the apo protein (Supplementary Table 8); however, no ligand electron density was observed. Soaking experiments did not yield diffraction-quality crystals.

### Fluorescence lifetime measurements

Time-correlated single photon counting (TCSPC) data were acquired using a home-built TCSPC apparatus.^70^ The output of a tunable high-power ultrafast oscillator (Chameleon Ultra II, 3.7 W, 80 MHz, Coherent) was frequency doubled to generate the required 352 nm pulses for DPH excitation. For Nile red and Nile blue, the light was first frequently shifted to 1120 nm with an optical parametric oscillator (Chameleon Compact, APE) and then frequency doubled to generate 560 nm excitation pulses. To avoid re-excitation of samples, the repetition rate was reduced to 6.67 MHz using a pulse picker (cavity dumper, APE). The diffracted output was spatially filtered with a pinhole to remove residual zeroth order diffraction, and the resulting light was focused into a sample (1 cm path length cell). Fluorescence was collected at 90° relative to excitation using an infinity-corrected microscope objective (4×/0.2 NA Plan Apochromat, Nikon) and through a series of achromatic lenses collimated and focused onto an avalanche photodiode detector (ID100-50-ULN, IDQ). Nile red and Nile blue fluorescence was collected with a >600 nm filter, DPH with a 450 nm bandpass filter. Photon count arrival times were recorded with a time-to-digital converter (Time Tagger 20, Swabian Instruments) and accumulated in 10 ps bins.

The instrument response function (IRF) was 188 ps, as determined by recording laser scatter from solvent in a 1 cm path length cuvette. All data were collected at room temperature (20 °C) using the magic angle condition.

### Calculation of FRET distance between DPH and Nile red in sc-apCC-4-DPH-1-sc-apCC-4-NR-1

Lifetime decays after the direct excitation of DPH and Nile red were modelled as a mono- or bi-exponential decay convolved with a Gaussian IRF function.

The energy transfer from the donor (DPH) was modelled as a bi-exponential decay accounting for the population of DPH (A_2_) that are not involved in FRET:

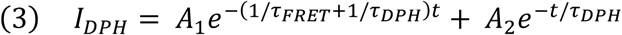

The energy transfer to the acceptor (Nile red) was modelled as a bi-exponential function where B_1_ is a negative amplitude reflecting the rise of acceptor fluorescence due to FRET:

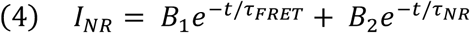

These were fitted to DPH and Nile red channels simultaneously while floating the exponential pre-factors (*A*_1_, *A*_2_, *B*_1_, *B*_2_) and *k*_FRET_. Average *k*_DPH_ and *k*_NR_ were kept fixed based on control measurements of the dyes alone within the protein. For readability, the equations shown above exclude the convolution terms required to model the instrument response function.

Assuming 1/τ_FRET_ = *k*_FRET_, the Förster distance and the error were calculated as:

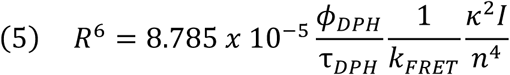

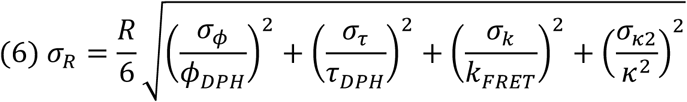

where Φ_DPH_ and *τ*_DPH_ are the fluorescence quantum yield and the lifetime of DPH inside the protein, *κ* is the orientation factor between the dipoles, *I* is the Förster spectral overlap integral of DPH fluorescence and Nile red absorption within the αHB, and *n* is the refractive index of the buffer (1.33). *κ*^2^ was derived from the molecular dynamics simulations. DPH fluorescence quantum yields within the protein were estimated by reference quantum yield measurements of DPH in acetonitrile (Φ_MeCN_ = 0.17) ^24^. The Förster spectral overlap integral was obtained using software freely available at FluorTools.com. The data were collected using 20 μM of each ligand, 30 μM protein, PBS, 10% v/v MeCN, pH 7.4.

### Nile red and Nile blue fluorescence quantum yield measurements in sc-apCC-4-SN38-1

Absolute fluorescence quantum yields were measured using an integrating sphere on a FS5 spectrofluorometer (Edinburgh Instruments) with λ_ex =_ 510 nm, λ_em_ = 485 – 870 nm for Nile red and λ_ex =_ 530 nm, λ_em_ = 510 – 870 nm for Nile blue. The data were collected at 2 μM of each ligand, 50 μM protein, PBS, 10% v/v MeCN, pH 7.4. The quantum yield for Nile blue was corrected for self-absorption. For Nile red, the self-absorption was less than the error.

### Transfection and imaging of HeLa cells

Codon-optimized linear DNA fragments for the designer constructs were synthesized by GeneArt or IDT and then subcloned in pTwist CMV vector (Twist Bioscience) for expression in HeLa cells. To make fusions of binder proteins with mEmerald and clone them in the target vector containing, PCR reactions were carried out using Q5 High Fidelity Hot Start DNA Polymerase (New England Biolabs (NEB)), following manufacturers’ instructions. PCR products were purified using Monarch® PCR & DNA Cleanup Kit (NEB). DNA and primer sequences are provided in Supplementary Data. Restriction digest reactions were carried out using NEB restriction enzymes at 37 °C for 1 h. Ligation reactions were carried out using T4 DNA Ligase and Rapid ligation buffer (Promega) for 1-2 h at 16 °C. Transformation was carried out in competent (subcloning efficiency) *Escherichia coli* cells (NEB® 5-alpha from NEB or DH5α from Invitrogen) for 30 min on ice, followed by heat shocking (30 s, 42 °C), and a further 5 min on ice. Cells were recovered by incubating in LB media at 37 °C for 1 h and then plated on LB-agar plates containing the ampicillin.

Plasmid DNA samples for transfection were purified using Plasmid Midi Kit according to the manufacturer’s instructions (QIAGEN) and verified by DNA sequencing using Rapid Sequencing Service provided by Source Bioscience.

HeLa cells (Cat. No. 93021013, ECACC, UK Health Security Agency) were maintained in high glucose Dulbecco’s Modified Eagle’s Medium with 10% (v/v) foetal calf serum (Sigma-Aldrich) and 5% penicillin/streptomycin (PAA) (herein referred to as DMEM) without phenol red at 37 °C and 5% CO_2_. For transfection, cells were seeded on tissue culture treated 35 mm CELLview cell culture dishes with 10 mm glass bottom (Greiner) at a density of 1x10^5^ cells per well and incubated at 37 °C and 5% CO2 for 16 h prior to transfection. Cells were transfected with 0.8 μg of the indicated plasmid DNA using Effectene transfection reagent according to the manufacturer’s instructions (QIAGEN). After transfection cells were incubated at 37 °C, 5% CO_2_ for 6 hours, washed with fresh DMEM and incubated further for 18 hours. Before imaging, cells were washed with PBS, stained with Hoechst 33342 (1 μg/mL) and Nile Red (0.1 μM) at 37 °C for 30 minutes, washed again, and then the medium was replaced for fresh DMEM.

Confocal images were collected using an Olympus IXplore SpinSR system with a 60x objective lens at 37 °C using the following imaging channels (λ_ex_-max λ_em_/width λ_em_): 405-447/60, 488-525/50, and 561-617/73 nm. Figures were assembled using the Fiji distribution of ImageJ2^71^.

FLIM was performed on a Leica SP8X laser-scanning confocal microscope equipped with a pulsed white light laser (Leica Microsystems). The laser was tuned to 561 nm to provide excitation of the Nile red, and a notch filter was used centred at 561 nm to minimise reflected light. Images were acquired with a 63x/1.2NA water immersion objective lens and fluorescence emission was collected between 590–750 nm. Images were acquired with 512 x 512 pixels with an additional zoom of 2x, yielding a pixel size of 180 nm. Laser power was adjusted to ensure photon counts avoid pulse pile-up artefact. Cells were maintained at 37 °C. A global-fitting algorithm for multi-exponential models was applied to analyse all pixels above an applied intensity threshold. Data was spatially binned 2x2 and fitted with a 2-exponential model^72^.

## Data availability

The coordinate and structure factor files for sc-apCC-4-DPH-2, sc-apCC-4-NR-1, sc-apCC-4-LMF-1, sc-apCC-4-A7F-1, sc-apCC-4-SN38-1 and sc-apCC-6-DPH-NR have been deposited in the PDB with accession codes 9R1N, 9R1J, 9R1M, 9R1K, 9R1L, and 9R1O, respectively.

Data from this study are openly available in Zenodo at DOI: 10.5281/zenodo.15333152. Source data for graphs in Figs. 1–6 is provided with this paper.

## Code availability

The code to run the RASSCCoL protocol can be found in the Woolfson group GitHub (https://github.com/woolfson-group/RASSCCoL). Archived code is available at DOI: 10.5281/zenodo.22673109.

## Acknowledgments

T.A.A.O., D.N.W. and R.P. thank Olivia Hawkins and Camilla Gajo for assisting with the fluorescence lifetime measurements. We thank the Diamond Light Source for access to beamlines B21, I03 and I24 (proposals mx37593 and mx31440) and The European Synchrotron Radiation Facility for access to beamline BM29 (proposal mx2551). Finally, we thank the members of the Woolfson group and collaborators from the Manchester Institute of Biotechnology, particularly Dr Derren Heyes, and the University of Sheffield for many valuable discussions.

## Authors contributions

D.N.W., J.J.C., K.O., and R.P. conceived the project. J.J.C., K.O., and R.P. wrote and tested the RASSCCoL code. J.J.C. and K.O. designed and characterised the lumiflavin and SN38 binders. R.P. designed and characterised the DPH and Nile red binders. R.P. and K.O. designed the arcyriaflavin A binders. R.P. collected the X-ray diffraction data and solved the crystal structures. A.V.R. devised and performed cell experiments. D.A. performed FLIM experiments and analysed the FLIM data. J.J.C, K.O., R.P., A.V.R. and D.N.W. wrote the paper, and all authors contributed to its revision. J.J.M., G.J.L., N.S.S., and D.N.W. raised the funding.

## Supplementary information

Supplementary Tables 1 – 14, Supplementary Figs. 1 – 75 and references.

