## Supplementary Figures and Tables for "Rapid Assessment of Chemical Complementarity of Ligands for Protein Design" (from another version of this preprint: version 2, 2026-05-22)

### 1 Supplementary Tables

**Supplementary Table 1.** Computational summary of RASSCCoL. *Since DPH is symmetric, the layers were constrained to 2-fold symmetry. **Active sampling was used to sample 100 000 sequences out of 1 277 192. CPU times are calculated as the number of sequences multiplied by the average packing and docking times measured in the benchmark (Supplementary Figure 1), rounded to the nearest second. For hydrophobic DPH and Nile red, amino acid palette was selected to reflect hydrophobic amino acid preference at **a** or **d** positions within the heptad. For flavin and SN38, amino acid palette was selected to reflect presence/absence of hydrogen bond donor/acceptor within each ligand layer.

| Ligand | Scaff old | Layer positions | Layer volumes, Å$^3$ | Amino acid palette | Number of sequences | Single CPU thread time, s |
|---|---|---|---|---|---|---|
| DPH* | 8a3k | [9, 77, 51, 118, 80]<br>[44, 111, 16, 84,114] | 82 – 192<br>82 – 192 | ***a*** = [AVLIG]<br>***d*** = [AVIG] | 3258 | 1890 |
| Nile red | 8a3k | [16,44,111,84,118,5 1, 114,12,19] | 324 – 389 | ***a*** = [AVLIG]<br>***d*** = [AVIG] | 396 774 | 273 774 |
| Nile red | 8qad | [23,95,167,44,191, 116,16,88,160] | 259 – 389 | ***a*** = [AVLIGF] | 100 000** | 69 000 |
| Lumiflavin | 8a3k | [9,77,121]<br>[12,51,118]<br>16,84,114] | 46-86<br>124-164<br>98-138 | [GAVLISTN]<br>[GAVLISTN]<br>[GAVLI] | 1 173 314 | 746 423 |
| Arcyriaflavin A | 8a3k | [16,44,111,84,118,5 1, 114,12,19] | 310 - 387 | ***a*** = [AVLGTS]<br>***d*** = [AVG] | 332 685 | 242 860 |
| SN38 | 8a3k | [12,51,118]<br>[16,84,114]<br>[19,44,87,111] | 145-165<br>145-165<br>180-200 | [GAVLISTN]<br>[GAVLISTN]<br>[GAVLISTN] | 51 262 | 38 447 |

**Supplementary Table 2.** AlphaFold2 metrics for all RASSCCoL proteins in this study. *DPH binders were designed with an earlier iteration of the protocol that used all 5 AlphaFold2 models and the default seed (“000”).

| Name | pLDDT | pTM | bb RMSD to parent, Å |
|---|---|---|---|
| sc-apCC-4-Gly | 89.8 | 0.8 | 1.1 |
| sc-apCC-4-Ala | 88.8 | 0.8 | 9.9 |
| sc-apCC-6-Leu | 95.2 | 0.9 | 0.5 |
| sc-apCC-4-DPH-1 | 91.2 | 0.7 | 0.5 |
| sc-apCC-4-DPH-2 | 90.5 | 0.8 | 10.2 |
| sc-apCC-4-NR-1 | 92.9 | 0.9 | 0.8 |
| sc-apCC-4-NR-2 | 92.1 | 0.8 | 0.9 |
| sc-apCC-4-LMF-1 | 92.4 | 0.8 | 0.4 |
| sc-apCC-4-LMF-2 | 92.6 | 0.9 | 1.1 |
| sc-apCC-4-LMF-3 | 92.6 | 0.9 | 0.8 |
| sc-apCC-4-SN38-1 | 92.2 | 0.9 | 0.9 |
| sc-apCC-4-SN38-2 | 92.5 | 0.9 | 0.9 |
| sc-apCC-4-SN38-3 | 92.0 | 0.8 | 0.9 |
| sc-apCC-4-SN38-4 | 92.4 | 0.8 | 0.9 |
| sc-apCC-4-A7F-1 | 93.3 | 0.9 | 0.6 |
| sc-apCC-4-A7F-2 | 93.2 | 0.9 | 0.7 |
| sc-apCC-6-DPH-NR | 95.5 | 0.9 | 0.4 |
| sc-apCC-6-NR | 95.6 | 0.9 | 0.4 |
| sc-apCC-4-DPH-1-sc-apCC-4-NR-1 | 91.22 | 0.7 | n/a |

**Supplementary Table 3.** Expressed RASSCCoL protein sequences for *in vitro* characterisation in this manuscript.

| Name | Scaffold | Sequence |
|---|---|---|
| sc-apCC-4-Gly | 8a3k | MGSSHHHHHHSSGLVPRGSHM<br>QLEEIAQQGEEGAKQGKKIAWQLKKIAQGEPSAQG<br>QLEEIAQQLEEGAKQGKKGAWQLKKIAQGPDSV<br>QLEEIAQQGEEGAKQGKKIAWQLKKIAQGGTSGG<br>QLEEIAQQLEEGAKQGKKGAWQLKKIAQ |
| sc-apCC-4-Ala | 8a3k | MGSSHHHHHHSSGLVPRGSHM<br>QLEEIAQQLEEAAKQAKKAAWQLKKIAQGEPSAQG<br>QLEEIAQQAEEAAKQAKKIAWQLKKIAQGPDSV<br>QLEEIAQQLEEAAKQAKKAAWQLKKIAQGGTSGG<br>QLEEIAQQAEEAAKQAKKIAWQLKKIAQ |
| sc-apCC-6-Leu | 8qad | MGSSHHHHHHSSGLVPRGSHM<br>LFEEIAQLLEEIAKLLKKIAWLLKKIAQGAQPLEM<br>LFEEIAQLLEEIAKLFKKIAWLLKKIAQTTETKKQGD<br>LFEEIAQLLEEIAKLLKKIAWLLKKIAQGYGDKRT<br>LFEEIAQLLEEIAKLFKKIAWLLKKIAQVAPTQRHRY<br>LFEEIAQLLEEIAKLLKKIAWLLKKIAQGTNSDSDLKS<br>LFEEIAQLLEEIAKLFKKIAWLLKKIAQG |
| sc-apCC-4-DPH-1 | 8a3k | MGSSHHHHHHSSGLVPRGSHM<br>QLEEIAQQGEEAAKQAKKIAWQLKKIAQGEPSAQG<br>QLEEIAQQIEEIAKQVKKIAWQLKKIAQGPDSV<br>QLEEIAQQIEEIAKQVKKIAQQLKKIAQGGTSGG<br>QLEEIAQQGEEAAKQAKKIAQQLKKIAQ |
| sc-apCC-4-DPH-2 | 8a3k | MGSSHHHHHHSSGLVPRGSHM<br>QLEEIAQQIEEIAKQVKKIAWQLKKIAQGEPSAQG<br>QLEEIAQQIEEIAKQVKKIAWQLKKIAQGPDSV<br>QLEEIAQQGEEAAKQAKKIAQQLKKIAQGGTSGG<br>QLEEIAQQGEEAAKQAKKIAQQLKKIAQ |
| sc-apCC-4-NR-1 | 8a3k | MGSSHHHHHHSSGENLYFQSHM<br>QLEEIAQQLEEGAKQAKKAAWQLKKIAQGEPSAQG<br>QLEEIAQQVEEIAKQIKKIAWQLKKIAQGPDSV<br>QLEEIAQQLEEIAKQVKKIAWQLKKIAQGGTSGG<br>QLEEIAQQAEEGAKQAKKIAWQLKKIAQ |
| sc-apCC-4-NR-2 | 8a3k | MGSSHHHHHHSSGENLYFQSHM<br>QLEEIAQQLEEGAKQGKKLAWQLKKIAQGEPSAQG<br>QLEEIAQQGEEIAKQGKKIAWQLKKIAQGPDSV<br>QLEEIAQQLEEIAKQVKKIAWQLKKIAQGGTSGG<br>QLEEIAQQIEEGAKQIKKIAWQLKKIAQ |
| sc-apCC-4-LMF-1 | 8a3k | MGSSHHHHHHSSGLVPRGSHM<br>QLEEIAQQSEEGAKQGKKIAWQLKKIAQGEPSAQG<br>QLEEIAQQLEEIAKQSKKIAWQLKKIAQGPDSV<br>QLEEIAQQVEEIAKQLKKIAWQLKKIAQGGTSGG<br>QLEEIAQQLEEGAKQAKKTAWQLKKIAQ |
| sc-apCC-4-LMF-2 | 8a3k | MGSSHHHHHHSSGENLYFQSHM<br>QLEEIAQQTEEGAKQAKKIAWQLKKIAQGEPSAQG<br>QLEEIAQQLEEIAKQSKKIAWQLKKIAQGPDSV |

| | | |
|---|---|---|
| | | QLEEIAQQTEEIAKQVKKIAWQLKKIAQGGTSGG<br>QLEEIAQQLEEGAKQAKKTAWQLKKIAQ |
| sc-apCC-4-LMF-3 | 8a3k | MGSSHHHHHHSSGENLYFQSHM<br>QLEEIAQQSEEGAKQAKKIAWQLKKIAQGEPSAQG<br>QLEEIAQQLEEIAKQSKKIAWQLKKIAQGPDSV<br>QLEEIAQQNEEIAKQVKKIAWQLKKIAQGGTSGG<br>QLEEIAQQLEEGAKQSKKTAWQLKKIAQ |
| sc-apCC-4-A7F-1 | 8a3k | MGSSHHHHHHSSGLVPRGSHM<br>QLEEIAQQLEEGAKQTKKVAWQLKKIAQGEPSAQG<br>QLEEIAQQVEEIAKQLKKIAWQLKKIAQGPDSV<br>QLEEIAQQLEEIAKQSKKIAWQLKKIAQGGTSGG<br>QLEEIAQQGEEGAKQAKKIAWQLKKIAQ |
| sc-apCC-4-A7F-2 | 8a3k | MGSSHHHHHHSSGLVPRGSHM<br>QLEEIAQQLEEGAKQAKKGAWQLKKIAQGEPSAQG<br>QLEEIAQQGEEIAKQLKKIAWQLKKIAQGPDSV<br>QLEEIAQQLEEIAKQVKKIAWQLKKIAQGGTSGG<br>QLEEIAQQTEEGAKQLKKIAWQLKKIAQ |
| sc-apCC-4-SN38-1 | 8a3k | MGSSHHHHHHSSGLVPRGSHMTSME<br>QLEEIAEQLEEGAQQAKKNAKQLKKIAKGEPSAQGE<br>QLEEIAEQAEEIAQQSKKIAKQLKKIAKGPDSVE<br>QLEEIAEQLEEIAQQSKKGAKQLKKIAKGGTWSGGE<br>QLEEIAEQAEEGAQQAKKIAKQLKKIAK |
| sc-apCC-4-SN38-2 | 8a3k | MGSSHHHHHHSSGLVPRGSHMTSME<br>QLEEIAEQLEEGAQQAKKNAKQLKKIAKGEPSAQGE<br>QLEEIAEQAEEIAQQTKKIAKQLKKIAKGPDSVE<br>QLEEIAEQLEEIAQQSKKGAKQLKKIAKGGTWSGGE<br>QLEEIAEQAEEGAQQAKKIAKQLKKIAK |
| sc-apCC-4-SN38-3 | 8a3k | MGSSHHHHHHSSGLVPRGSHMTSME<br>QLEEIAEQLEEGAQQAKKAAKQLKKIAKGEPSAQGE<br>QLEEIAEQGEEIAQQSKKIAKQLKKIAKGPDSVE<br>QLEEIAEQLEEIAQQSKKNAKQLKKIAKGGTWSGGE<br>QLEEIAEQAEEGAQQAKKIAKQLKKIAK |
| sc-apCC-4-SN38-4 | 8a3k | MGSSHHHHHHSSGLVPRGSHMTSME<br>QLEEIAEQLEEGAQQAKKNAKQLKKIAKGEPSAQGE<br>QLEEIAEQAEEIAQQSKKIAKQLKKIAKGPDSVE<br>QLEEIAEQLEEIAQQSKKGAKQLKKIAKGGTWSGGE<br>QLEEIAEQAEEGAQQAKKIAKQLKKIAK |
| sc-apCC-6-DPH-NR | 8qad | MGSSHHHHHHSSGLVPRGSHM<br>LFEEIAQLLEEIAKLVKKIAWLVKKIAQGAQPLEM<br>LFEEIAQLAEEIAKLFKKIAWLLKKIAQTTETKKQGD<br>LFEEIAQLLEEIAKLIKKIAWLIKKIAQGYGDKRT<br>LFEEIAQLFEEIAKLFKKIAWLLKKIAQVAPTQRHRY<br>LFEEIAQLLEEIAKLFKKIAWLFKKIAQGTNSDSDLKS<br>LFEEIAQLVEEIAKLFKKIAWLLKKIAQG |
| sc-apCC-6-NR | 8qad | MGSSHHHHHHSSGLVPRGSHM<br>LFEEIAQLLEEIAKLIKKIAWLFKKIAQGAQPLEM<br>LFEEIAQLAEEIAKLFKKIAWLLKKIAQTTETKKQGD<br>LFEEIAQLLEEIAKLFKKIAWLIKKIAQGYGDKRT<br>LFEEIAQLIEEIAKLFKKIAWLLKKIAQVAPTQRHRY<br>LFEEIAQLLEEIAKLFKKIAWLGKKIAQGTNSDSDLKS |

| | | |
|---|---|---|
| | | LFEEIAQLVEEIAKLFKKIAWLLKKIAQG |
| sc-apCC-4-DPH-1-<br>sc-apCC-4-NR-1 | 2 x<br>8a3k | MGSSHHHHHHSSGENLYFQSHMM<br>QLEEIAQQGEEAAKQAKKIAWQLKKIAQGEPSAQG<br>QLEEIAQQIEEIAKQVKKIAWQLKKIAQGPDSV<br>QLEEIAQQIEEIAKQVKKIAQQLKKIAQGGTSGG<br>QLEEIAQQGEEAAKQAKKIAQQLKKIAQQLEQIAM<br>QLEEIAQQLEEGAKQGKKLAWQLKKIAQGEPSAQG<br>QLEEIAQQGEEIAKQGKKIAWQLKKIAQGPDSV<br>QLEEIAQQLEEIAKQVKKIAWQLKKIAQGGTSGG<br>QLEEIAQQIEEGAKQIKKIAWQLKKIAQ |

**Supplementary Table 4**. Experimentally determined dissociation constants $K_D$ (µM) and derived from AutoDock Vinda scores (in parentheses) for RASSCCoL proteins in this study.

| Name | DPH | Nile red | Lumiflavin | Arcyriaflavin A | SN38 |
|---|---|---|---|---|---|
| sc-apCC-4-Ala | > 50<br>(600) | > 20<br>(-) | no binding<br>(-) | 1.8 ± 0.3<br>(-) | no binding<br>(-) |
| sc-apCC-6-Leu | 6.1 ± 1.9<br>(0.03) | > 20<br>(2) | | | |
| sc-apCC-4-DPH-1 | 1.3 ± 0.1<br>(0.02) | > 30<br>(20) | no binding<br>(80) | no binding<br>(2) | no binding<br>(-) |
| sc-apCC-4-DPH-2 | 2.0 ± 0.2<br>(0.01) | | | | |
| sc-apCC-4-NR-1 | > 15<br>(10) | 2.4 ± 0.1<br>(0.04) | no binding<br>(0.05) | 0.050 ± 0.020<br>(0.3) | no binding<br>(30) |
| sc-apCC-4-NR-2 | | 26 ± 3<br>(0.007) | | | |
| sc-apCC-4-LMF-1 | > 70<br>(10) | 1.2 ± 0.1<br>(0.5) | > 50<br>(0.01) | 0.13 ± 0.02<br>(0.004) | no binding<br>(32) |
| sc-apCC-4-LMF-2 | | | no binding<br>(0.008) | | |
| sc-apCC-4-LMF-3 | | | no binding<br>(0.002) | | |
| sc-apCC-4-A7F-1 | no binding<br>(10) | 2.0 ± 0.2<br>(0.2) | 9.6 ± 0.2<br>(0.05) | 0.11 ± 0.02<br>(0.001) | no binding<br>(2) |
| sc-apCC-4-A7F-2 | | | | 6.5 ± 0.3<br>(0.002) | |
| sc-apCC-4-SN38-1 | 4.4 ± 0.7<br>(0.3) | 0.22 ± 0.04<br>(2) | > 90<br>(0.4) | > 20<br>(0.02) | 7.8 ± 0.6<br>(100) |
| sc-apCC-4-SN38-2 | | | | | no binding<br>(50) |
| sc-apCC-4-SN38-3 | | | | | no binding<br>(1) |
| sc-apCC-4-SN38-4 | | | | | no binding<br>(100) |
| sc-apCC-6-DPH-NR | 1.9 ± 0.2<br>(0.06) | 2.2 ± 0.2<br>(0.01) | | | |
| sc-apCC-6-NR | > 20<br>(5) | 0.11 ± 0.04<br>(0.04) | | | |
| sc-apCC-4-DPH-1-<br>sc-apCC-4-NR-1 | 1.3 ± 0.4<br>(0.02) | 0.53 ± 0.05<br>(0.01) | | | |

**Supplementary Table 5.** Boltz2 metrics for LigandMPNN-refined proteins in this study.

| Name | iPDE | ipTM | bb RMSD to parent, Å | Sequence identity, % |
|---|---|---|---|---|
| sc-apCC-4-DPH-3 | 0.45 | 0.97 | 0.3 | 29.5 |
| sc-apCC-4-DPH-4 | 0.45 | 0.97 | 0.6 | 31.2 |
| sc-apCC-4-NR-3 | 0.32 | 0.99 | 0.5 | 27.0 |
| sc-apCC-4-NR-4 | 0.33 | 0.98 | 0.5 | 28.8 |
| sc-apCC-4-LMF-4 | 0.27 | 0.99 | 0.5 | 27.6 |
| sc-apCC-4-LMF-5 | 0.29 | 0.99 | 0.5 | 30.1 |
| sc-apCC-4-SN38-5 | 0.29 | 0.99 | 0.7 | 27.6 |
| sc-apCC-4-SN38-6 | 0.29 | 0.99 | 0.5 | 22.9 |
| sc-apCC-4-SN38-7 | 0.29 | 0.99 | 0.5 | 28.1 |
| sc-apCC-4-SN38-8 | 0.3 | 0.99 | 0.4 | 23.4 |
| sc-apCC-4-A7F-3 | 0.29 | 0.99 | 0.6 | 26.4 |
| sc-apCC-4-A7F-4 | 0.31 | 0.99 | 0.5 | 25.2 |

**Supplementary Table 6.** Expressed LigandMPNN-refined protein sequences for *in vitro* characterisation in this manuscript.

| Name | Scaffold | Sequence |
|---|---|---|
| sc-apCC-4-DPH-3 | 8a3k | MGSSHHHHHHSSGENLYFQSGS<br>ELEEIADKGREIAREGREIAWRLREIAEEDPEHRE<br>RLLAIADRCEEIARRAEEIAWRLREGVRRGGLSPE<br>ELNAIADELEAIADEVEAIAWELREIAVGNPHGE<br>ELRAIADRCREIAAECRAIAWRLREIAE |
| sc-apCC-4-DPH-4 | 8a3k | MGSSHHHHHHSSGENLYFQSGS<br>ELLEIADRGEEIADRGDEIAWRLREIAAKDPENRE<br>ELEAIAERCREIAERGREIAWRLREGVERGGLSPE<br>ELRRIAAEIEEIAEELREIAWRLREIAEGNPHGE<br>ELREIADEMEEIAEELREIAWRLREIAE |
| sc-apCC-4-NR-3 | 8a3k | MGSSHHHHHHSSGENLYFQSGS<br>ELLAIAEELRRLAAEGRALAWRLREIAAKDPENAE<br>ELNAIAAQGEAIAARSEEIAWRLEEGVARGGLSPE<br>ELRAIAAELEAIAAEAEAIAWRLERIADGNPFGA<br>ELRALADRARELAAEGRAIAWRLRELAA |
| sc-apCC-4-NR-4 | 8a3k | MGSSHHHHHHSSGENLYFQSGS<br>ELLAIADELERIAERCRENAWRLREIAAEDPASRA<br>ALEAIAARLEEIAERGREIAWRLRDGVARGGLSPE<br>ELRAIADELEAIAAEGEALAWELEAIAEGHPHGA<br>ELRAIADQGRADAAQGRAIAWRLRELAA |

| | | |
|---|---|---|
| sc-apCC-4-LMF-4 | 8a3k | MGSSHHHHHHSSGENLYFQSGS<br>ELLAIADASRAGAAKGREIAWRLREIAAREPAHRA<br>ELEAIADELEAIAARSEEIAWRLRAGVERGGLSPE<br>ELRAIADEIDAIADRLEAIAWRLEEIAATSPSGA<br>ELRAIAAELRAEAAAARANAWKLRELAA |
| sc-apCC-4-LMF-5 | 8a3k | MGSSHHHHHHSSGENLYFQSGS<br>ELREIAEETREGAREGREIAWELRRIAEEDPAHRE<br>RLLAIADRLEEIARESERIAWRILEGVERGGLSPE<br>ELREIADRQEAIAEELRRIAWELREIAEGSPHGA<br>RLREIADRLEEAAERTRENAWRLRRLAE |
| sc-apCC-4-A7F-3 | 8a3k | MGSSHHHHHHSSGENLYFQSGS<br>ELLAIADELRAGAWETRAAADRLREIAAVDPANRA<br>ELEAIADQLRAIAWRLEAIADRLRAGVARGGLSPE<br>ELRAIADELRAIAAESRAIADRLERIADGNPHGA<br>ELRAIADGARAGAARAEEIAARLEELAA |
| sc-apCC-4-A7F-4 | 8a3k | MGSSHHHHHHSSGENLYFQSGS<br>ELAAIADELEAGADAAEAAAWELERIAAEDPAHAA<br>ELRAIADELRAQAWQLRAIAARLRAGVERGGLSPE<br>ELRAIADELDALADAERATAARLREIAEGNPHGA<br>RLRAIAAGAEAGAARARAIAARLRALAA |
| sc-apCC-4-SN38-5 | 8a3k | MGSSHHHHHHSSGENLYFQSGS<br>ELEAIAAQLEAAAWQARAAAAQLRAIAAQDPANGA<br>ALNAIADAGDAVADEARAIAWELRAAVAAGGPTAE<br>QLRAIAARLEAIAKETEAAADQLRAIAEGNPHGA<br>ELRALADQIRAGAAQARAIAAQLRALAA |
| sc-apCC-4-SN38-6 | 8a3k | MGSSHHHHHHSSGENLYFQSGS<br>ELLAIADQLRANAWAARAAAAALERIAAVDPANRA<br>ALLAIAWEGHAVADEATAIADELRAAVAAGGPTPA<br>QLRAIADRLDAIAAATAAAAAQLRAIAVGNPHGA<br>ELNAIAAAVAAGATAATAIAAQLRALAA |
| sc-apCC-4-SN38-7 | 8a3k | MGSSHHHHHHSSGENLYFQSGS<br>ELEAIAAELEAAADEAEAHAWRLRAIAAKDPANRA<br>ALEAIAWKGEEVAARAREIAAQLRDAVAAGGPTAA<br>QLRAIAAQLEEIARETRAAAAQLREIADGNPFGA<br>ELRAIADETEKGAEKARAIAARLRELAE |
| sc-apCC-4-SN38-8 | 8a3k | MGSSHHHHHHSSGENLYFQSGS<br>ELLAIADQLRANAWAARAAAAALEAIAARTPADAA<br>ALRALAAAGRATAWAAEAIAQQLRDAVARGGPTAA<br>QLNAIAAQLDAIAAATTAAAAELRALAVGNPAAA<br>ELRAIADAVAAGATETAAIAAQLRAIAA |

**Supplementary Table 7.** Crystallisation conditions for crystal structures in this study.

| Protein | Condition name | Condition |
|---|---|---|
| sc-apCC-4-DPH-2 | JCSG-plus™ D9 | 0.17 M Ammonium sulfate, 25.5% w/v Polyethylene glycol 4,000, 15% v/v Glycerol |
| sc-apCC-4-NR-1 | Pact Premier™ D2 | 0.1 M MMT buffer pH 5.0, 25 % w/v PEG 1500 |
| sc-apCC-4-LMF-1 | JCSG-plus™ A12 | 0.2 M Potassium nitrate, 20% w/v Polyethylene glycol 3,350 |
| sc-apCC-4-A7F-1 | Pact Premier™ C7 | 0.2 M sodium chloride, 0.1 M HEPES pH 7.0, 20 % w/v PEG 6000 |
| sc-apCC-4-SN38-1 | Pact Premier™ A3 | 0.1 M SPG buffer pH 6.0, 25 % w/v PEG 1500 |
| sc-apCC-6-DPH-NR | Pact Premier™ B11 | 0.01 M zinc chloride, 0.1 M MES pH 6.0, 20 % w/v PEG 6000 |

**Supplementary Table 8.** Merging and refinement statistics for crystal structures in this study.

| | sc-apCC-4-DPH-2 | sc-apCC-4-NR-1 | sc-apCC-4-LMF-1 |
|---|---|---|---|
| PDB id | 9R1N | 9R1J | 9R1M |
| Wavelength | 0.6199 Å | 0.6199 Å | 0.9999 Å |
| Resolution range | 42.2 - 2.4 (2.486 - 2.4) | 27.82 - 1.702 (1.763 - 1.702) | 31.33 - 2.4 (2.486 - 2.4) |
| Space group | P 65 2 2 | P 1 21 1 | P 1 21 1 |
| Unit cell | 68.1003 68.1003 120.793 90 90 120 | 27.662 38.1608 56.6897 90 101.082 90 | 28.047 38.26 55.455 90 100.172 90 |
| Total reflections | 636957 (64114) | 73226 (3364) | 8829 (822) |
| Unique reflections | 6975 (674) | 12141 (904) | 4572 (431) |
| Multiplicity | 91.3 (95.1) | 6.0 (3.7) | 1.9 (1.9) |
| Completeness (%) | 99.89 (100.00) | 94.02 (70.06) | 98.62 (97.07) |
| Mean I/sigma(I) | 18.12 (1.89) | 25.48 (5.36) | 3.90 (1.45) |
| Wilson B-factor | 52.29 | 21.54 | 50.63 |
| R-merge | 0.2364 (2.615) | 0.05404 (0.1202) | 0.07497 (0.4823) |
| R-meas | 0.2377 (2.629) | 0.05866 (0.1388) | 0.106 (0.682) |
| R-pim | 0.02488 (0.2674) | 0.0226 (0.06724) | 0.07497 (0.4823) |
| CC1/2 | 1 (0.895) | 0.998 (0.983) | 0.984 (0.612) |
| CC* | 1 (0.972) | 1 (0.996) | 0.996 (0.871) |
| Reflections used in refinement | 6969 (674) | 12129 (901) | 4570 (430) |
| Reflections used for R-free | 345 (32) | 610 (49) | 235 (19) |
| R-work | 0.2267 (0.2737) | 0.1905 (0.2137) | 0.2286 (0.3102) |
| R-free | 0.2422 (0.2908) | 0.1881 (0.2109) | 0.2353 (0.2515) |
| CC(work) | 0.951 (0.851) | 0.961 (0.911) | 0.956 (0.728) |
| CC(free) | 0.943 (0.801) | 0.944 (0.948) | 0.912 (0.856) |
| Number of non-hydrogen atoms | 973 | 1104 | 821 |
| macromolecules | 943 | 973 | 821 |
| ligands | 23 | 14 | 0 |

| | | | |
|---|---|---|---|
| solvent | 7 | 125 | 0 |
| Protein residues | 131 | 132 | 129 |
| Nucleic acid bases | | | |
| RMS(bonds) | 0.007 | 0.005 | 0.004 |
| RMS(angles) | 0.79 | 0.52 | 0.55 |
| Ramachandran favored (%) | 100 | 99.23 | 99.21 |
| Ramachandran allowed (%) | 0 | 0.77 | 0.79 |
| Ramachandran outliers (%) | 0 | 0 | 0 |
| Rotamer outliers (%) | 2.6 | 0 | 0 |
| Clashscore | 5.17 | 3.59 | 5.77 |
| Average B-factor | 61.18 | 27.89 | 63.97 |
| macromolecules | 61.03 | 26.72 | 63.97 |
| ligands | 69.25 | 43.69 | - |
| solvent | 54.83 | 36.28 | - |
| Number of TLS groups | 4 | 1 | 5 |

| | sc-apCC-4-A7F-1 | sc-apCC-4-SN38-1 | sc-apCC-6-DPH-NR |
|---|---|---|---|
| PDB id | 9R1K | 9R1L | 9R1O |
| Wavelength | 0.6199 Å | 0.6199 Å | 0.9762 Å |
| Resolution range | 27.91 - 2.2 (2.279 - 2.2) | 27.24 - 1.85 (1.916 - 1.85) | 45.66 - 3.229 (3.344 - 3.229) |
| Space group | P 1 21 1 | P 1 21 1 | P 1 21 1 |
| Unit cell | 27.5551 37.8603 57.0232 90 101.821 90 | 27.7628 37.5806 54.86 90 101.16 90 | 58.699 59.583 64.307 90 96.293 90 |
| Total reflections | 40693 (4149) | 42875 (4053) | 12924 (1169) |
| Unique reflections | 5977 (598) | 9615 (912) | 7018 (654) |
| Multiplicity | 6.8 (6.9) | 4.5 (4.4) | 1.8 (1.8) |

| | | | |
|---|---|---|---|
| Completeness (%) | 99.85 (99.83) | 99.73 (98.59) | 96.97 (93.97) |
| Mean I/sigma(I) | 9.41 (2.25) | 14.39 (1.30) | 3.47 (1.43) |
| Wilson B-factor | 41.85 | 35.34 | 58.96 |
| R-merge | 0.08956 (0.4694) | 0.05523 (0.4269) | 0.1003 (0.4693) |
| R-meas | 0.097 (0.5074) | 0.06274 (0.4868) | 0.1419 (0.6637) |
| R-pim | 0.03696 (0.1915) | 0.02923 (0.2306) | 0.1003 (0.4693) |
| CC1/2 | 0.998 (0.932) | 0.999 (0.876) | 0.995 (0.624) |
| CC* | 0.999 (0.982) | 1 (0.966) | 0.999 (0.877) |
| Reflections used in refinement | 5970 (597) | 9600 (907) | 7004 (654) |
| Reflections used for R-free | 298 (26) | 434 (40) | 322 (35) |
| R-work | 0.2175 (0.2449) | 0.2001 (0.2885) | 0.2652 (0.3194) |
| R-free | 0.2530 (0.4221) | 0.2142 (0.2839) | 0.2853 (0.3528) |
| CC(work) | 0.963 (0.906) | 0.970 (0.854) | 0.924 (0.739) |
| CC(free) | 0.901 (0.872) | 0.942 (0.892) | 0.772 (0.570) |
| Number of non-hydrogen atoms | 898 | 1020 | 2860 |
| macromolecules | 881 | 989 | 2847 |
| ligands | 4 | 17 | 13 |
| solvent | 13 | 24 | 0 |
| Protein residues | 126 | 134 | 389 |
| Nucleic acid bases | | | |
| RMS(bonds) | 0.005 | 0.003 | 0.001 |
| RMS(angles) | 0.62 | 0.57 | 0.26 |
| Ramachandran favored (%) | 100 | 99.22 | 99.45 |
| Ramachandran allowed (%) | 0 | 0.78 | 0.55 |
| Ramachandran outliers (%) | 0 | 0 | 0 |
| Rotamer outliers (%) | 3.23 | 0 | 0 |
| Clashscore | 4.7 | 3.58 | 3.51 |

| | | | |
|---|---|---|---|
| Average B-factor | 53.81 | 46.45 | 47.04 |
| macromolecules | 53.82 | 46.28 | 47.07 |
| ligands | 61.14 | 59.44 | 39.56 |
| solvent | 50.95 | 49.73 | - |
| Number of TLS groups | 1 | 3 | 2 |

**Supplementary Table 9. Simultaneous FRET lifetime fits from DPH and Nile red channels after excitation at 352 nm.** Floated parameters in bold. Fluorescence decays were fitted to bi-exponential decay equations while keeping average DPH and Nile red lifetimes fixed. DPH population $A_2$ is not involved in FRET; the ratio $A_1/(A_1+A_2)$ represents the fraction of DPH-bound proteins undergoing FRET.

| | |
|---|---|
| ***A*₁(DPH)** | **0.24** |
| ***A*₂(DPH)** | **0.80** |
| ***A*₂(NR)** | **3.2** |
| **$t_{FRET}$** | **2.0 ± 0.2 ns** |
| $t_{DPH}$ | 7.6 ± 0.2 ns |
| $t_{NR}$ | 5.2 ± 0.2 ns |

**Supplementary Table 10.** Spectral properties of Nile red and Nile blue bound to sc-apCC-4-SN38-1. Absolute quantum yield was measured with an integrating sphere (Nile red $\lambda_{ex}$ = 510 nm, Nile blue $\lambda_{ex}$ = 530 nm). Extinction coefficient was derived through comparison with a known standard – Nile red and Nile blue in methanol.

| | Φ | τ | ε | $\lambda_{abs}$ max | $\lambda_{em}$ max |
|---|---|---|---|---|---|
| Nile red | 0.90 ± 0.02 | 4.8 ± 0.2 ns | 52 400 $cm^{-1}$ $M^{-1}$ at 552 nm | 559 nm | 607 nm |
| Nile blue | 0.74 ± 0.02 | 4.2 ± 0.2 ns | 34 400 $cm^{-1}$ $M^{-1}$ at 627 nm | 620 nm | 645 nm |

**Supplementary Table 11.** Expressed protein sequences for in-cell characterisation in this manuscript.

| Name | Sequence |
|---|---|
| sc-apCC-4-SN38-1 : mEmerald | MQLEEIAQQLEEGAKQTKKNAWQLKKIAQGEPSAQGQLEEI AQQAEEIAKQSKKIAWQLKKIAQGPDSVQLEEIAQQLEEIA KQAKKGAWQLKKIAQGGTSGGQLEEIAQQAEEGAKQAKKIA WQLKKIAQGSGSGMVSKGEELFTGVVPILVELDGDVNGHKF SVSGEGEGDATYGKLTLKFICTTGKLPVPWPTLVTTLTYGV QCFARYPDHMKQHDFFKSAMPEGYVQERTIFFKDDGNYKTR AEVKFEGDTLVNRIELKGIDFKEDGNILGHKLEYNYNSHKV YITADKQKNGIKVNFKTRHNIEDGSVQLADHYQQNTPIGDG PVLLPDNHYLSTQSKLSKDPNEKRDHMVLLEFVTAAGITLG MDELYK* |
| NLS : sc-apCC-4-SN38-1 : mEmerald | MPKKKRKVTGSTGSGTMQLEEIAQQLEEGAKQTKKNAWQLK KIAQGEPSAQGQLEEIAQQAEEIAKQSKKIAWQLKKIAQGP DSVQLEEIAQQLEEIAKQAKKGAWQLKKIAQGGTSGGQLEE IAQQAEEGAKQAKKIAWQLKKIAQGSGSGMVSKGEELFTGV VPILVELDGDVNGHKFSVSGEGEGDATYGKLTLKFICTTGK LPVPWPTLVTTLTYGVQCFARYPDHMKQHDFFKSAMPEGYV QERTIFFKDDGNYKTRAEVKFEGDTLVNRIELKGIDFKEDG NILGHKLEYNYNSHKVYITADKQKNGIKVNFKTRHNIEDGS |

| | |
|---|---|
| | VQLADHYQQNTPIGDGPVLLPDNHYLSTQSKLSKDPNEKRD<br>HMVLLEFVTAAGITLGMDELYK* |
| Fyn : sc-apCC-4-SN38-1 :<br><br>mEmerald | MGCVQCKDKEATKLTEERDGSLNQTSGSGSSRGRSRPPEGT<br>MQLEEIAQQLEEGAKQTKKNAWQLKKIAQGEPSAQGQLEEI<br>AQQAEEIAKQSKKIAWQLKKIAQGPDSVQLEEIAQQLEEIA<br>KQAKKGAWQLKKIAQGGTSGGQLEEIAQQAEEGAKQAKKIA<br>WQLKKIAQGSGSGMVSKGEELFTGVVPILVELDGDVNGHKF<br>SVSGEGEGDATYGKLTLKFICTTGKLPVPWPTLVTTLTYGV<br>QCFARYPDHMKQHDFFKSAMPEGYVQERTIFFKDDGNYKTR<br>AEVKFEGDTLVNRIELKGIDFKEDGNILGHKLEYNYNSHKV<br>YITADKQKNGIKVNFKTRHNIEDGSVQLADHYQQNTPIGDG<br>PVLLPDNHYLSTQSKLSKDPNEKRDHMVLLEFVTAAGITLG<br>MDELYK* |
| LifeAct : sc-apCC-4-SN38-1 :<br><br>mEmerald | MGVADLIKKFESISKEETGSTGSGTMQLEEIAQQLEEGAKQ<br>TKKNAWQLKKIAQGEPSAQGQLEEIAQQAEEIAKQSKKIAW<br>QLKKIAQGPDSVQLEEIAQQLEEIAKQAKKGAWQLKKIAQG<br>GTSGGQLEEIAQQAEEGAKQAKKIAWQLKKIAQGSGSGMVS<br>KGEELFTGVVPILVELDGDVNGHKFSVSGEGEGDATYGKLT<br>LKFICTTGKLPVPWPTLVTTLTYGVQCFARYPDHMKQHDFF<br>KSAMPEGYVQERTIFFKDDGNYKTRAEVKFEGDTLVNRIEL<br>KGIDFKEDGNILGHKLEYNYNSHKVYITADKQKNGIKVNFK<br>TRHNIEDGSVQLADHYQQNTPIGDGPVLLPDNHYLSTQSKL<br>SKDPNEKRDHMVLLEFVTAAGITLGMDELYK* |
| sc-apCC-4-NR-1 : mEmerald | MQLEEIAQQLEEGAKQAKKAAWQLKKIAQGEPSAQGQLEEI<br>AQQVEEIAKQIKKIAWQLKKIAQGPDSVQLEEIAQQLEEIA<br>KQVKKIAWQLKKIAQGGTSGGQLEEIAQQAEEGAKQAKKIA<br>WQLKKIAQGSGSGMVSKGEELFTGVVPILVELDGDVNGHKF<br>SVSGEGEGDATYGKLTLKFICTTGKLPVPWPTLVTTLTYGV<br>QCFARYPDHMKQHDFFKSAMPEGYVQERTIFFKDDGNYKTR<br>AEVKFEGDTLVNRIELKGIDFKEDGNILGHKLEYNYNSHKV<br>YITADKQKNGIKVNFKTRHNIEDGSVQLADHYQQNTPIGDG<br>PVLLPDNHYLSTQSKLSKDPNEKRDHMVLLEFVTAAGITLG<br>MDELYK* |
| NLS : sc-apCC-4-NR-1 :<br><br>mEmerald | MPKKKRKVTGSTGSGTMQLEEIAQQLEEGAKQAKKAAWQLK<br>KIAQGEPSAQGQLEEIAQQVEEIAKQIKKIAWQLKKIAQGP<br>DSVQLEEIAQQLEEIAKQVKKIAWQLKKIAQGGTSGGQLEE<br>IAQQAEEGAKQAKKIAWQLKKIAQGSGSGMVSKGEELFTGV<br>VPILVELDGDVNGHKFSVSGEGEGDATYGKLTLKFICTTGK<br>LPVPWPTLVTTLTYGVQCFARYPDHMKQHDFFKSAMPEGYV<br>QERTIFFKDDGNYKTRAEVKFEGDTLVNRIELKGIDFKEDG<br>NILGHKLEYNYNSHKVYITADKQKNGIKVNFKTRHNIEDGS<br>VQLADHYQQNTPIGDGPVLLPDNHYLSTQSKLSKDPNEKRD<br>HMVLLEFVTAAGITLGMDELYK* |
| Fyn : sc-apCC-4-NR-1 :<br><br>mEmerald | MGCVQCKDKEATKLTEERDGSLNQTSGSGSSRGRSRPPEGT<br>MQLEEIAQQLEEGAKQAKKAAWQLKKIAQGEPSAQGQLEEI<br>AQQVEEIAKQIKKIAWQLKKIAQGPDSVQLEEIAQQLEEIA<br>KQVKKIAWQLKKIAQGGTSGGQLEEIAQQAEEGAKQAKKIA<br>WQLKKIAQGSGSGMVSKGEELFTGVVPILVELDGDVNGHKF<br>SVSGEGEGDATYGKLTLKFICTTGKLPVPWPTLVTTLTYGV<br>QCFARYPDHMKQHDFFKSAMPEGYVQERTIFFKDDGNYKTR<br>AEVKFEGDTLVNRIELKGIDFKEDGNILGHKLEYNYNSHKV |

| | |
|---|---|
| | YITADKQKNGIKVNFKTRHNIEDGSVQLADHYQQNTPIGDG PVLLPDNHYLSTQSKLSKDPNEKRDHMVLLEFVTAAGITLG MDELYK* |
| LifeAct : sc-apCC-4-NR-1 : mEmerald | MGVADLIKKFESISKEETGSTGSGTMQLEEIAQQLEEGAKQ AKKAAWQLKKIAQGEPSAQGQLEEIAQQVEEIAKQIKKIAW QLKKIAQGPDSVQLEEIAQQLEEIAKQVKKIAWQLKKIAQG GTSGGQLEEIAQQAEEGAKQAKKIAWQLKKIAQGSGSGMVS KGEELFTGVVPILVELDGDVNGHKFSVSGEGEGDATYGKLT LKFICTTGKLPVPWPTLVTTLTYGVQCFARYPDHMKQHDFF KSAMPEGYVQERTIFFKDDGNYKTRAEVKFEGDTLVNRIEL KGIDFKEDGNILGHKLEYNYNSHKVYITADKQKNGIKVNFK TRHNIEDGSVQLADHYQQNTPIGDGPVLLPDNHYLSTQSKL SKDPNEKRDHMVLLEFVTAAGITLGMDELYK* |
| sc-apCC-4 : mEmerald | MQLEEIAQQLEEIAKQLKKIAWQLKKIAQGEPSAQGQLEEI AQQLEEIAKQLKKIAWQLKKIAQGPDSVQLEEIAQQLEEIA KQLKKIAWQLKKIAQGGTSGGQLEEIAQQLEEIAKQLKKIA WQLKKIAQGSGSGMVSKGEELFTGVVPILVELDGDVNGHKF SVSGEGEGDATYGKLTLKFICTTGKLPVPWPTLVTTLTYGV QCFARYPDHMKQHDFFKSAMPEGYVQERTIFFKDDGNYKTR AEVKFEGDTLVNRIELKGIDFKEDGNILGHKLEYNYNSHKV YITADKQKNGIKVNFKTRHNIEDGSVQLADHYQQNTPIGDG PVLLPDNHYLSTQSKLSKDPNEKRDHMVLLEFVTAAGITLG MDELYK* |
| NLS : sc-apCC-4 : mEmerald | MQLEEIAQQLEEIAKQLKKIAWQLKKIAQGEPSAQGQLEEI AQQLEEIAKQLKKIAWQLKKIAQGPDSVQLEEIAQQLEEIA KQLKKIAWQLKKIAQGGTSGGQLEEIAQQLEEIAKQLKKIA WQLKKIAQGSGSGMVSKGEELFTGVVPILVELDGDVNGHKF SVSGEGEGDATYGKLTLKFICTTGKLPVPWPTLVTTLTYGV QCFARYPDHMKQHDFFKSAMPEGYVQERTIFFKDDGNYKTR AEVKFEGDTLVNRIELKGIDFKEDGNILGHKLEYNYNSHKV YITADKQKNGIKVNFKTRHNIEDGSVQLADHYQQNTPIGDG PVLLPDNHYLSTQSKLSKDPNEKRDHMVLLEFVTAAGITLG MDELYK* |
| Fyn : sc-apCC-4 : mEmerald | MGCVQCKDKEATKLTEERDGSLNQTSGSGSSRGRSRPPEGT MQLEEIAQQLEEIAKQLKKIAWQLKKIAQGEPSAQGQLEEI AQQLEEIAKQLKKIAWQLKKIAQGPDSVQLEEIAQQLEEIA KQLKKIAWQLKKIAQGGTSGGQLEEIAQQLEEIAKQLKKIA WQLKKIAQGSGSGMVSKGEELFTGVVPILVELDGDVNGHKF SVSGEGEGDATYGKLTLKFICTTGKLPVPWPTLVTTLTYGV QCFARYPDHMKQHDFFKSAMPEGYVQERTIFFKDDGNYKTR AEVKFEGDTLVNRIELKGIDFKEDGNILGHKLEYNYNSHKV YITADKQKNGIKVNFKTRHNIEDGSVQLADHYQQNTPIGDG PVLLPDNHYLSTQSKLSKDPNEKRDHMVLLEFVTAAGITLG MDELYK* |
| LifeAct : sc-apCC-4 : mEmerald | MGVADLIKKFESISKEETGSTGSGTMQLEEIAQQLEEIAKQ LKKIAWQLKKIAQGEPSAQGQLEEIAQQLEEIAKQLKKIAW QLKKIAQGPDSVQLEEIAQQLEEIAKQLKKIAWQLKKIAQG GTSGGQLEEIAQQLEEIAKQLKKIAWQLKKIAQGSGSGMVS KGEELFTGVVPILVELDGDVNGHKFSVSGEGEGDATYGKLT LKFICTTGKLPVPWPTLVTTLTYGVQCFARYPDHMKQHDFF KSAMPEGYVQERTIFFKDDGNYKTRAEVKFEGDTLVNRIEL |

| | |
|---|---|
| | KGIDFKEDGNILGHKLEYNYNSHKVYITADKQKNGIKVNFKTRHNIEDGSVQLADHYQQNTPIGDGPVLLPDNHYLSTQSKLSKDPNEKRDHMVLLEFVTAAGITLGMDELYK* |
| sc-apCC-4-SN38-1 | MQLEEIAQQLEEGAKQTKKNAWQLKKIAQGEPSAQGQLEEIAQQAEEIAKQSKKIAWQLKKIAQGPDSVQLEEIAQQLEEIAKQAKKGAWQLKKIAQGGTSGGQLEEIAQQAEEGAKQAKKIAWQLKKIAQGS |
| NLS : sc-apCC-4-SN38-1 | MPKKKRKVTGSTGSGTMQLEEIAQQLEEGAKQTKKNAWQLKKIAQGEPSAQGQLEEIAQQAEEIAKQSKKIAWQLKKIAQGPDSVQLEEIAQQLEEIAKQAKKGAWQLKKIAQGGTSGGQLEEIAQQAEEGAKQAKKIAWQLKKIAQGS |
| Fyn : sc-apCC-4-SN38-1 | MGCVQCKDKEATKLTEERDGSLNQTSGSGSSRGRSRPPEGTMQLEEIAQQLEEGAKQTKKNAWQLKKIAQGEPSAQGQLEEIAQQAEEIAKQSKKIAWQLKKIAQGPDSVQLEEIAQQLEEIAKQAKKGAWQLKKIAQGGTSGGQLEEIAQQAEEGAKQAKKIAWQLKKIAQGS |
| LifeAct : sc-apCC-4-SN38-1 | MGVADLIKKFESISKEETGSTGSGTMQLEEIAQQLEEGAKQTKKNAWQLKKIAQGEPSAQGQLEEIAQQAEEIAKQSKKIAWQLKKIAQGPDSVQLEEIAQQLEEIAKQAKKGAWQLKKIAQGGTSGGQLEEIAQQAEEGAKQAKKIAWQLKKIAQGS |

### 2 Supplementary Figures

#### 2.1 Computational design of 4-helix binders with RASSCCoL

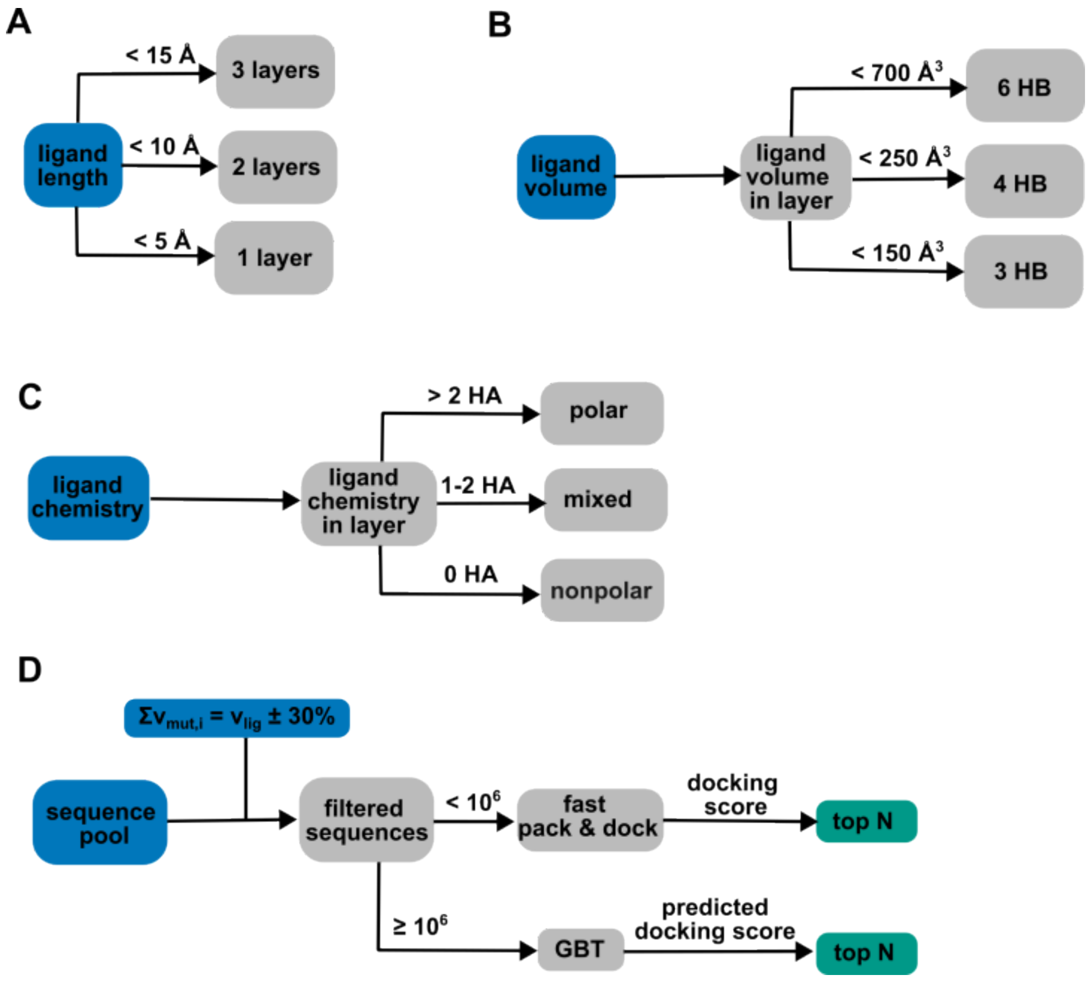


**Supplementary Figure 1. Decision scheme for sequence design and screening.** Design specifications are defined based on ligand size and chemistry. Ligand length **(A)** determines how many layers of the scaffold are modified. Ligand volume **(B)** defines the required size of the scaffold, and ligand chemistry **(C)**, approximated by the number of heteroatoms, determines the amino acid palette used in each layer. These constraints are translated into sequence-level requirements by applying a volumetric matching criterion, generating plausible binding sequences **(D).** The resulting sequences are then evaluated structurally: smaller sets are assessed using fast packing and docking, whereas larger sets are screened using docking score prediction from sequence alone with a gradient-boosted trees (GBT). Top-ranked candidates are selected for further in silico and experimental analysis.

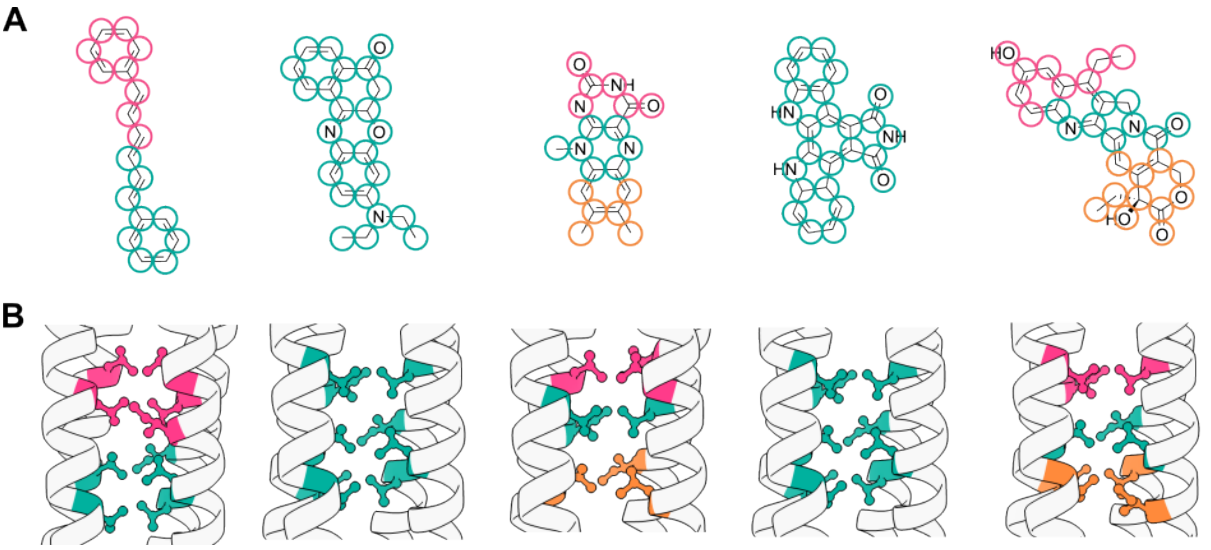


**Supplementary Figure 2. Binding site design approach (A)** Ligands used in the RASSCCoL pipeline—DPH, Nile red, lumiflavin, arcyriaflavin A, and SN38 (left to right)—divided into colored sections that align with layers in the protein scaffold. **(B)** 4-Helix bundle protein scaffolds are shown directly below the corresponding ligands, with binding pocket residues candidates represented as ball-and-stick and colored to match the ligand layers. The length of each binding pocket matches the length of its corresponding ligand, allowing for a layer-based design strategy.

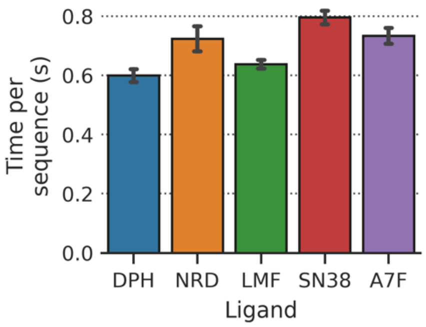


**Supplementary Figure 3. CPU benchmarking for backbone sequence packing and ligand docking.** The time required to pack a single sequence onto sc-apCC-4 using FASPR and dock a ligand using AutoDock Vina with low sampling parameters was measured on a single thread of a Xeon Gold 6226R processor. A total of 100 sequences were tested per ligand. Bar plots represent the mean time for 1 packing and docking cycle, with error bars denoting one standard deviation. The observed times (in seconds per CPU) were as follows: DPH: 0.58 (± 0.02), Nile red (NRD): 0.69 (± 0.04), lumiflavin (LMF): 0.64 (± 0.01), SN38: 0.75 (± 0.02) and arcyriaflavin A (A7F): 0.73 (± 0.03).

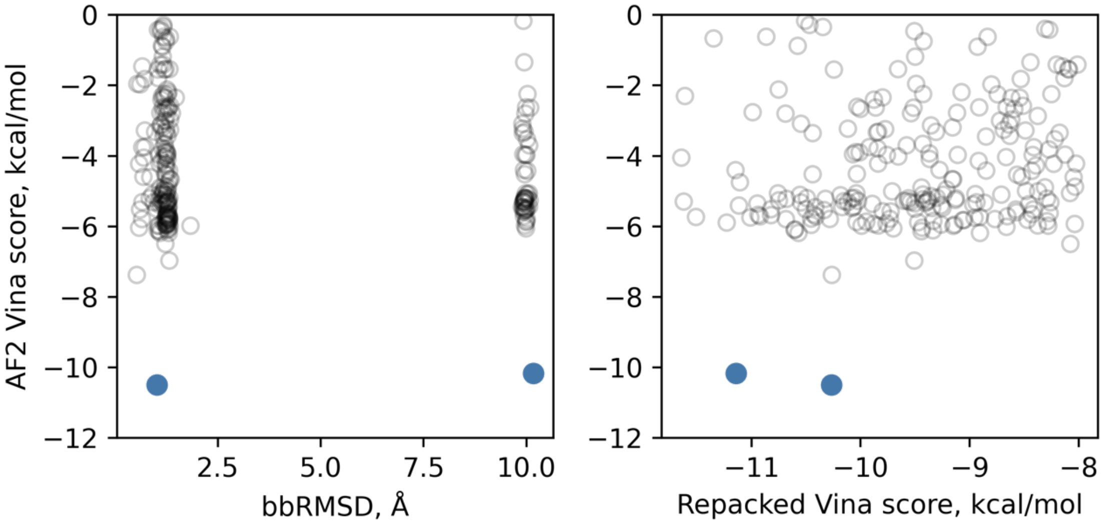


**Supplementary Figure 4. Sequence selection for DPH binders based on AutoDock Vina scores.** Blue circles denote experimentally characterised designs. Top 270 sequences after repacking (< -8.0 kcal/mol) were analysed with AlphaFold2.

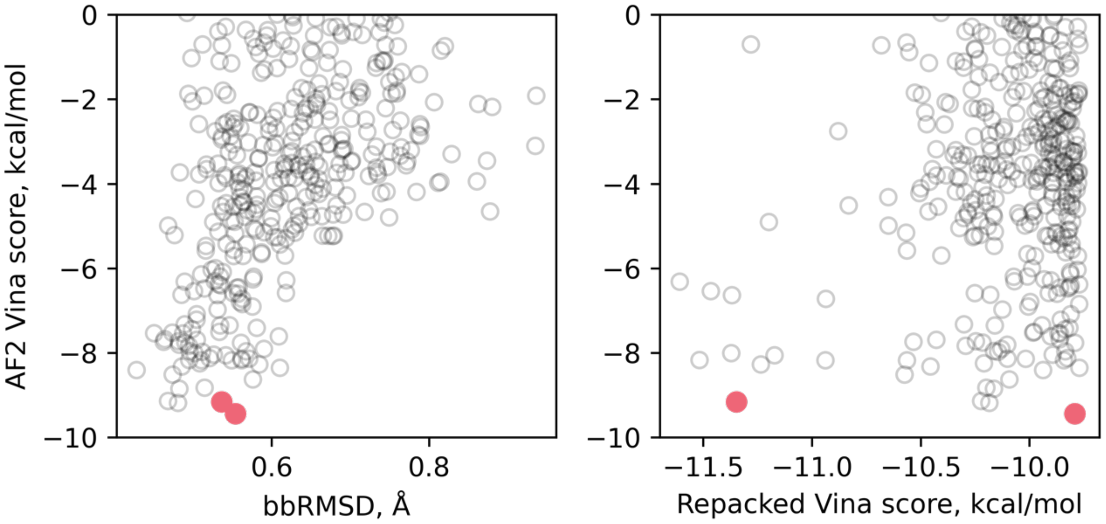


**Supplementary Figure 5. Sequence selection for Nile red binders based on AutoDock Vina scores.** Red circles denote experimentally characterised designs. Top 600 sequences after repacking (< -9.8 kcal/mol) were analysed with AlphaFold2.

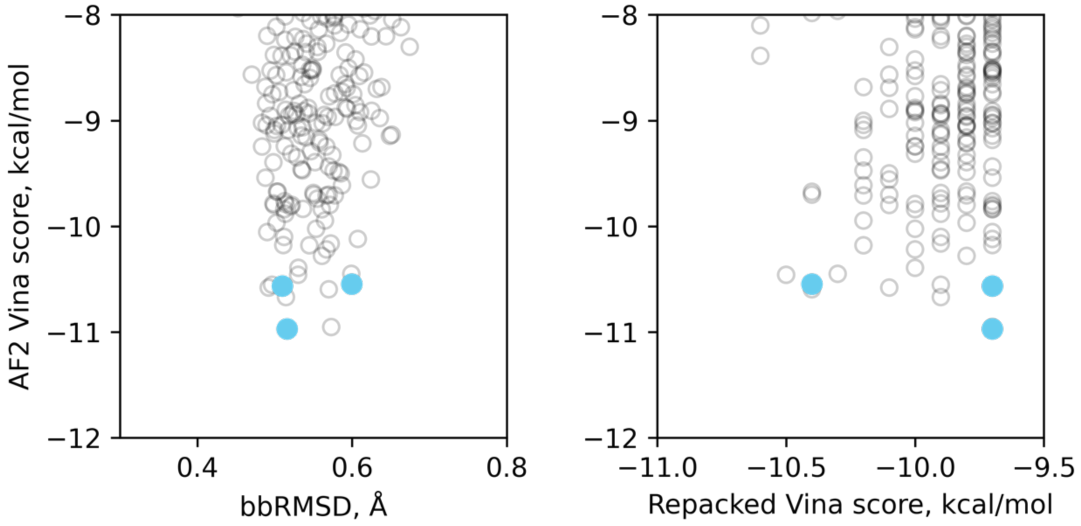


**Supplementary Figure 6. Sequence selection for flavin binders based on AutoDock Vina scores.** Cyan circles denote experimentally characterised designs. Top 359 sequences after repacking (< -9.6 kcal/mol) were analysed with AlphaFold2.

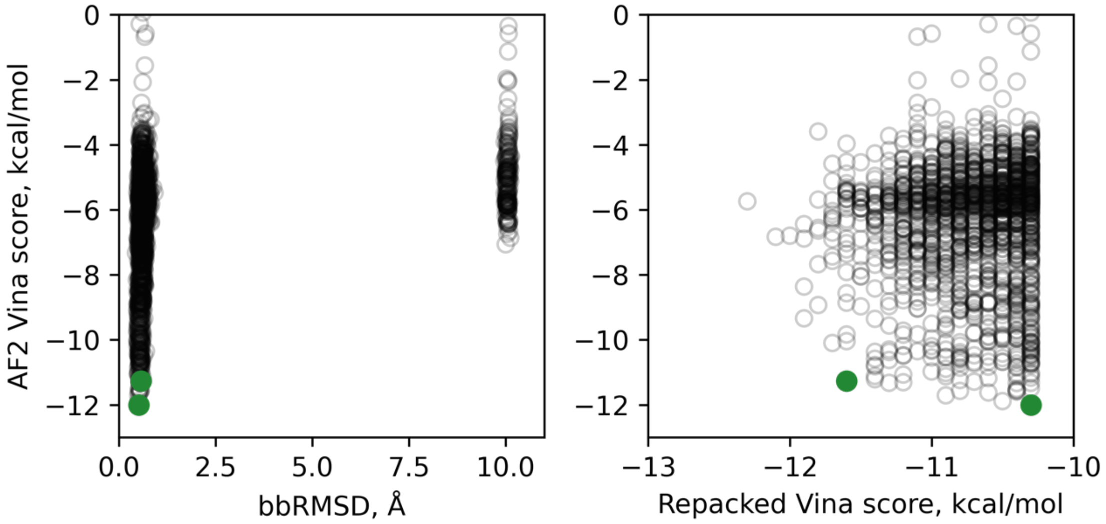


**Supplementary Figure 7. Sequence selection for arcyriaflavin A binders based on AutoDock Vina scores.** Green circles denote experimentally characterised designs. Top 1900 sequences after repacking (< -10.2 kcal/mol) were analysed with AlphaFold2.

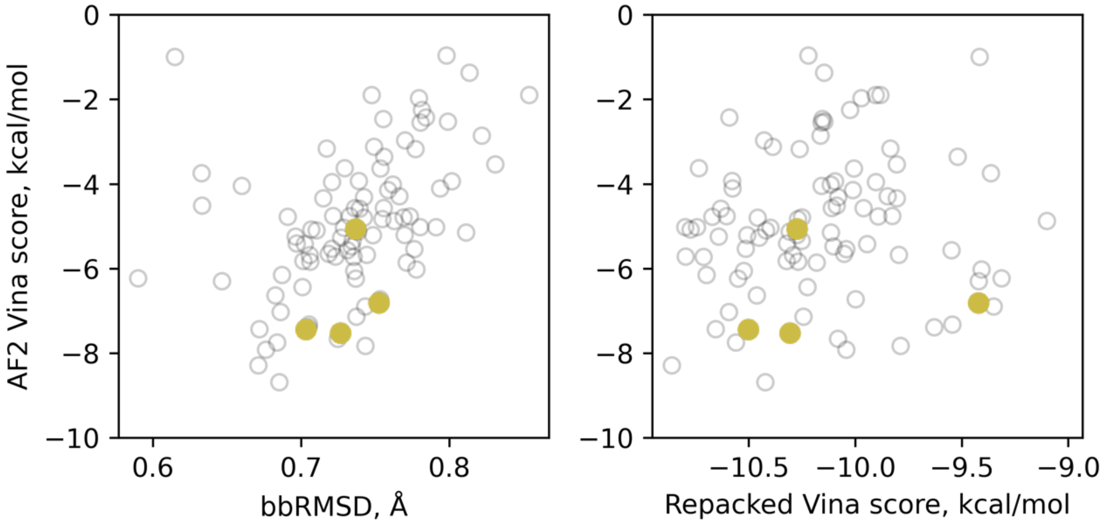


**Supplementary Figure 8. Sequence selection for SN38 binders based on AutoDock Vina scores.** Yellow circles denote experimentally characterised designs. No design passed the filtering threshold after being folded with AlphaFold2 based on DPH, Nile red, flavin or arcyriaflavin A designs (AF2+Viina score < -9.0 kcal/mol, bbRMSD < 0.6 Å). To better account for ligand-protein hydrogen bonding and rotamer flexibility, we used an additional filtering step with ultrafast MD simulations, see Figures S9&S10.

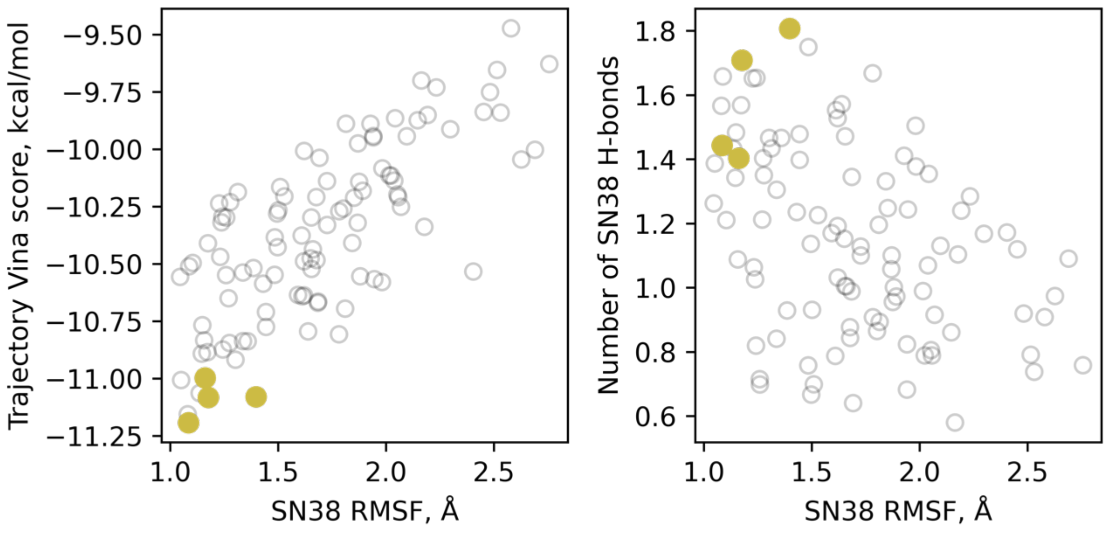


**Supplementary Figure 9. Sequence selection for SN38 binders based on ultrafast 10 x 5 ns MD simulations at 400K.** RMSF: root-mean-square-fluctuation of SN38**.** The system with implicit water was minimised, heated to 400 K over 400 ps, and equilibrated for 1.6 ns prior to a 5 ns production run. To enhance sampling, 10 independent replicates of the minimization, equilibration and production runs were performed. Weak positional restraints of 20 cal mol$^{-1}$Å$^{-2}$ were applied to the backbone atoms to maintain simulation stability at elevated temperatures [1].

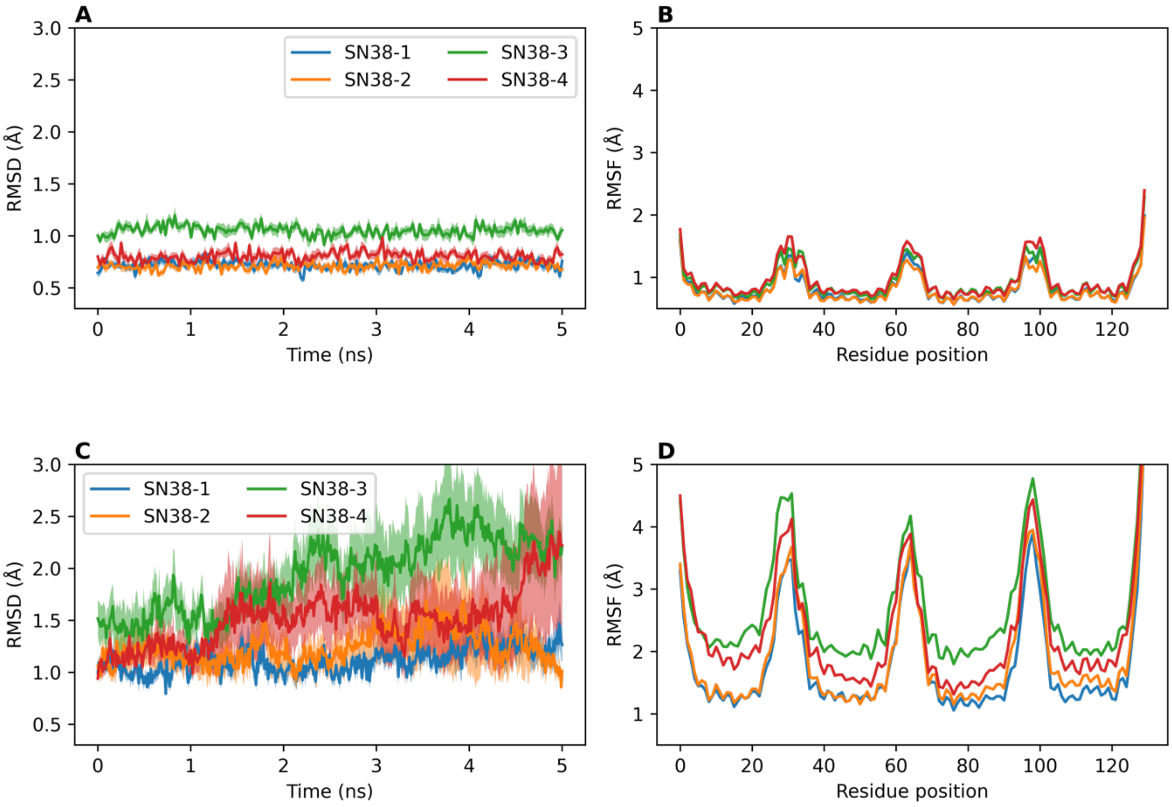


**Supplementary Figure 10. Selected SN38 binder binding-site backbone RMSD and Cα RMSF throughout simulations with (A&B) and without (C&D) the 20 cal mol$^{-1}$Å$^{-2}$ restraint.** Weak backbone restraints allow accelerated sampling of the binding site and ligand motion while preventing the structure from undergoing large conformational changes at elevated temperatures [1].

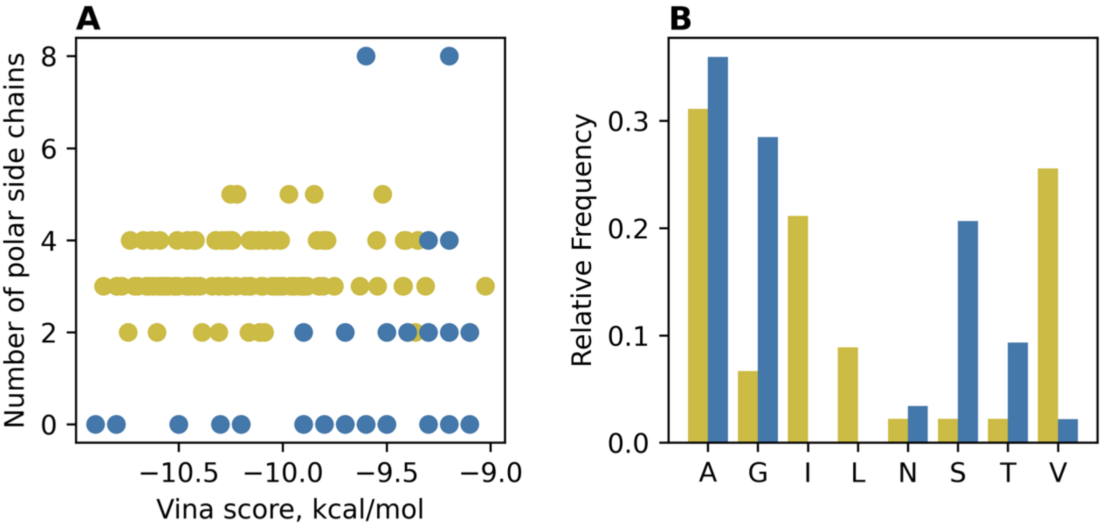


**Supplementary Figure 11. Enrichment of polar side chains for SN38 (the most polar ligand, yellow) compared to the DPH (most hydrophobic ligand, blue). (A)** AutoDock Vina scores for sequences with varying number of polar side chains. For DPH, the best scoring sequences had no polar side chains. For SN38, sequences with binding sites with 2 – 4 polar side chains returned the best scores. **(B)** Frequency of binding-site residues in sequences with <-9.5 kcal/mol Vina score. In both cases, 10 positions were designed in the 4-helix bundle, and the amino acid pallet was limited to AGILNSTV.

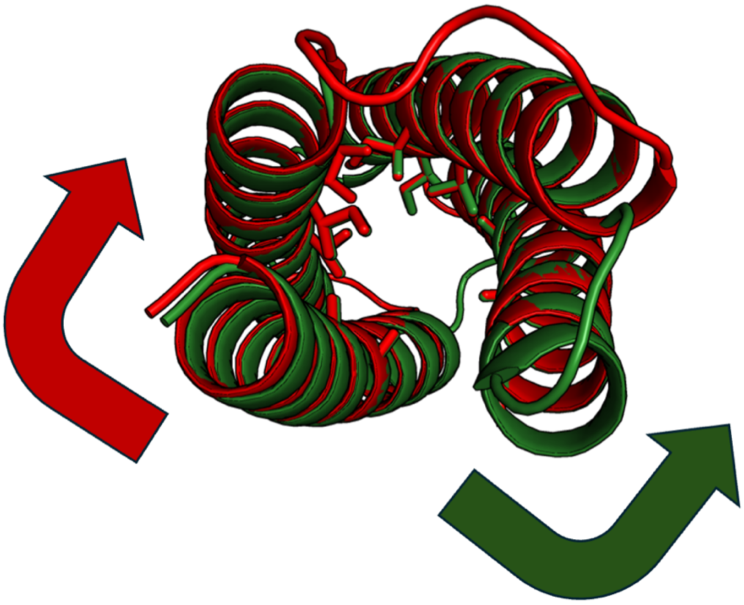

**Supplementary Figure 12. AlphaFold2 predicted topology swap by binding site rotation, sc-apCC-4-DPH-2 (clockwise, red) and sc-apCC-4-DPH-1 (anticlockwise, green, same as parent 8a3k).** The direction of the topology is shown by arrows and binding site side chains shown as sticks. Despite the global structural change (backbone RMSD = 10.3 Å), the binding site is retained (heavy atom RMSD = 0.2 Å). The topology swap restores optimal knobs-into-holes packing by placing all **Ile** at **d** within the same side of the protein.

#### 2.2 Purification and characterisation of 4-helix binders designed with RASSCCoL

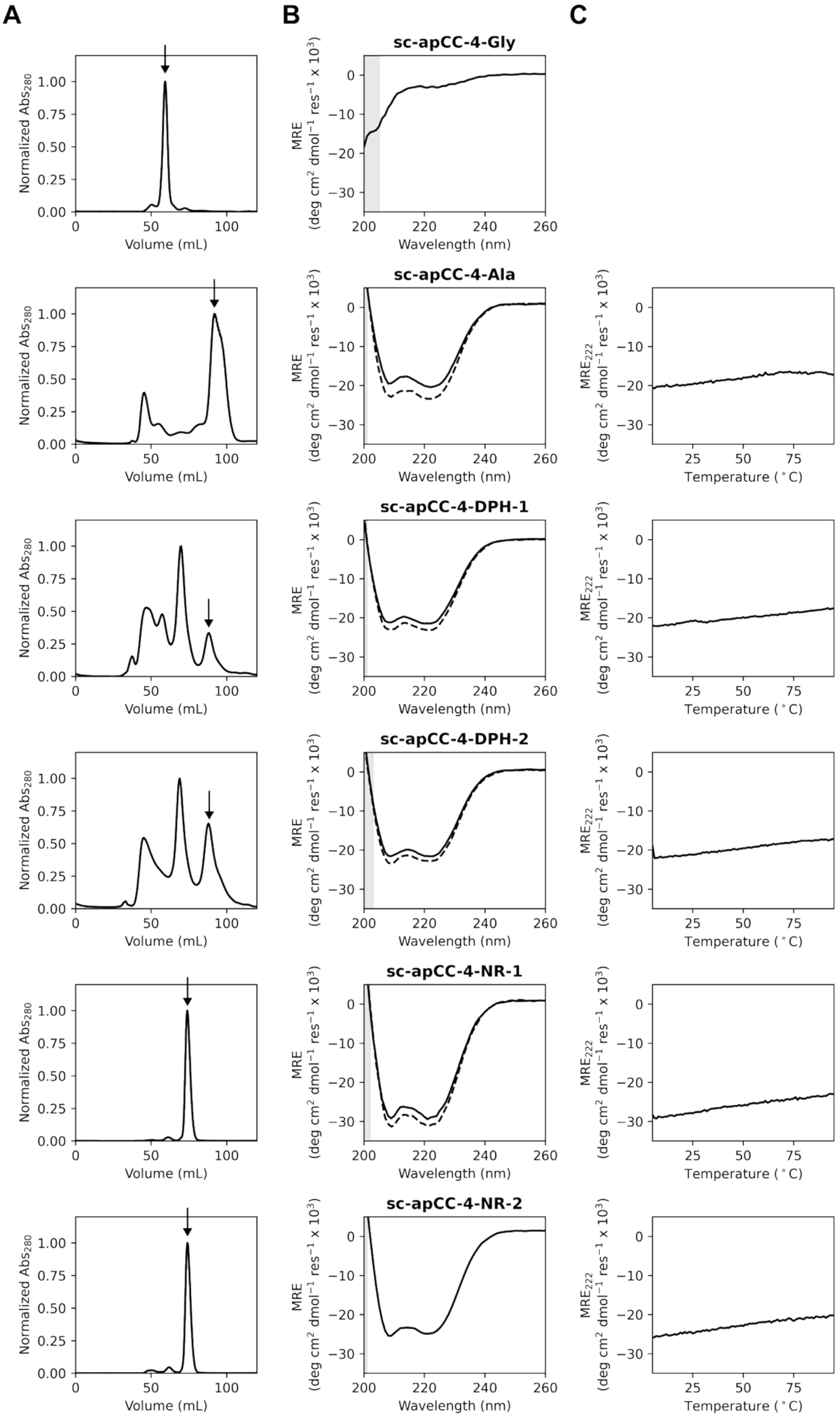

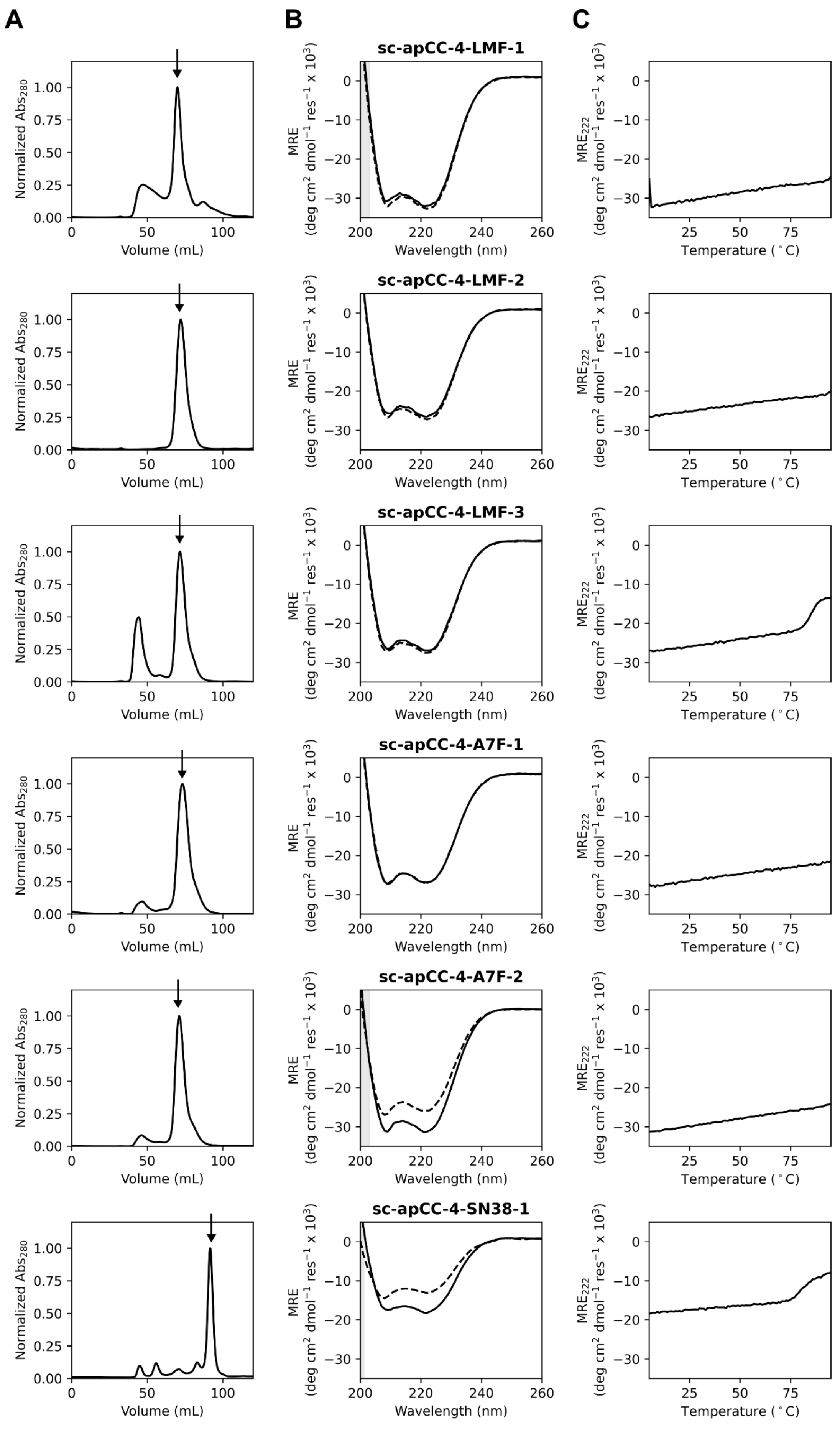
A
B
C
sc-apCC-4-LMF-1
sc-apCC-4-LMF-2
sc-apCC-4-LMF-3
sc-apCC-4-A7F-1
sc-apCC-4-A7F-2
sc-apCC-4-SN38-1
Normalized Abs280
Volume (mL)
MRE (deg cm2 dmol−1 res−1 x 103)
Wavelength (nm)
MRE222 (deg cm2 dmol−1 res−1 x 103)
Temperature (°C)

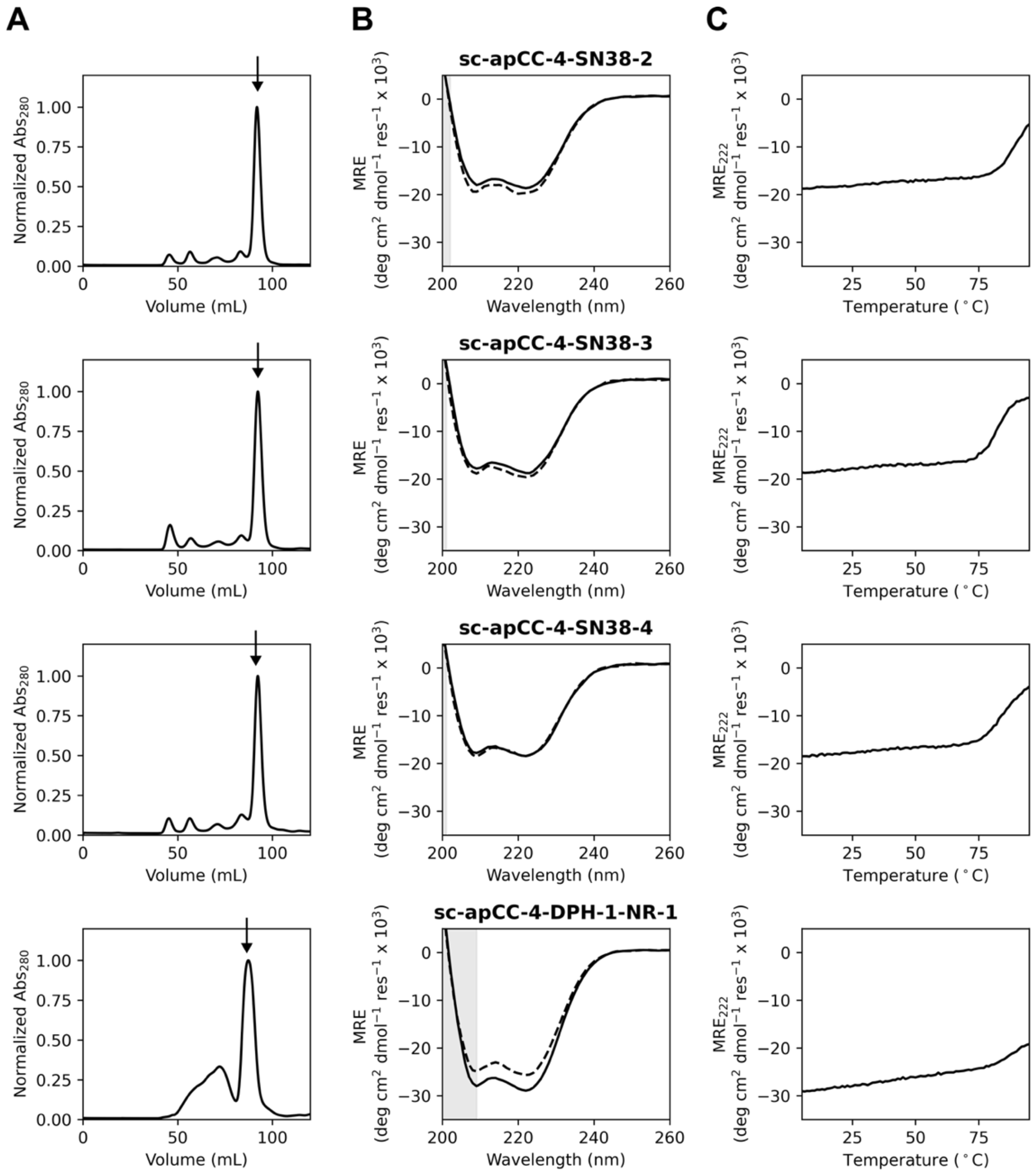


**Supplementary Figure 13. Biophysical characterization of apo sc-apCC-4 variants designed with RASSCCoL. (A)** Preparative size-exclusion chromatography (SEC) elution profiles. Proteins were purified on either Superdex 75 pg or Superdex 200 pg columns in 50 mM sodium phosphate, pH 7.4, with 150 mM NaCl. The arrow marks the elution fractions selected for further analysis. **(B)** Far-UV circular dichroism (CD) spectral scans collected at 5 °C (solid lines) and after heating to 95 °C followed by cooling to 5 °C (dashed lines). Filled grey regions indicate HT voltage ≥600 V. **Conditions**: 10 µM protein in 50 mM sodium phosphate, 150 mM NaCl, pH 7.4. **(C)** Temperature-dependent CD monitored at 222 nm. Thermal melts were recorded on heating (solid lines) and on cooling (dashed lines) to/from 95 °C. **Conditions**: 10 µM protein in 50 mM sodium phosphate, 150 mM NaCl, pH 7.4.

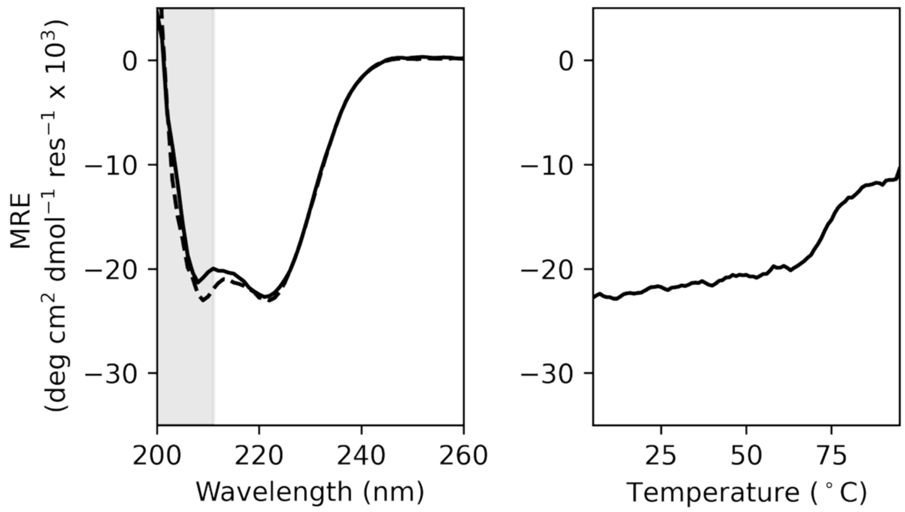


**Supplementary Figure 14. Biophysical characterisation of sc-apCC-4-SN38-1 at pH 4. (A)** Far-UV circular dichroism (CD) spectral scans collected at 5 °C (solid lines) and after heating to 95 °C followed by cooling to 5 °C (dashed lines). Filled grey regions indicate HT voltage ≥600 V. **(B)** Temperature-dependent CD monitored at 222 nm. Thermal melts were recorded on heating (solid lines) and on cooling (dashed lines) to/from 95 °C. **Conditions**: 10 µM protein in 100 mM sodium acetate, 150 mM NaCl, pH 4.

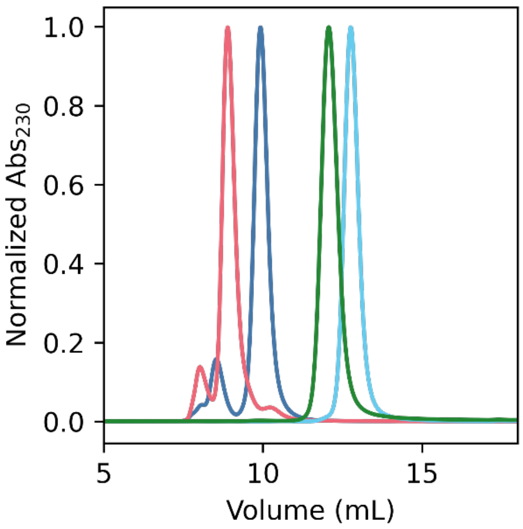


**Supplementary Figure 15. Calibration curve used for aFPLC analysis.** Conalbumin (76kDa); red. Ovalbumin (44 kDa); blue. Sc-apCC-6-SLLA (26 kDa, PDB id: 8QAE), green. Sc-apCC-4 (17 kDa, PDB id: 8A3K); cyan. **Conditions:** 50 mM sodium phosphate, pH 7.4, 150 mM NaCl. Proteins run on Superdex 75 Increase 10/300 GL at 0.5 – 2 mg/ml.

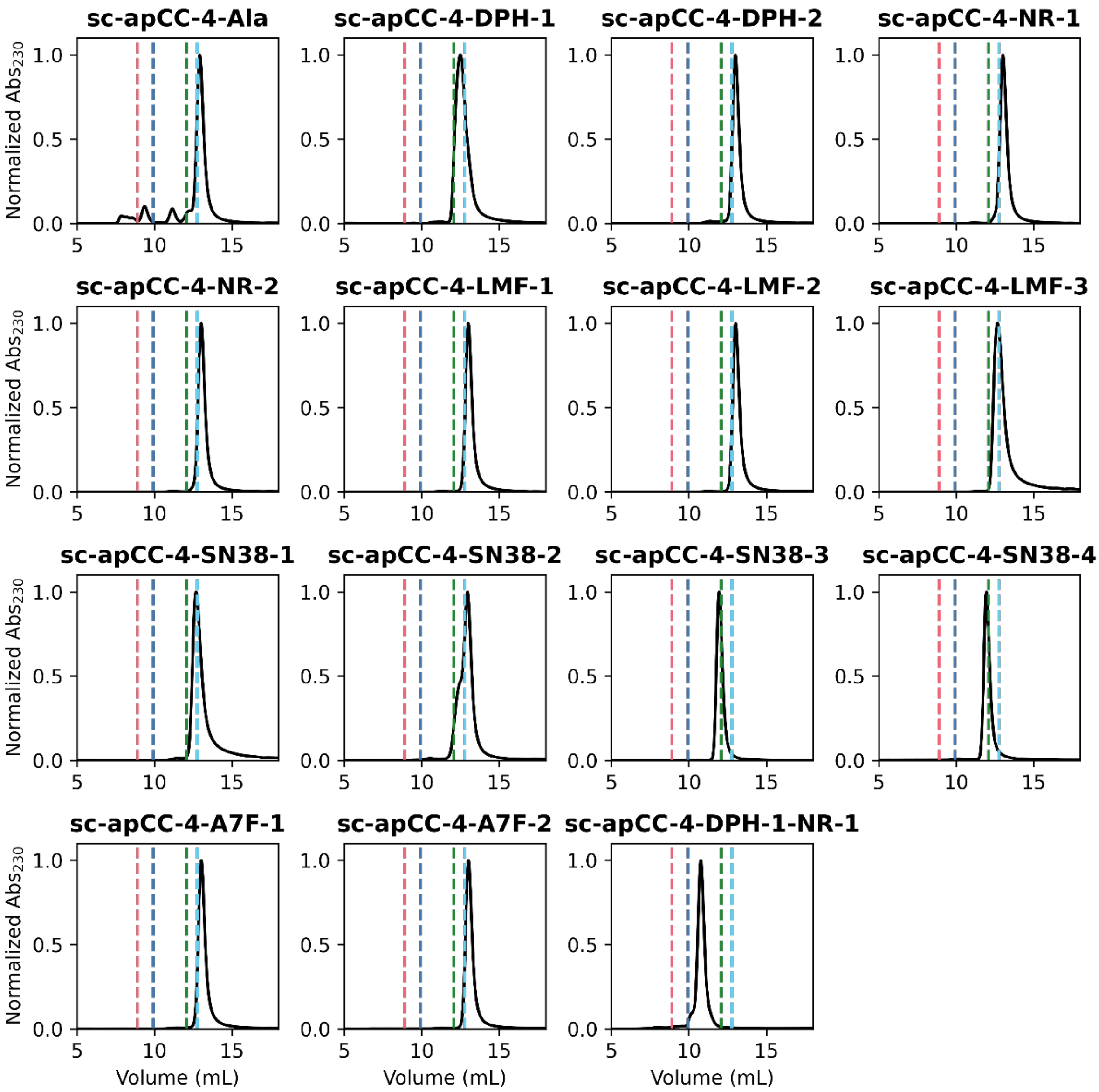


**Supplementary Figure 16. aFPLC traces of all 4-helix bundle proteins in this study.** Dashed lines represent elution volumes of calibration standards (Figure S15). **Conditions:** 50 mM sodium phosphate, pH 7.4, 150 mM NaCl. Proteins run on Superdex 75 Increase 10/300 GL at 0.5 – 2 mg/ml.

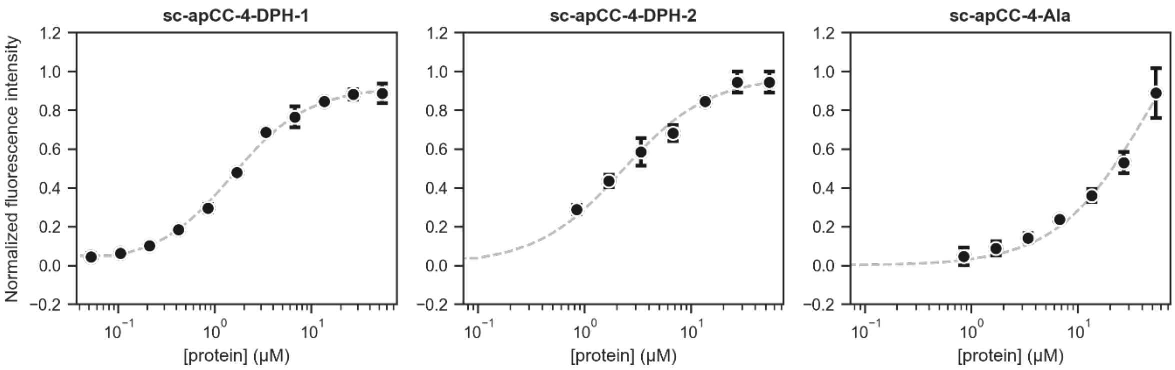


**Supplementary Figure 17. Saturation binding curves for 4-helix bundle DPH binders designed with RASSCCoL.** Fits returned $K_{D(\text{sc-apCC-4-DPH-1})}$ = 1.3 ± 0.1 μM, $K_{D(\text{sc-apCC4-DPH-2})}$ = 2.0 ± 0.2 μM, $K_{D(\text{sc-apCC-4-Ala})}$ > 50 μM**. Conditions**: 0.5 μM ligand, 0 – 60 μM protein, data are the mean of three independent repeats, error bars represent the standard deviation from the mean. All data collected in PBS, 10% v/v MeCN, pH 7.4.

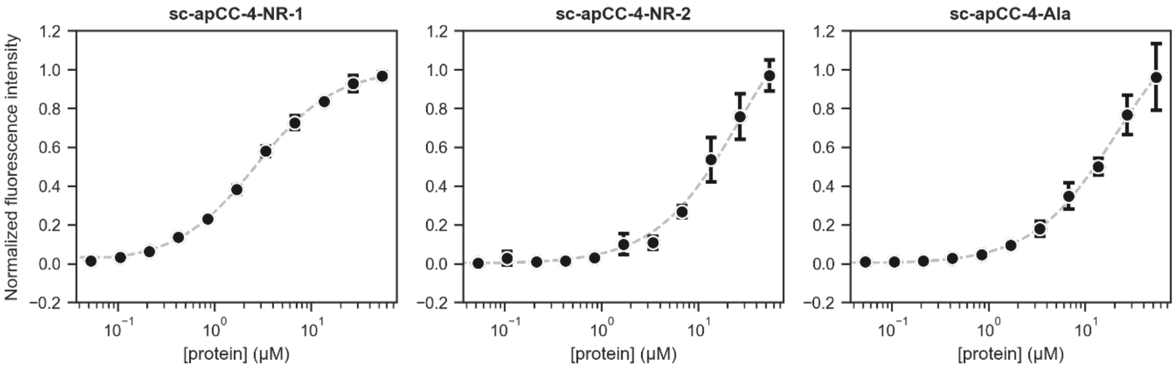


**Supplementary Figure 18. Saturation binding curves for 4-helix bundle Nile red binders designed with RASSCCoL.** Fits returned $K_{D(\text{sc-apCC-4-NR-1})}$ = 2.4 ± 0.1 μM, $K_{D(\text{sc-apCC-4-NR-2})}$ > 25 μM, $K_{D(\text{sc-apCC-4-Ala})}$ > 20 μM**. Conditions**: 0.5 μM ligand, 0 – 60 μM protein, data are the mean of three independent repeats, error bars represent the standard deviation from the mean. All data collected in PBS, 10% v/v MeCN, pH 7.4.

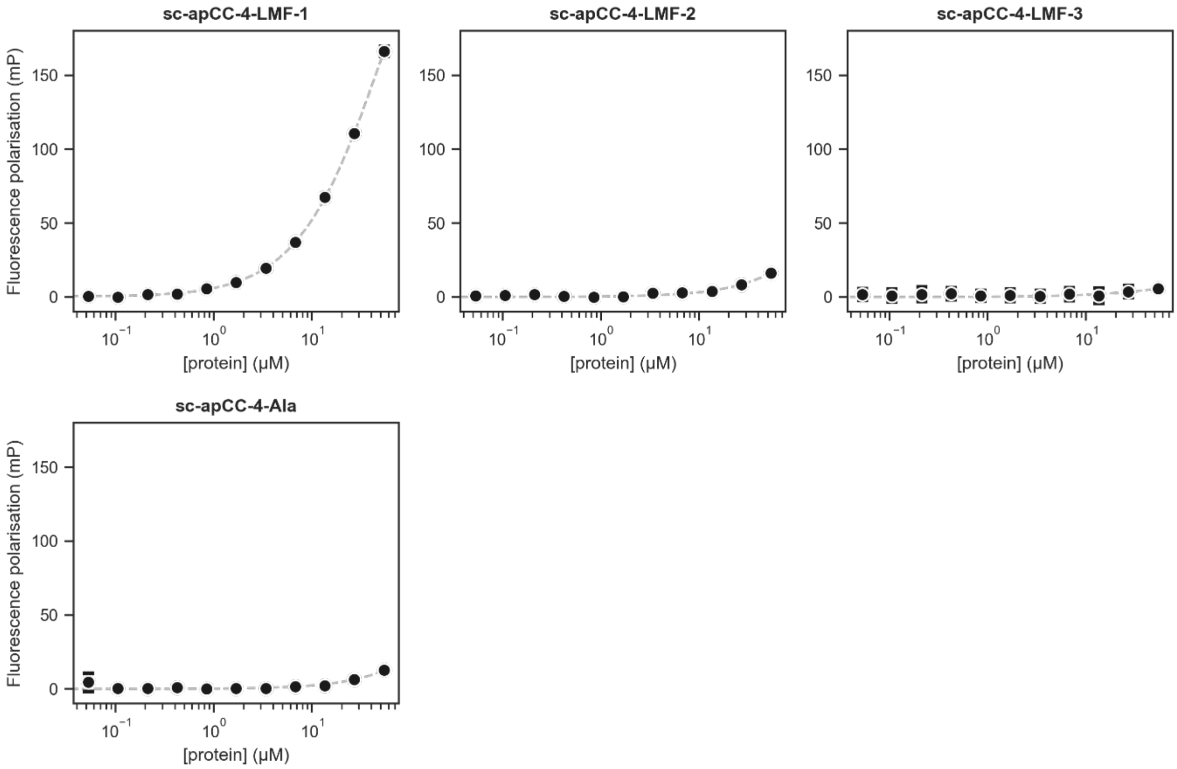


**Supplementary Figure 19. Saturation binding curves for 4-helix bundle lumiflavin binders designed with RASSCCoL.** Fit returned $K_{D(sc\text{-}apCC\text{-}4\text{-}LMF\text{-}1)}$ > 50 μM, sc-apCC-4-LMF-2, sca-apCC-4-LMF-3 and sc-apCC-4-Ala showed no binding. **Conditions**: 1 μM ligand, 0 – 60 μM protein, data are the mean of three independent repeats, error bars represent the standard deviation from the mean. All data collected in PBS, 10% v/v DMSO, pH 7.4.

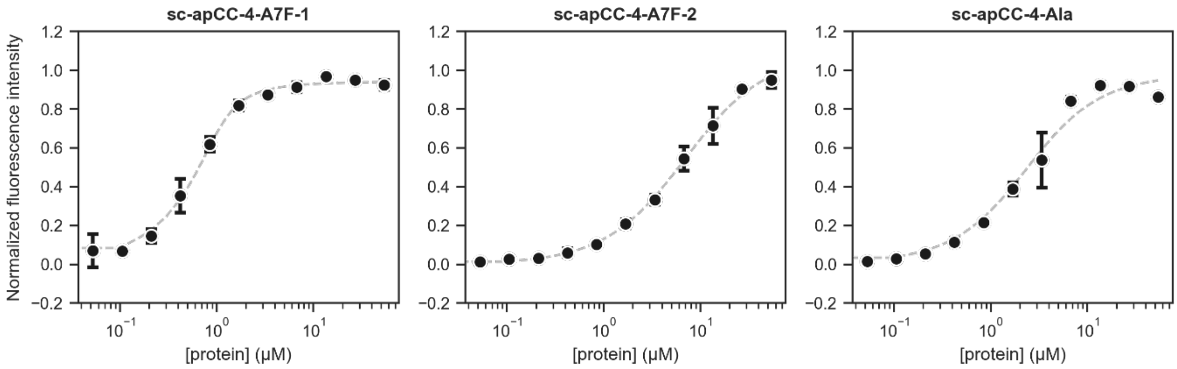


**Supplementary Figure 20. Saturation binding curves for 4-helix bundle arcyriaflavin A binders designed with RASSCCoL.** Fits returned $K_{D(sc\text{-}apCC\text{-}4\text{-}A7F\text{-}1)}$ = 0.11 ± 0.02 μM, $K_{D(sc\text{-}apCC\text{-}4\text{-}A7F\text{-}2)}$ = 6.5 ± 0.3 μM, $K_{D(sc\text{-}apCC\text{-}4\text{-}Ala)}$ = 1.8 ± 0.3 μM. **Conditions**: 1 μM ligand, 0 – 60 μM protein, data are the mean of three independent repeats, error bars represent the standard deviation from the mean. All data collected in PBS, 10% v/v MeCN, pH 7.4.

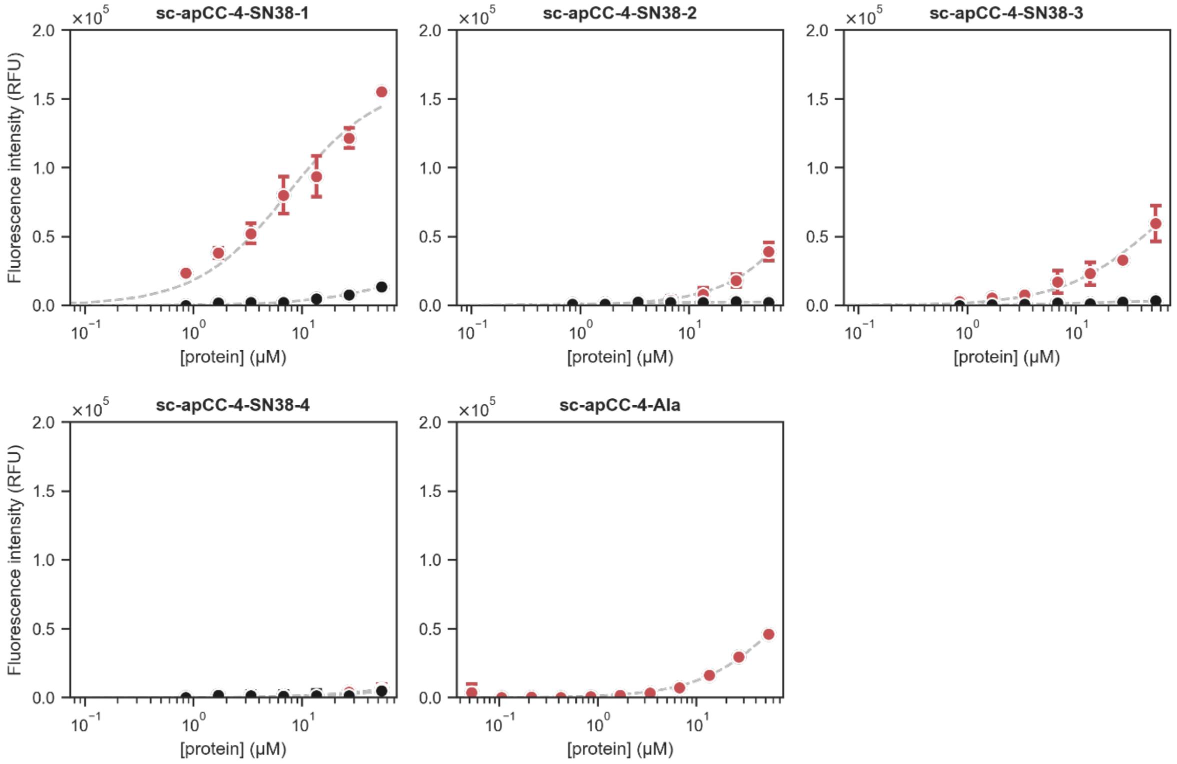


**Supplementary Figure 21. Saturation binding curves for 4-helix bundle SN38 binders designed with RASSCCoL; measured at pH 7.4 (black) or pH 4 (red).** At pH ≤ 4.5, SN38 lactone ring is stable (our design target), resulting in stronger binding, whereas at pH 7.4 the ring undergoes reversible opening. Fits for pH 4 returned $K_{D(\text{sc-apCC-4-SN38-1})}$ > 5 μM. **Conditions**: 1 μM ligand, 0 – 60 μM protein, data are the mean of three independent repeats, error bars represent the standard deviation from the mean. All data collected either in ABS, 5% v/v DMSO, pH 4 (red) or PBS, 5% v/v DMSO, pH 7.4 (black).

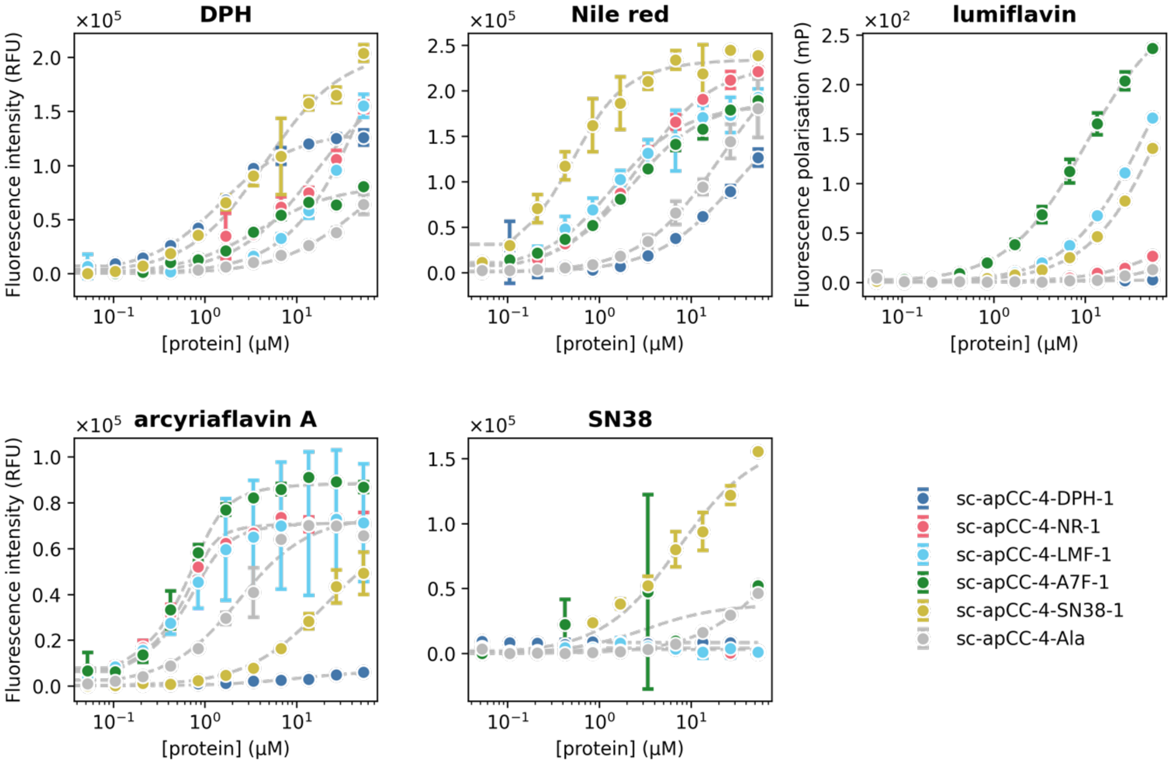


**Supplementary Figure 22. Saturation binding curves showing ligand selectivity for the highest-affinity designs in sc-apCC-4.** Conditions: 0.5 μM DPH/Nile red or 1 μM lumiflavin/arcyriaflavinA/SN38, 0 – 60 μM protein, data are the mean of three independent repeats, error bars represent the standard deviation from the mean. All data were collected in PBS, pH 7.4 with either 10% v/v MeCN (DPH/Nile red/aryriaflavin A) or 10% v/v DMSO (lumiflavin/SN38). See Supplementary Table 5 for fitted binding affinities.

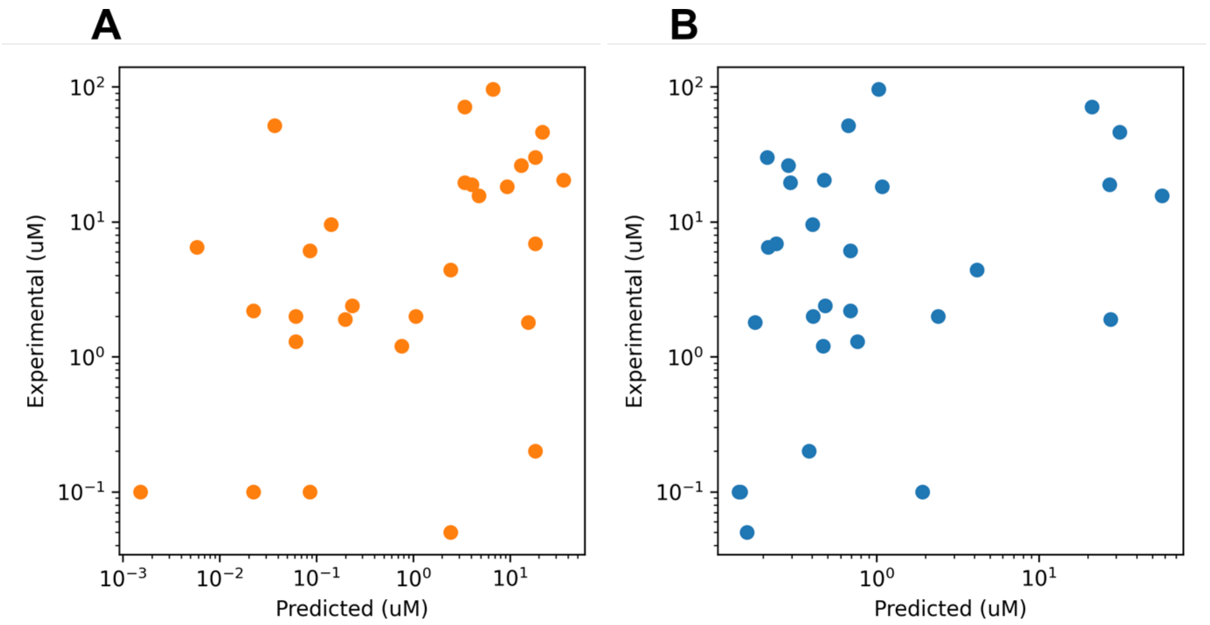


**Supplementary Figure 23. Correlation of experimental and predicted binding constants for all RASSCCoL designs.** **(A)** Affinities calculated from AutoDock Vina docking scores using the AlphaFold2 models for all experimentally validated ligand–binder pairs. **(B)** Predicted affinities calculated from Boltz-2 structure predictions with the ligand. Since both in silico methods produce binding energies, predicted affinities were derived using the relationship $\Delta G = RT \ln K_D$.

#### 2.3 Purification and characterisation of 4-helix binders refined with LigandMPNN and Boltz2

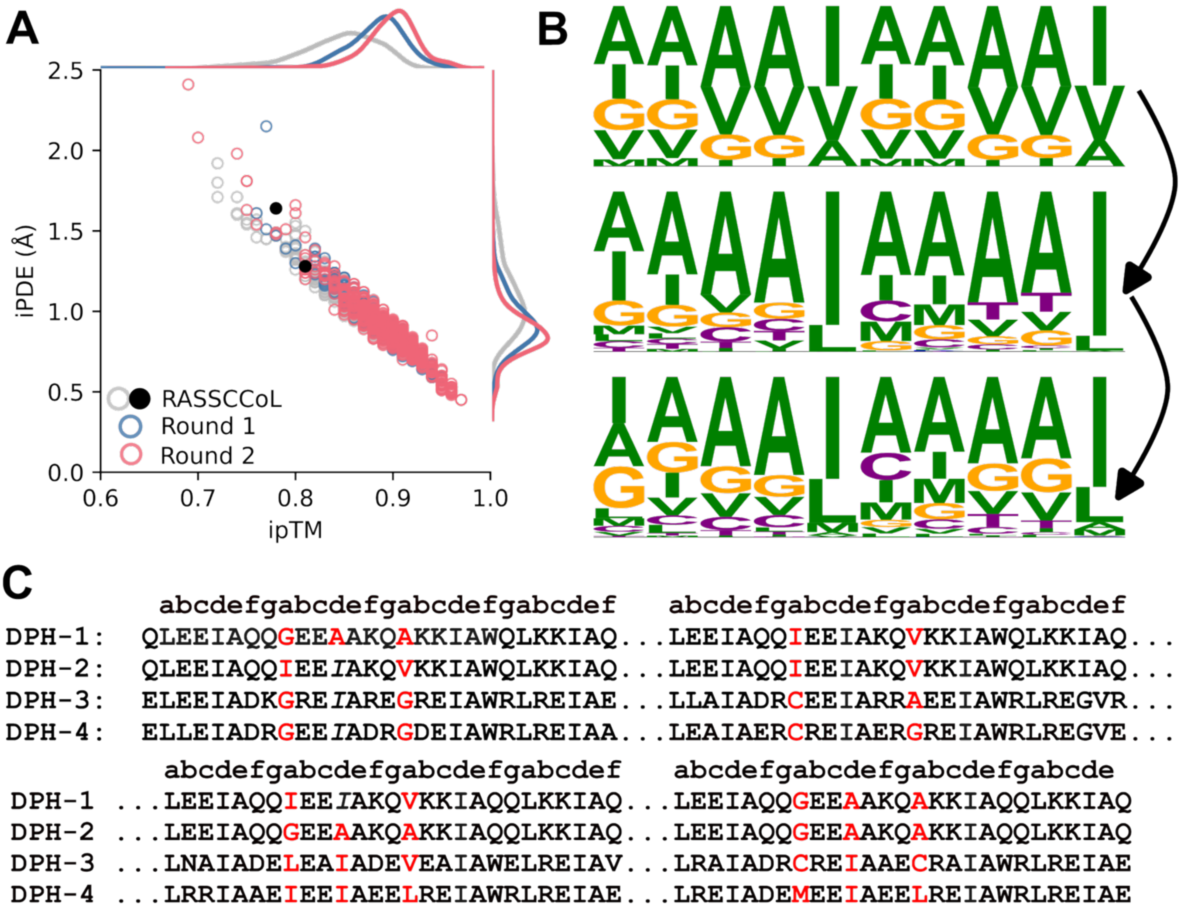


**Supplementary Figure 24. RASSCCoL hit optimisation using Boltz2 and LigandMPNN for DPH binders.** Change in Boltz2 confidence scores following two rounds of redesign with LigandMPNN. Black circles indicate experimentally characterized RASSCCoL designs (Figures S3). **(B)** Sequence logo comparison of the top 1000 designs ranked by ipTM relative to the original RASSCCoL sequences (top panel). 270 RASSCCoL sequences with the lowest AutoDock Vina scores were selected for redesign using LigandMPNN (default settings) and subsequently refolded with Boltz2. The top 1000 sequences ranked by ipTM were then subjected to a second round of redesign and refolding. After two design iterations, the ipTM scores reached ~0.97, and the sequence space has converged. **(C)** Sequence alignments of RASSCCoL designs (DPH-1/2) and LigandMPNN designs (DPH-3/4). Binding-site residues (shown in panel **B**) are highlighted in red. Loop regions are omitted for clarity, and coiled-coil heptad positions are labelled a–g. Because DPH-1 and DPH-2/3/4 were designed with opposite handedness, position 9 in DPH-1 (italicised) corresponds to position 80 in DPH-2/3/4. The maximum sequence identity between RASSCCoL and LigandMPNN designs is 41%.

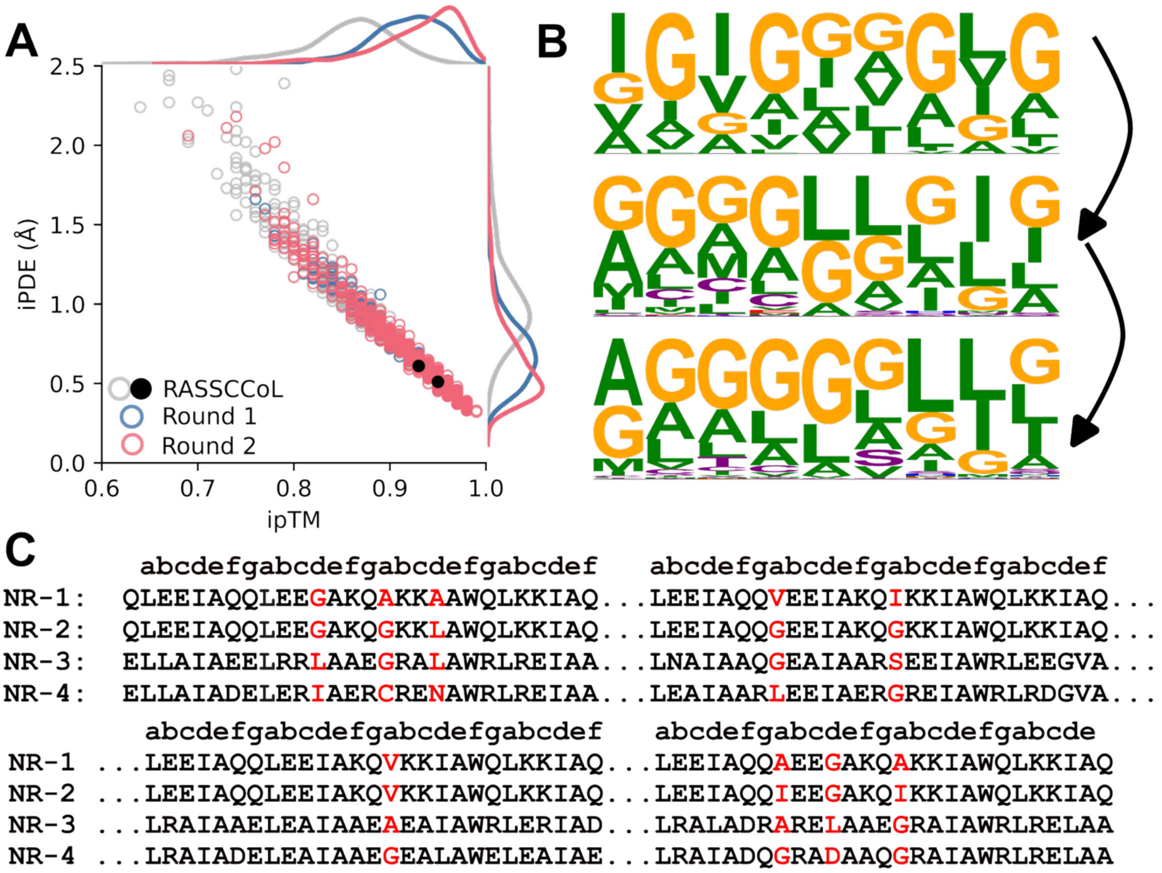


**Supplementary Figure 25. RASSCCoL hit optimisation using Boltz2 and LigandMPNN for NR binders.** Change in Boltz2 confidence scores following two rounds of redesign with LigandMPNN. Black circles indicate experimentally characterized RASSCCoL designs (Figures S4). **(B)** Sequence logo comparison of the top 1000 designs ranked by ipTM relative to the original RASSCCoL sequences (top panel). 600 RASSCCoL sequences with the lowest AutoDock Vina scores were selected for redesign using LigandMPNN (default settings, 5 sequences) and subsequently refolded with Boltz2. The top 1000 sequences ranked by ipTM were then subjected to a second round of redesign and refolding. After two design iterations, the ipTM scores reached ~0.99, and the sequence space has converged. **(C)** Sequence alignments of RASSCCoL designs (NR-1/2) and LigandMPNN designs (NR-3/4). Binding-site residues (shown in panel **B**) are highlighted in red. Loop regions are omitted for clarity, and coiled-coil heptad positions are labelled a–g. The maximum sequence identity between RASSCCoL and LigandMPNN designs is 40%.

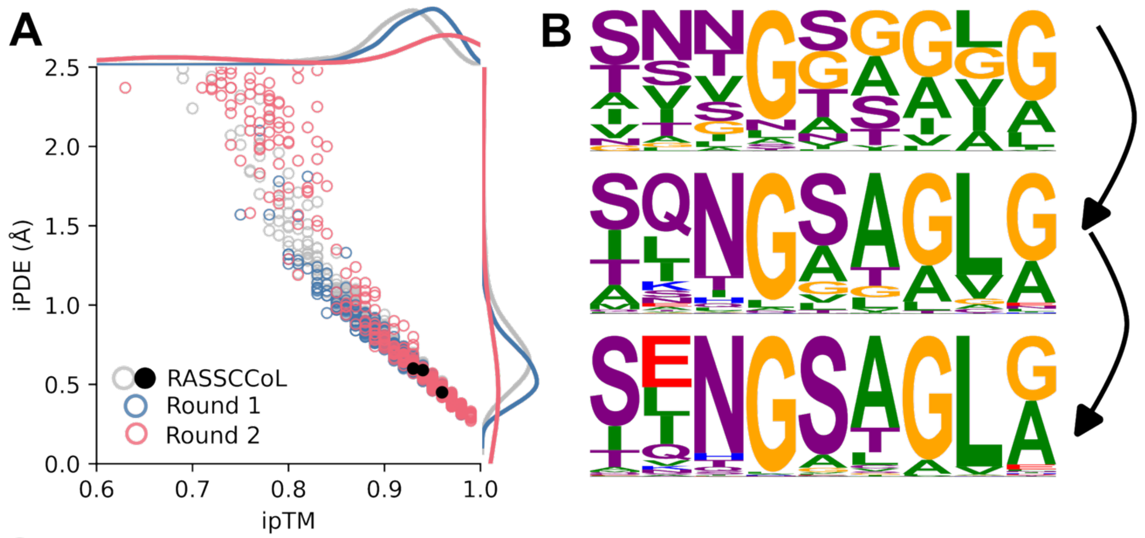


**C**

```
        abcdefgabcdefgabcdefgabcdef   abcdefgabcdefgabcdefgabcdef
LMF-1:  QLEEIAQQSEEGAKQGKKIAWQLKKIAQ...LEEIAQQLEEIAKQSKKIAWQLKKIAQ...
LMF-2:  QLEEIAQQTEEGAKQAKKIAWQLKKIAQ...LEEIAQQLEEIAKQSKKIAWQLKKIAQ...
LMF-3:  QLEEIAQQSEEGAKQAKKIAWQLKKIAQ...LEEIAQQLEEIAKQSKKIAWQLKKIAQ...
LMF-4:  ELREIAEETREGAREGREIAWELRRIAE...LEAIADELEAIAARSEEIAWRLRAGVE...
LMF-5:  ELREIAEETREGAREGREIAWELRRIAE...LLAIADRLEEIARESERIAWRILEGVE...

         abcdefgabcdefgabcdefgabcdef   abcdefgabcdefgabcdefgabcde
LMF-1 ...LEEIAQQVEEIAKQLKKIAWQLKKIAQ...LEEIAQQLEEGAKQAKKTAWQLKKIAQ
LMF-2 ...LEEIAQQTEEIAKQVKKIAWQLKKIAQ...LEEIAQQLEEGAKQAKKTAWQLKKIAQ
LMF-3 ...LEEIAQQNEEIAKQVKKIAWQLKKIAQ...LEEIAQQLEEGAKQSKKTAWQLKKIAQ
LMF-4 ...LRAIADEIDAIADRLEAIAWRLEEIAA...LRAIAAELRAEAAAARANAWKLRELAA
LMF-5:...LREIADRQEAIAEELRRIAWELREIAE...LREIADRLEEAAERTRENAWRLRRLAE
```

**Supplementary Figure 26. RASSCCoL hit optimisation using Boltz2 and LigandMPNN for LMF binders.** Change in Boltz2 confidence scores following two rounds of redesign with LigandMPNN. Black circles indicate experimentally characterized RASSCCoL designs (Figures S5). **(B)** Sequence logo comparison of the top 1000 designs ranked by ipTM relative to the original RASSCCoL sequences (top panel). 1000 RASSCCoL sequences with the lowest AutoDock Vina scores were selected for redesign using LigandMPNN (default settings) and subsequently refolded with Boltz2. The top 1000 sequences ranked by ipTM were then subjected to a second round of redesign and refolding. After two design iterations, the ipTM scores reached ~0.99, and the sequence space has converged. **(C)** Sequence alignments of RASSCCoL designs (LMF-1/2/3) and LigandMPNN designs (LMF-4/5). Binding-site residues (shown in panel **B**) are highlighted in red. Loop regions are omitted for clarity, and coiled-coil heptad positions are labelled a–g. The maximum sequence identity between RASSCCoL and LigandMPNN designs is 44%.

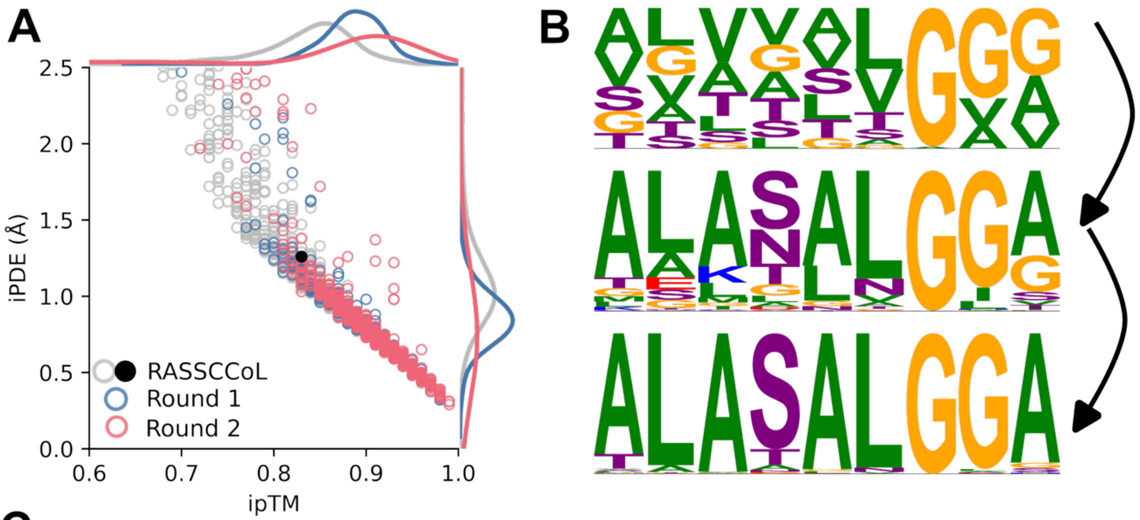


**C**

```
           abcdefgabcdefgabcdefgabcdef   abcdefgabcdefgabcdefgabcdef
A7F-1:   QLEEIAQQLEEGAKQTKKVAWQLKKIAQ...LEEIAQQVEEIAKQLKKIAWQLKKIAQ...
A7F-2:   QLEEIAQQLEEGAKQAKKGAWQLKKIAQ...LEEIAQQGEEIAKQLKKIAWQLKKIAQ...
A7F-3:   ELLAIADELRAGAWETRAAADRLREIAA...LEAIADQLRAIAWRLEAIADRLRAGVA...
A7F-4:   ELAAIADELEAGADAAEAAAWELERIAA...LRAIADELRAQAWQLRAIAARLRAGVE...
            abcdefgabcdefgabcdefgabcdef   abcdefgabcdefgabcdefgabcde
A7F-1  ...LEEIAQQLEEIAKQSKKIAWQLKKIAQ...LEEIAQQGEEGAKQAKKIAWQLKKIAQ
A7F-2  ...LEEIAQQLEEIAKQVKKIAWQLKKIAQ...LEEIAQQTEEGAKQLKKIAWQLKKIAQ
A7F-3  ...LRAIADELRAIAAESRAIADRLERIAD...LRAIADGARAGAARAEEIAARLEELAA
A7F-4  ...LRAIADELDALADAERATAARLREIAE...LRAIAAGAEAGAARARAIAARLRALAA
```

**Supplementary Figure 27. RASSCCoL hit optimisation using Boltz2 and LigandMPNN for arcyriaflavin A binders.** Change in Boltz2 confidence scores following two rounds of redesign with LigandMPNN. Black circles indicate experimentally characterized RASSCCoL designs (Figure S6). **(B)** Sequence logo comparison of the top 1000 designs ranked by ipTM relative to the original RASSCCoL sequences (top panel). 1000 RASSCCoL sequences with the lowest AutoDock Vina scores were selected for redesign using LigandMPNN (default settings) and subsequently refolded with Boltz2. The top 1000 sequences ranked by ipTM were then subjected to a second round of redesign and refolding. After two design iterations, the ipTM scores reached ~0.99, and the sequence space has converged. **(C)** Sequence alignments of RASSCCoL designs (A7F-1/2) and LigandMPNN designs (A7F-3/4). Binding-site residues (shown in panel **B**) are highlighted in red. Loop regions are omitted for clarity, and coiled-coil heptad positions are labelled a–g. The maximum sequence identity between RASSCCoL and LigandMPNN designs is 35%.

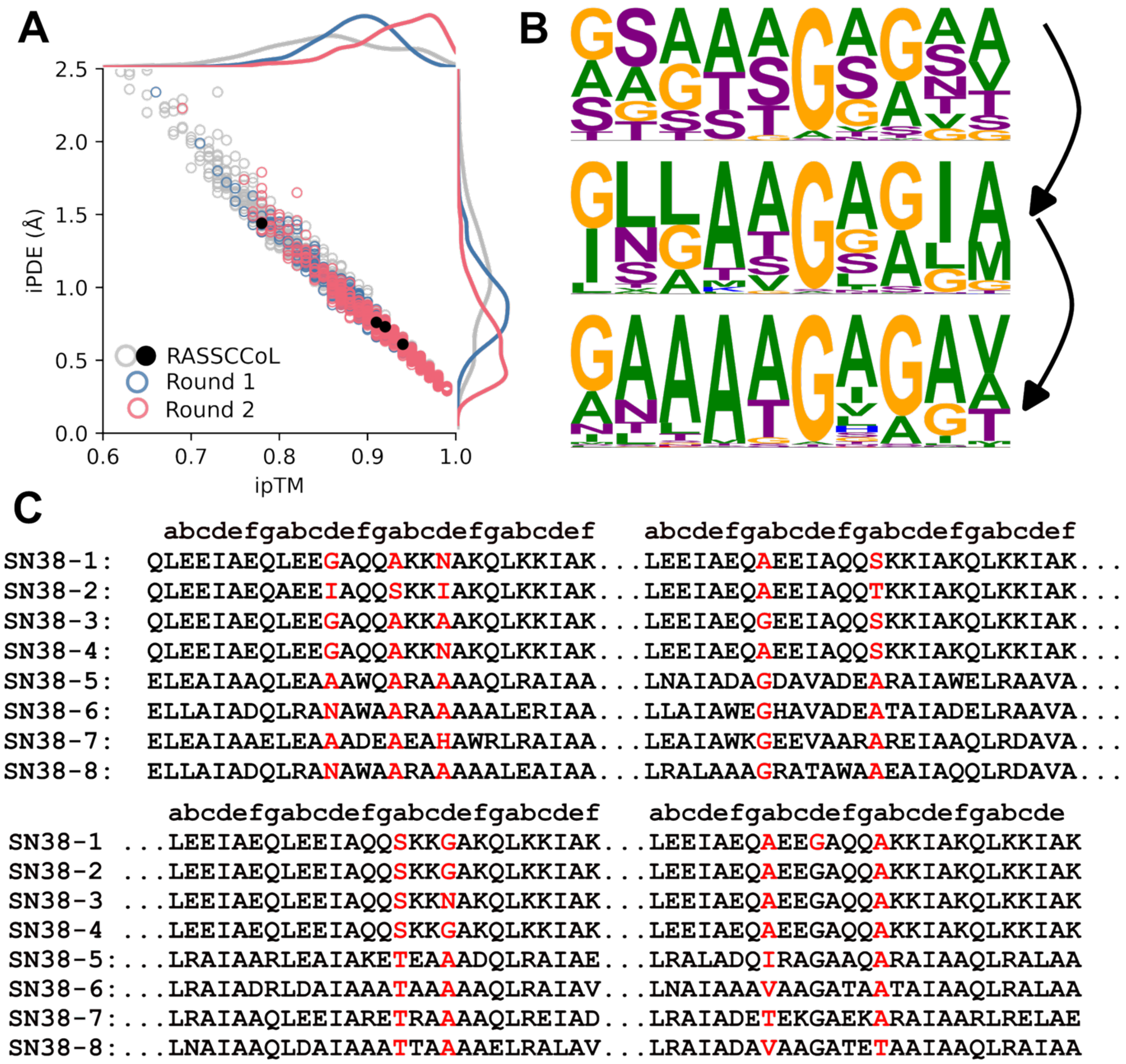


**Supplementary Figure 28. RASSCCoL hit optimisation using Boltz2 and LigandMPNN for SN38 binders. (A)** Change in Boltz2 confidence scores following two rounds of redesign with LigandMPNN. Black circles indicate experimentally characterized RASSCCoL designs (Figures S7–S9). **(B)** Sequence logo comparison of the top 1000 designs ranked by ipTM relative to the original RASSCCoL sequences (top panel). 1000 RASSCCoL sequences with the lowest AutoDock Vina scores were selected for redesign using LigandMPNN (default settings) and subsequently refolded with Boltz2. The top 1000 sequences ranked by ipTM were then subjected to a second round of redesign and refolding. After two design iterations, the ipTM scores reached ~0.99, and the sequence space has converged. **(C)** Sequence alignments of RASSCCoL designs (SN38-1/2/3/4) and LigandMPNN designs (SN38-5/6/7/8). Binding-site residues (shown in panel **B**) are highlighted in red. Loop regions are omitted for clarity, and coiled-coil heptad positions are labelled a–g. The maximum sequence identity between RASSCCoL and LigandMPNN designs is 43%.

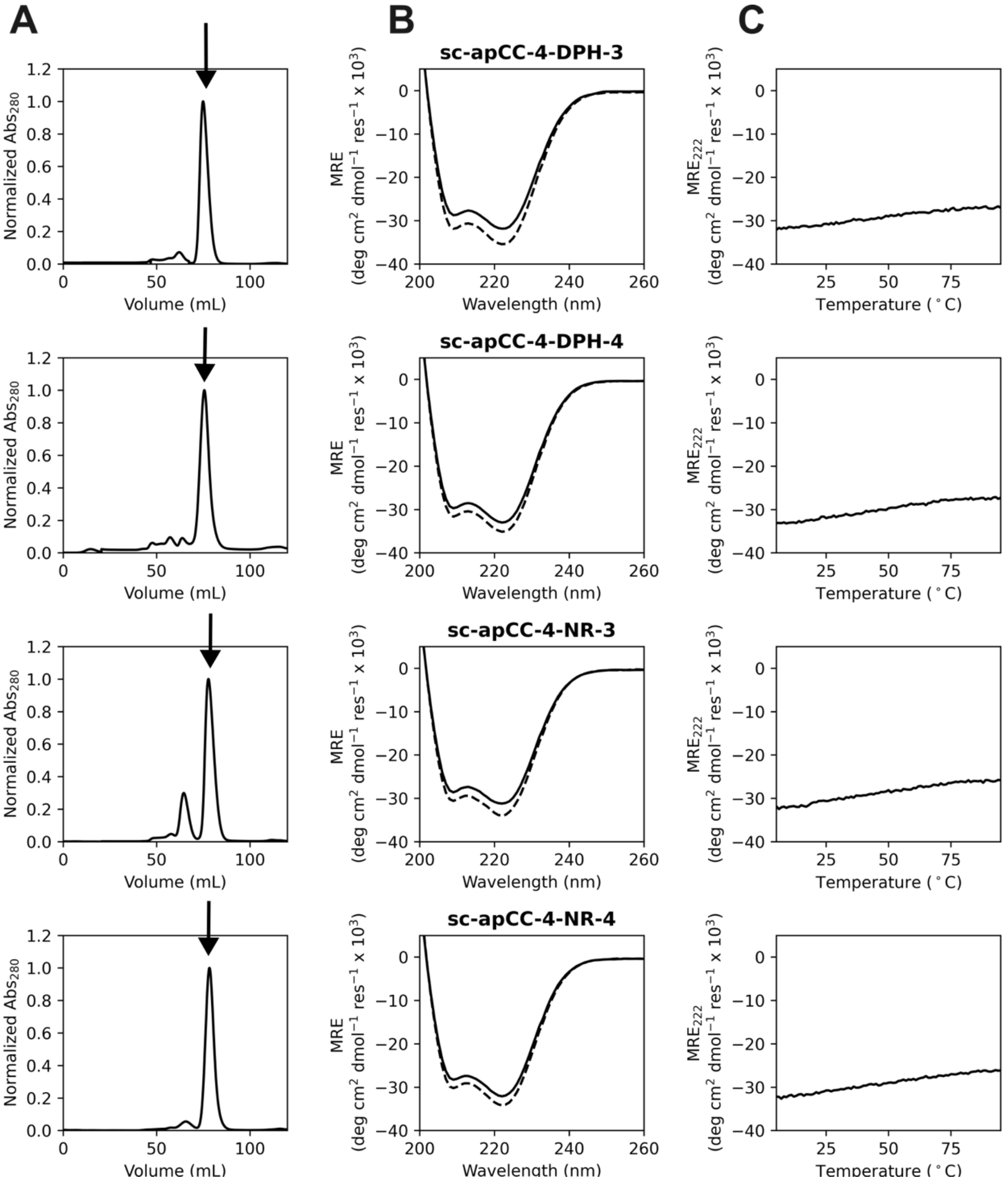

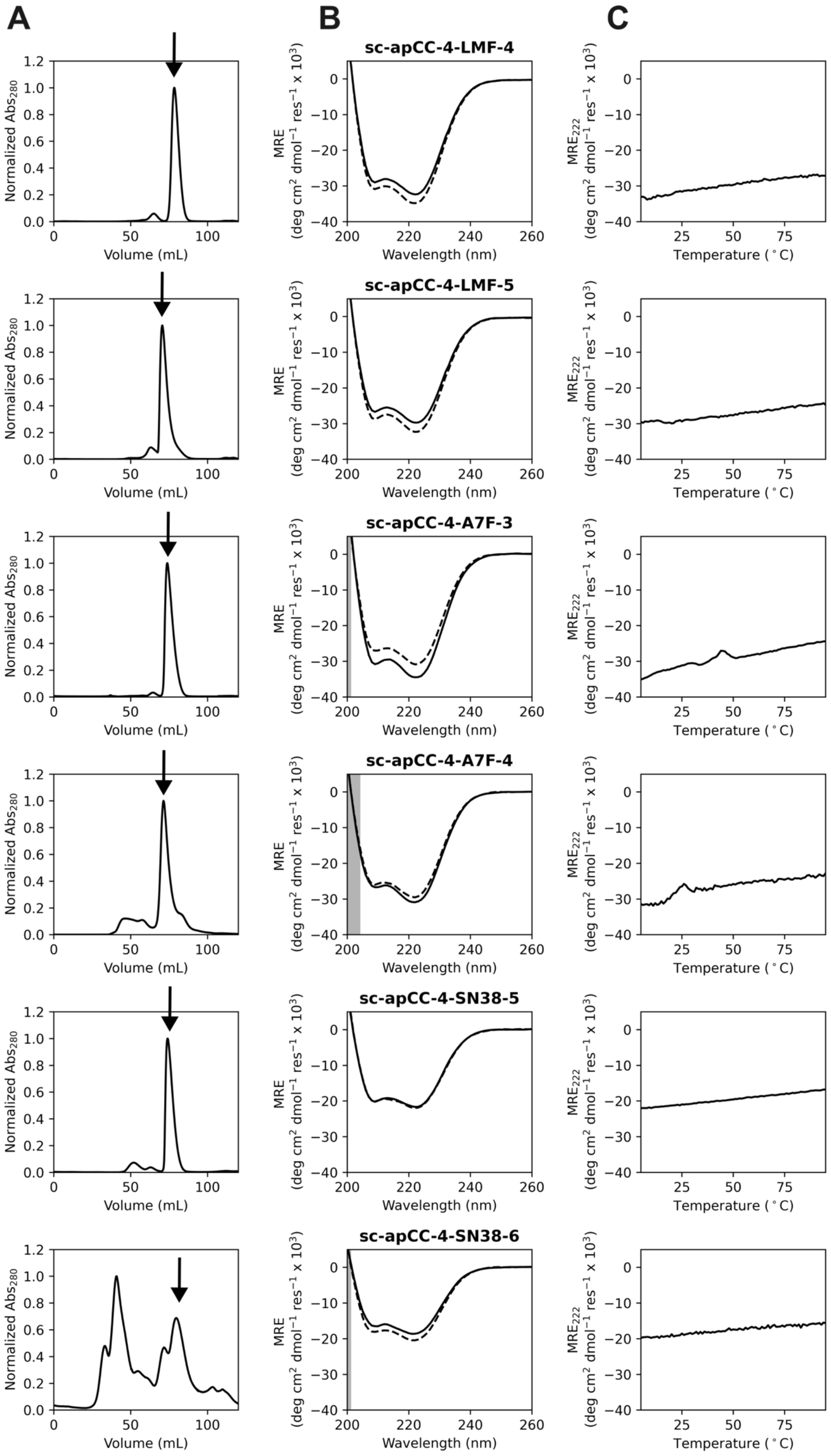
A
B
C
sc-apCC-4-LMF-4
sc-apCC-4-LMF-5
sc-apCC-4-A7F-3
sc-apCC-4-A7F-4
sc-apCC-4-SN38-5
sc-apCC-4-SN38-6
Normalized Abs280
Volume (mL)
MRE (deg cm2 dmol−1 res−1 x 103)
Wavelength (nm)
MRE222 (deg cm2 dmol−1 res−1 x 103)
Temperature (°C)

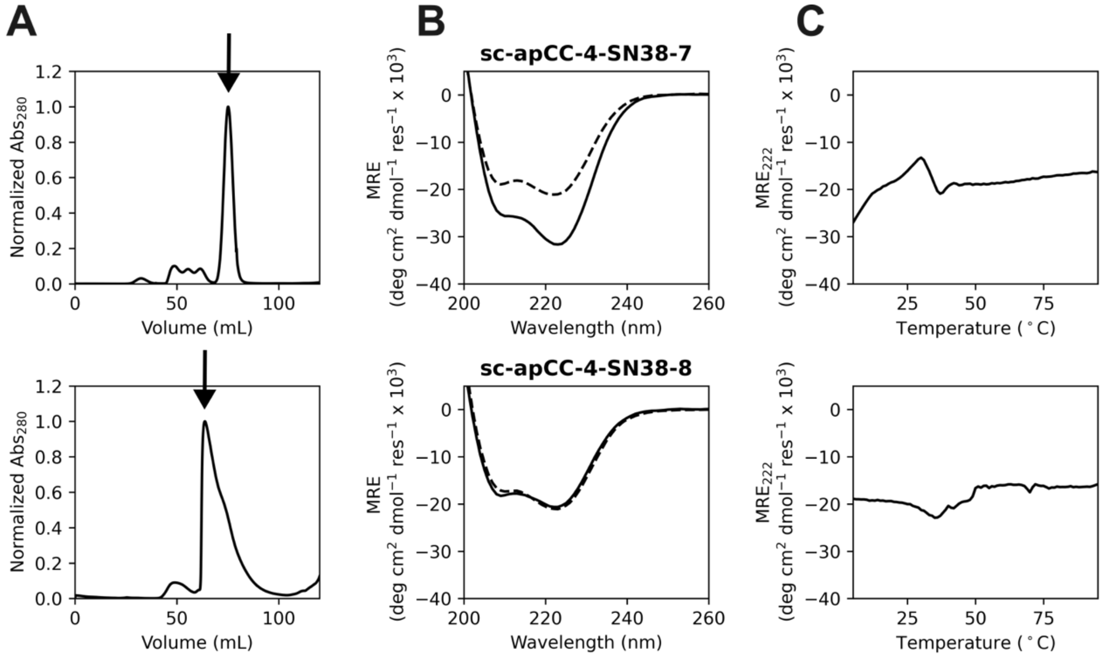


**Supplementary Figure 29. Biophysical characterization of apo sc-apCC-4 variants refined with Boltz2 and LigandMPNN. (A)** Preparative size-exclusion chromatography (SEC) elution profiles. Proteins were purified on either Superdex 75 pg or Superdex 200 pg columns in 50 mM sodium phosphate, pH 7.4, with 150 mM NaCl. The arrow marks the elution fractions selected for further analysis. **(B)** Far-UV circular dichroism (CD) spectral scans collected at 5 °C (solid lines) and after heating to 95 °C followed by cooling to 5 °C (dashed lines). Filled grey regions indicate HT voltage ≥600 V. **Conditions**: 10 µM protein in 50 mM sodium phosphate, 150 mM NaCl, pH 7.4. **(C)** Temperature-dependent CD monitored at 222 nm. Thermal melts were recorded on heating (solid lines) and on cooling (dashed lines) to/from 95 °C. **Conditions**: 10 µM protein in 50 mM sodium phosphate, 150 mM NaCl, pH 7.4.

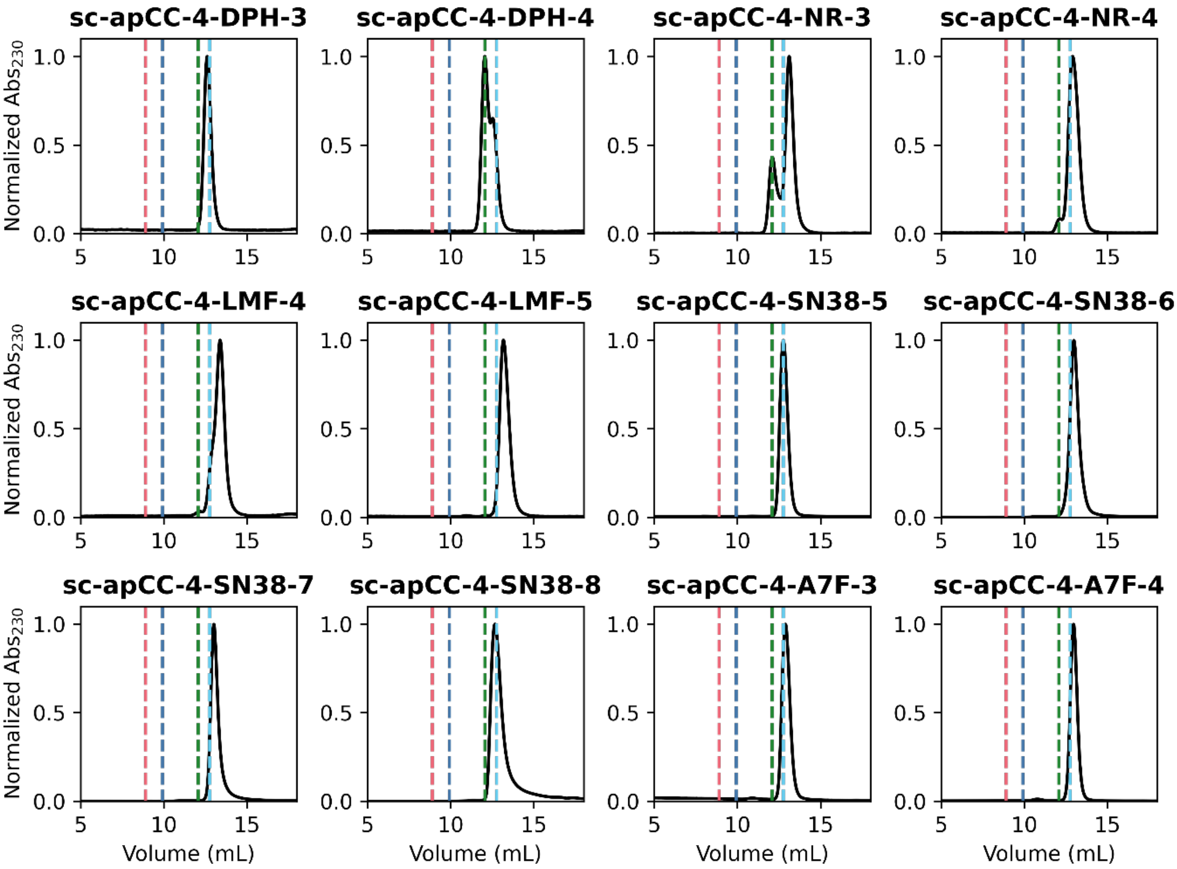


**Supplementary Figure 30. aFPLC traces of 4-helix bundle proteins design with LigandMPNN and Boltz2.** Dashed lines represent elution volumes of calibration standards (Figure S15). **Conditions:** 50 mM sodium phosphate, pH 7.4, 150 mM NaCl. Proteins run on Superdex 75 Increase 10/300 GL at 0.5 – 2 mg/ml.

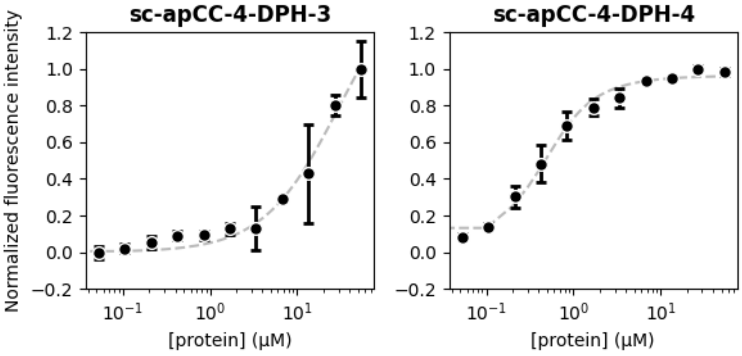


**Supplementary Figure 31. Saturation binding curves for 4-helix bundle DPH binders designed with LigandMPNN and Boltz2.** Fits returned $K_{D(sc\text{-}apCC\text{-}4\text{-}DPH\text{-}3)}$ = 30 ± 6 μM, $K_{D(sc\text{-}apCC4\text{-}DPH\text{-}4)}$ = 0.20 ± 0.03 μM. **Conditions**: 0.5 μM ligand, 0 – 60 μM protein, data are the mean of three independent repeats, error bars represent the standard deviation from the mean. All data collected in PBS, 10% v/v MeCN, pH 7.4.

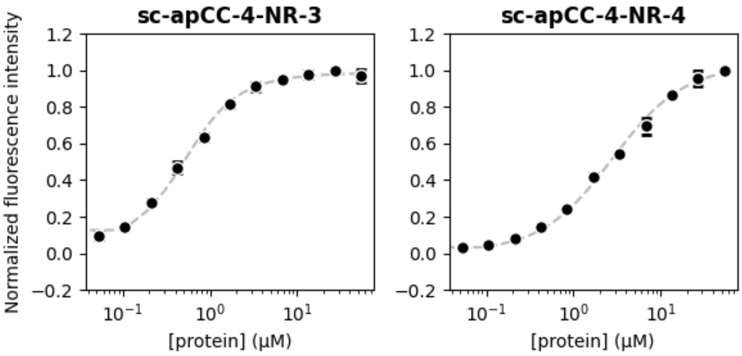


**Supplementary Figure 32. Saturation binding curves for 4-helix bundle NR binders designed with LigandMPNN and Boltz2.** Fits returned $K_{D(\text{sc-apCC-4-NR-3})}$ = 0.24 ± 0.02 μM, $K_{D(\text{sc-apCC4-NR-4})}$ = 2.6 ± 0.2 μM**. Conditions**: 0.5 μM ligand, 0 – 60 μM protein, data are the mean of three independent repeats, error bars represent the standard deviation from the mean. All data collected in PBS, 10% v/v MeCN, pH 7.4.

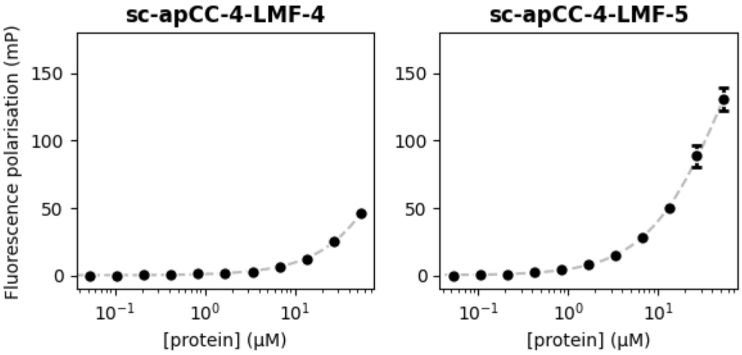


**Supplementary Figure 33. Saturation binding curves for 4-helix bundle LMF binders designed with LigandMPNN and Boltz2.** Fits returned $K_{D(\text{sc-apCC-4-LMF-4})}$ = 350 ± 77 μM, $K_{D(\text{sc-apCC4-LMF-5})}$ = 55 ± 2 μM**. Conditions**: 1 μM ligand, 0 – 60 μM protein, data are the mean of three independent repeats, error bars represent the standard deviation from the mean. All data collected in PBS, 10% v/v DMSO, pH 7.4.

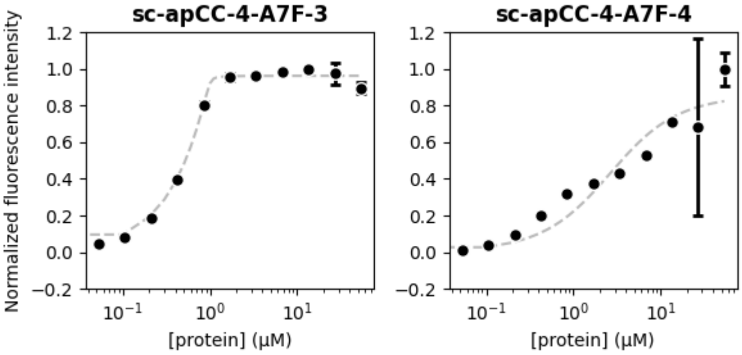

**Supplementary Figure 34. Saturation binding curves for 4-helix bundle arcyriaflavin A binders designed with LigandMPNN and Boltz2.** Fits returned $K_{D(sc\text{-}apCC\text{-}4\text{-}A7F\text{-}3)}$ = 0.002 ± 0.006 μM, $K_{D(sc\text{-}apCC4\text{-}A7F\text{-}4)}$ = 2.1 ± 0.8 μM. **Conditions**: 1 μM ligand, 0 – 60 μM protein, data are the mean of three independent repeats, error bars represent the standard deviation from the mean. All data collected in PBS, 10% v/v MeCN, pH 7.4.

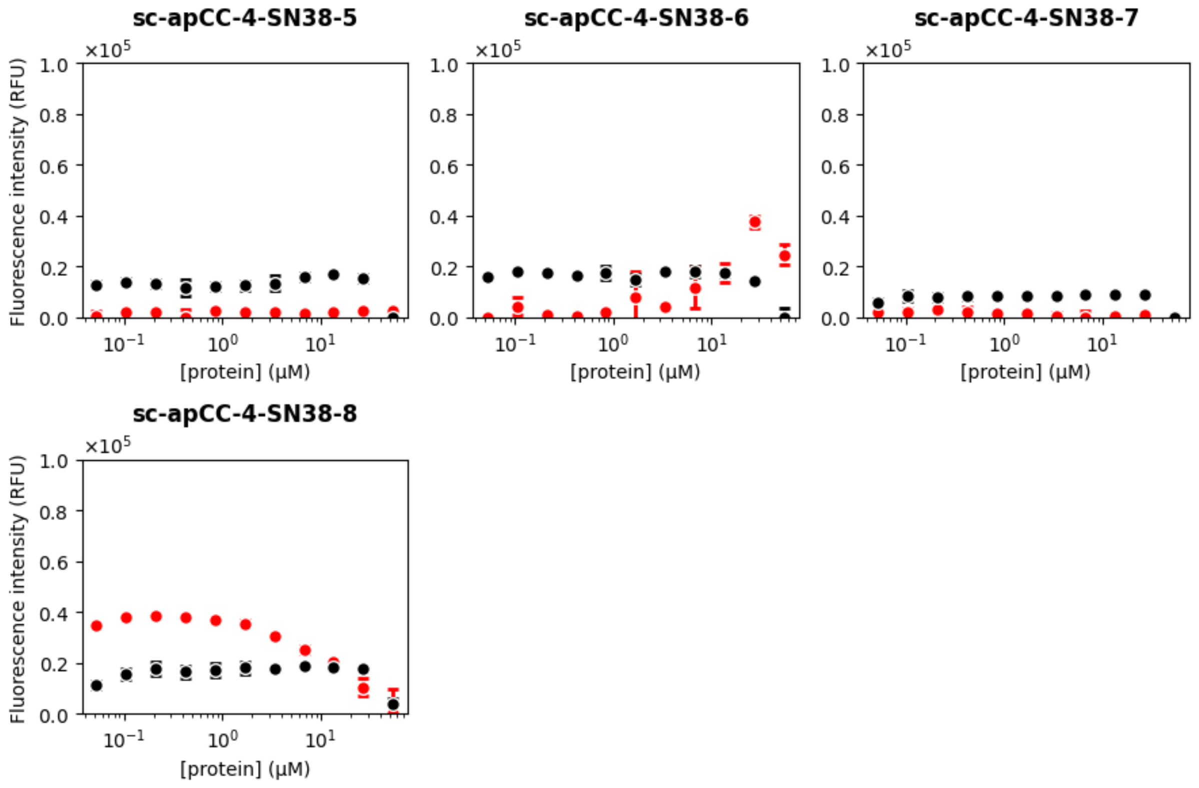


**Supplementary Figure 35. Saturation binding curves for 4-helix bundle SN38 binders designed with LigandMPNN and Boltz2.** No binding can be observed in acidic (red) or neutral (black) pH. **Conditions**: 1 μM ligand, 0 – 60 μM protein, data are the mean of three independent repeats, error bars represent the standard deviation from the mean. All data collected either in ABS, 5% v/v DMSO, pH 4 or PBS, 5% v/v DMSO, pH 7.4.

#### 2.4 Structural characterisation of 4-helix binders

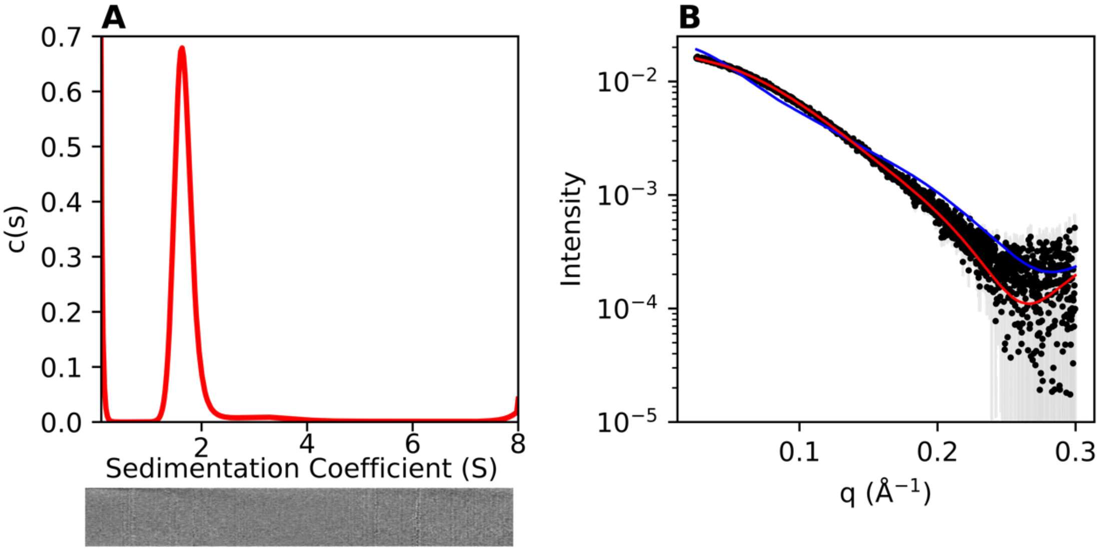


**Supplementary Figure 36. A) Sedimentation velocity and (B) small-angle X-ray scattering analysis of the sc-apCC-4-DPH-2.** Sedimentation velocity fit returned MW = 17.6 kDa (1x monomer), SAXS fits returned **$\chi^2$ = 1.7** for a monomer (red) and **$\chi^2$ = 33.8** for the dimer (blue). **Sedimentation velocity conditions**: 40 000 RPM, 30 µM protein, 50 mM sodium phosphate and 150 mM NaCl (pH 7.4**). SAXS conditions**: 20 mg/ml protein in 20 mM Tris pH 8.0, 50 mM NaCl, 1% v/v glycerol.

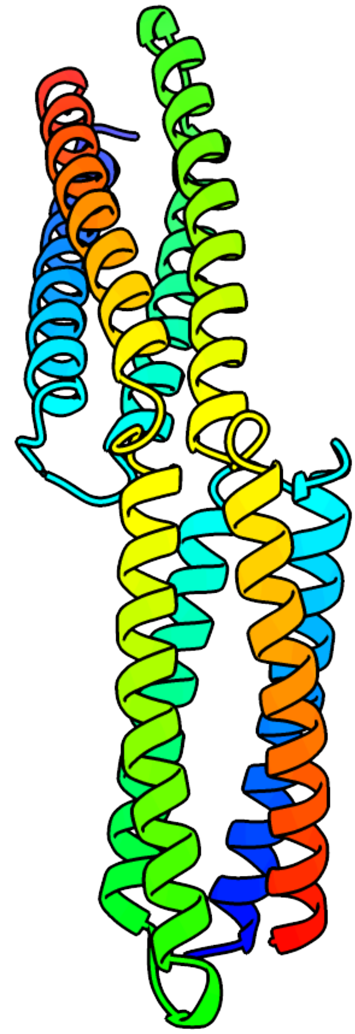

**Supplementary Figure 37. The helix-swapped DPH binder, sc-apCC-4-DPH-2 at 2.4 Å resolution (PDB id 9R1N).** Two chains are coloured as chainbow, transitioning from blue at the N-terminus to red at the C-terminus.

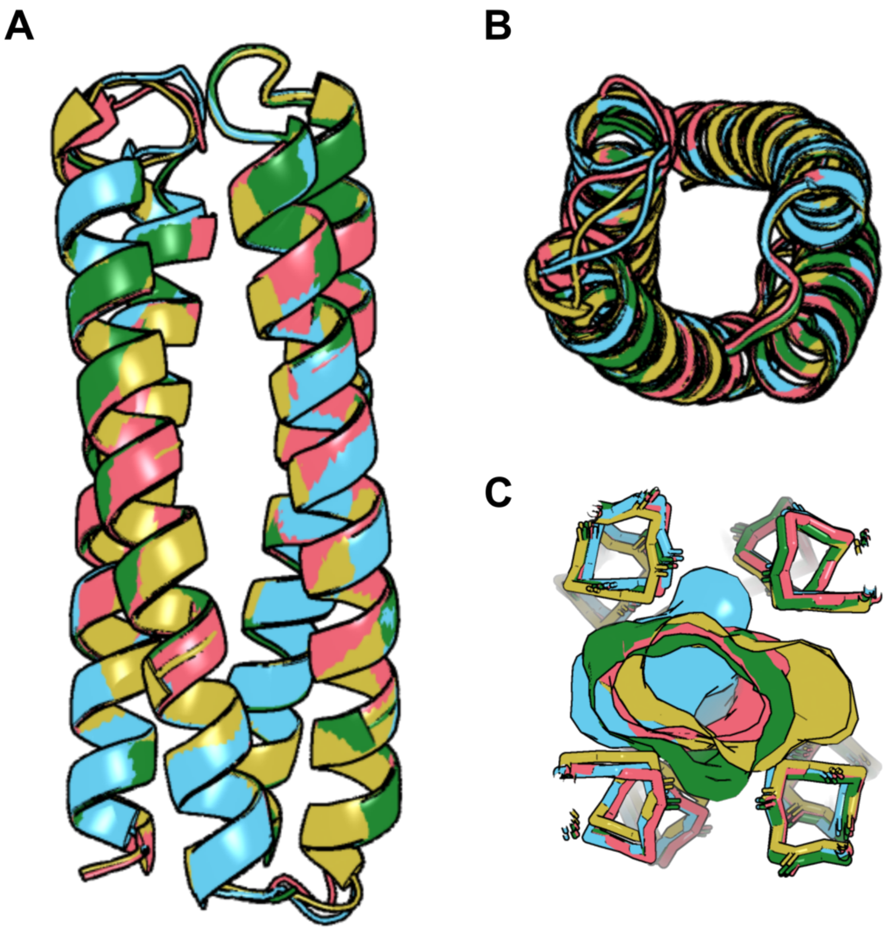


**Supplementary Figure 38. Superimposition of the backbones of the best binders (PDB IDs 9R1J, 9R1M, 9R1K, and 9R1N).** Cartoon representations of the superimposed backbones shown from the side **(A)** and from the top **(B). (C)** Backbone shown as sticks, with a slice through the center of the binding cavity. Structures are coloured as in Figure 2. The DPH binder is omitted due to its domain-swapped architecture.

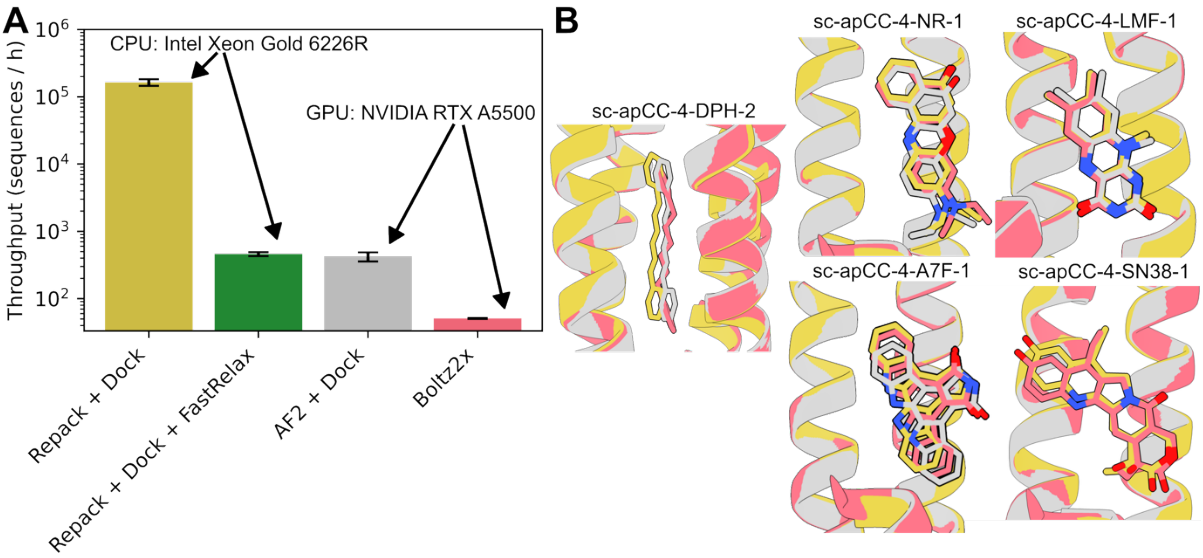


**Supplementary Figure 39. Comparison of protein–ligand modelling**. **(A)** Computational cost comparison. Throughput is reported as the number of sequences modelled per CPU or GPU in one hour. One model was generated per sequence for all methods. AF2 structures were minimised using Amber; Boltz2x predictions used steering potentials. **(B)** Ligand pose comparison between “Repack + Dock” (yellow), “AF2 + Dock” (grey) and Boltz2x (red) predictions. Structures after FastRelax not shown for clarity as the ligand pose did not change during the relaxation. ipTM > 0.9 for all Boltz2x models.

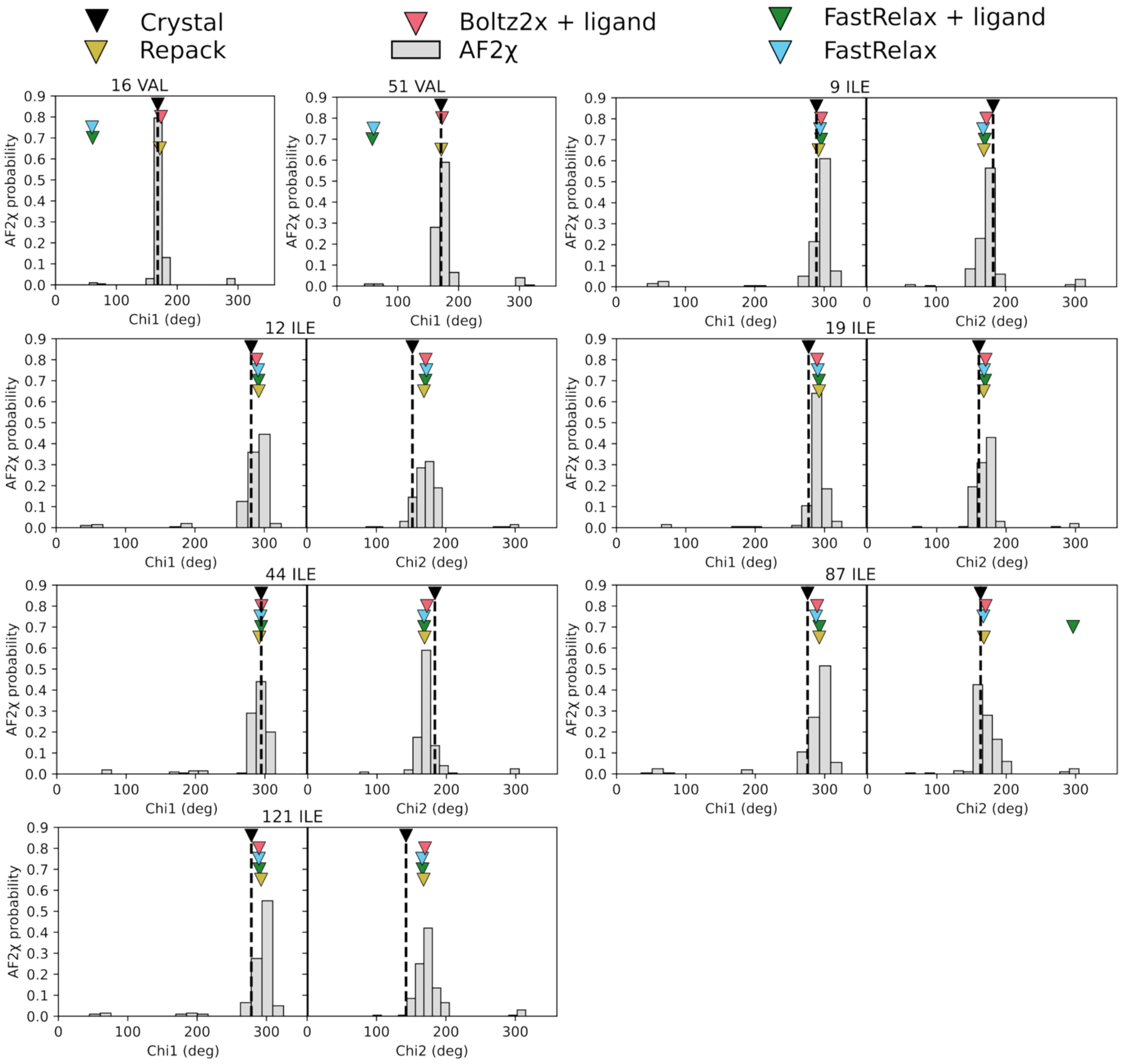


**Supplementary Figure 40. Side-chain rotamer comparison within the binding site across different structural models: crystal, repacked, relaxed, and predicted structures for sc-ap-CC4-DPH-2.** 25 models were generated with Rosetta FastRelax and Boltz2x; only the lowest energy/highest confidence model is shown for clarity as there was no difference between the predicted rotamers. 200 structures were generated with AF2 χ.

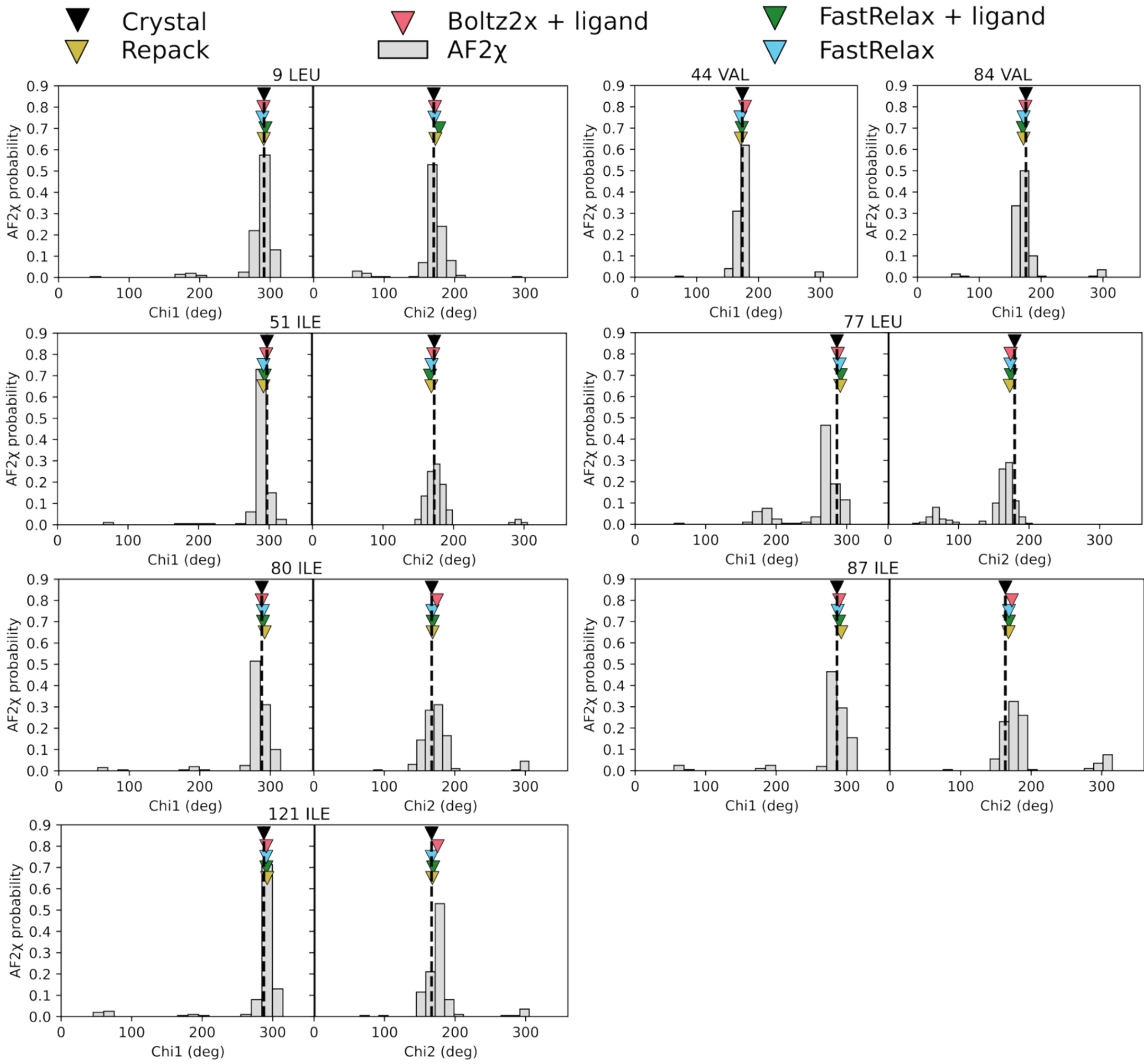


**Supplementary Figure 41. Side-chain rotamer comparison within the binding site across different structural models: crystal, repacked, relaxed, and predicted structures for sc-ap-CC4-NR-1.** 25 models were generated with Rosetta FastRelax and Boltz2x; only the lowest energy/highest confidence model is shown for clarity as there was no difference between the predicted rotamers. 200 structures were generated with AF2 χ.

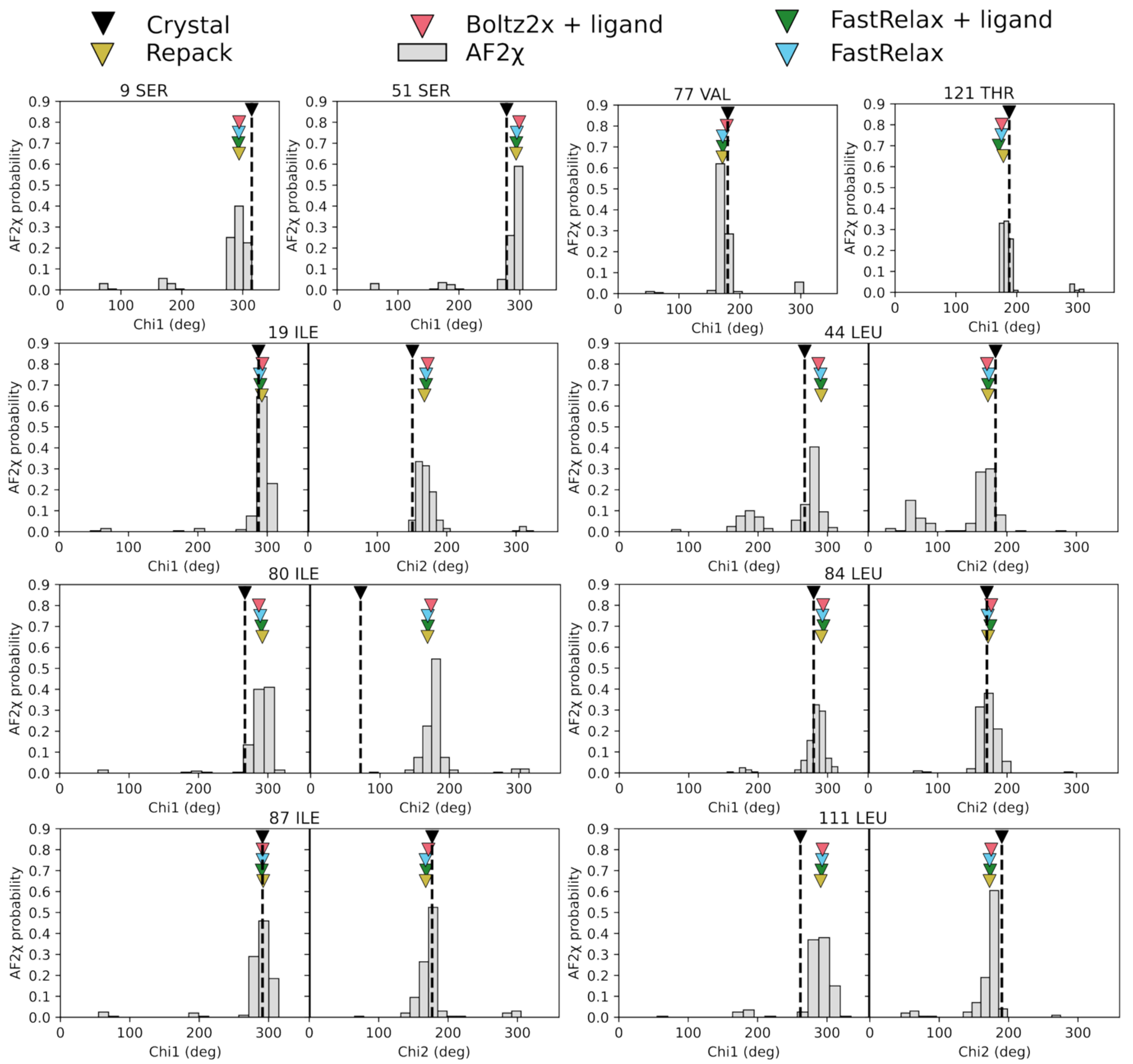


**Supplementary Figure 42. Side-chain rotamer comparison within the binding site across different structural models: crystal, repacked, relaxed, and predicted structures for sc-ap-CC4-LMF-1.** 25 models were generated with Rosetta FastRelax and Boltz2x; only the lowest energy/highest confidence model is shown for clarity as there was no difference between the predicted rotamers. 200 structures were generated with AF2 χ.

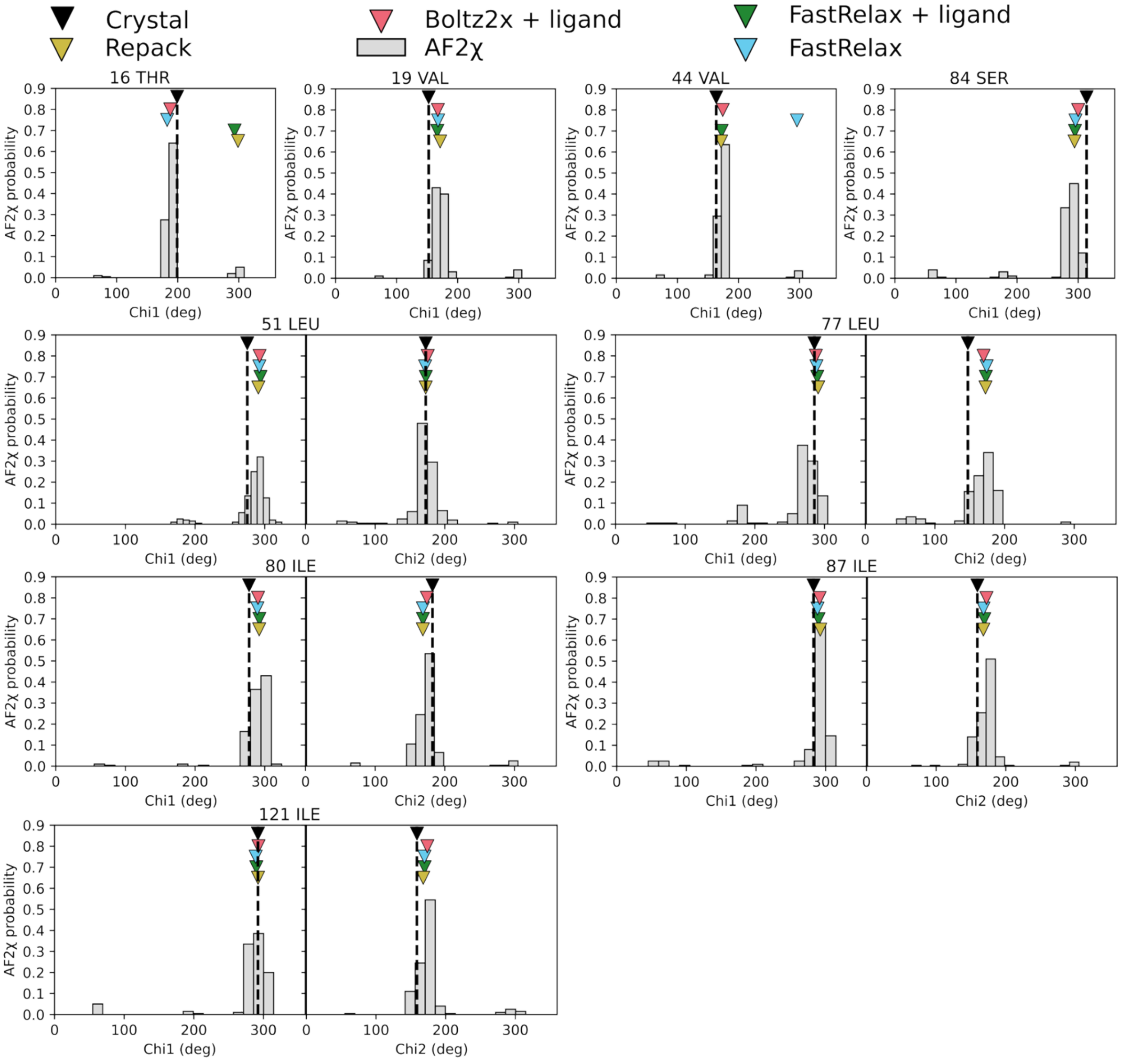


**Supplementary Figure 43. Side-chain rotamer comparison within the binding site across different structural models: crystal, repacked, relaxed, and predicted structures for sc-ap-CC4-A7F-1.** 25 models were generated with Rosetta FastRelax and Boltz2x; only the lowest energy/highest confidence model is shown for clarity as there was no difference between the predicted rotamers. 200 structures were generated with AF2χ.

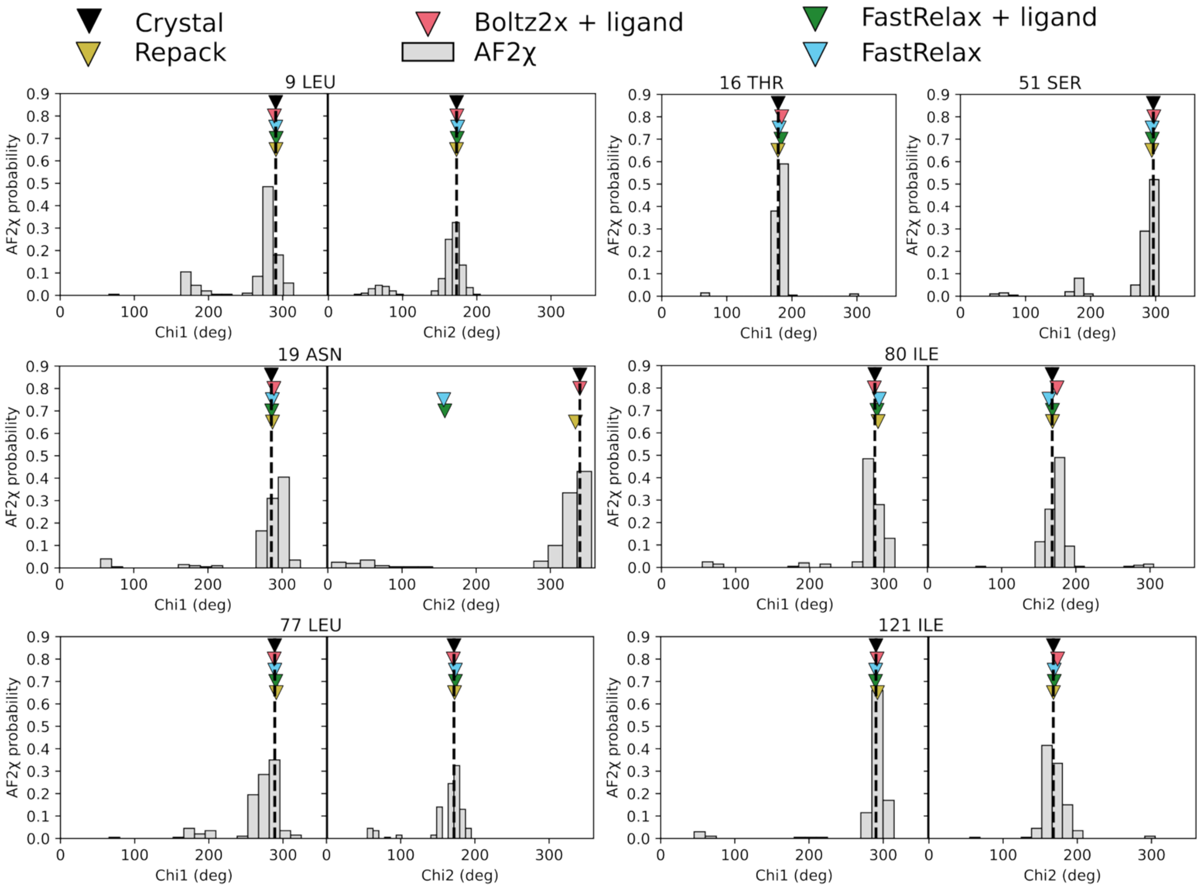


**Supplementary Figure 44. Side-chain rotamer comparison within the binding site across different structural models: crystal, repacked, relaxed, and predicted structures for sc-ap-CC4-SN38-1.** Importantly, Rosetta FastRelax identifies an out-of-distribution rotamer for 19ASN. 25 models were generated with Rosetta FastRelax and Boltz2x; only the lowest energy/highest confidence model is shown for clarity as there was no difference between the predicted rotamers. 200 structures were generated with AF2χ.

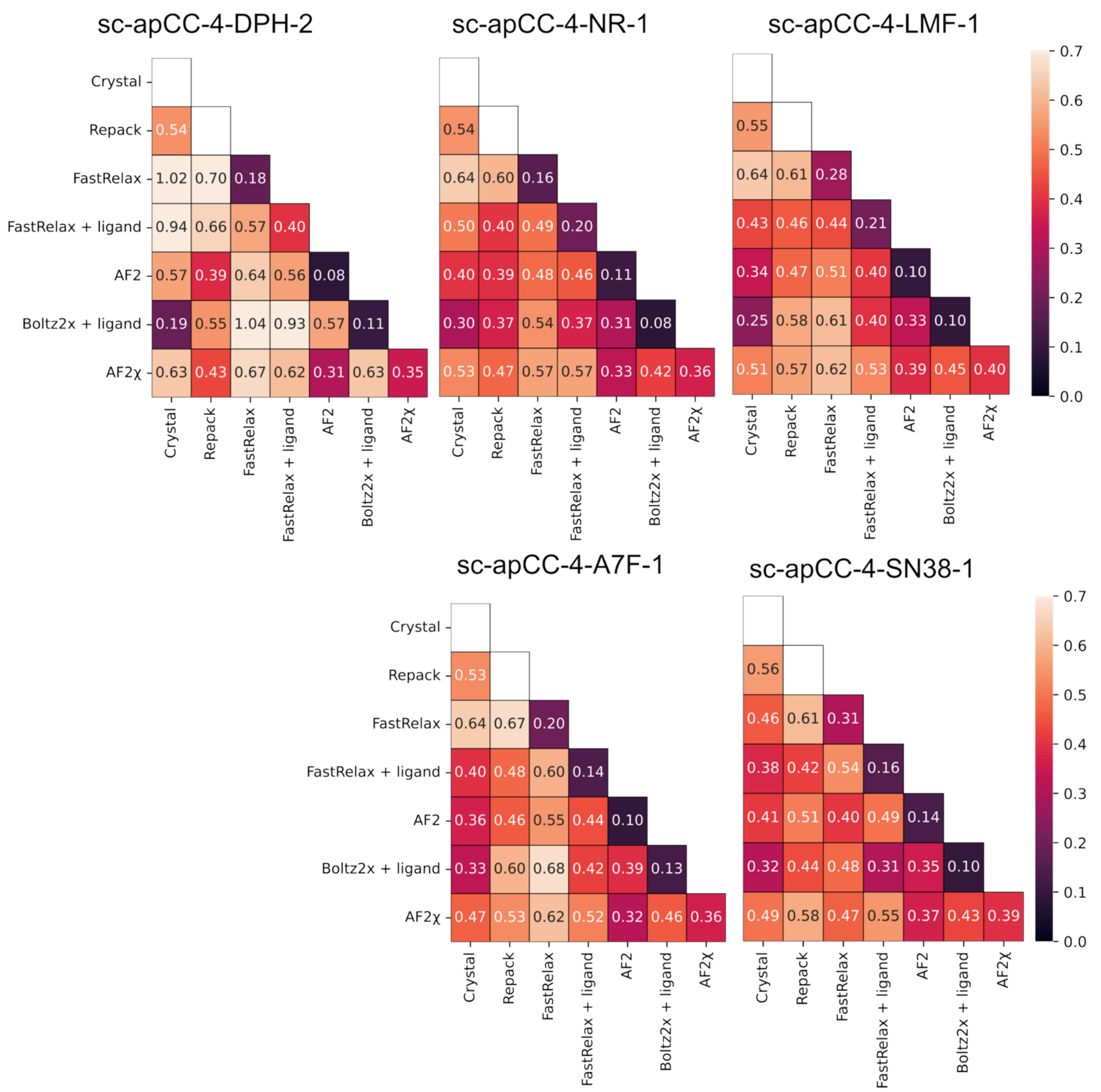


**Supplementary Figure 45. Backbone alignments of the binding site across different structural models: crystal, repacked, relaxed, and predicted structures.** For FastRelax and Boltz2x, 25 structures were generated each. For AF2, 5 models were used to generate 5 structures. For AF2χ, 200 structures were generated. For the DPH design, RMSD calculations account for changes in handedness relative to the parent structure (PDB ID: 8A3K). Overall, Boltz2x with the ligand has the closest binding site geometry to the apo crystal structure, with the repacked and relaxed structures showing similar bbRMSDs.

#### 2.5 Design and characterisation of 6-helix binders using active sampling

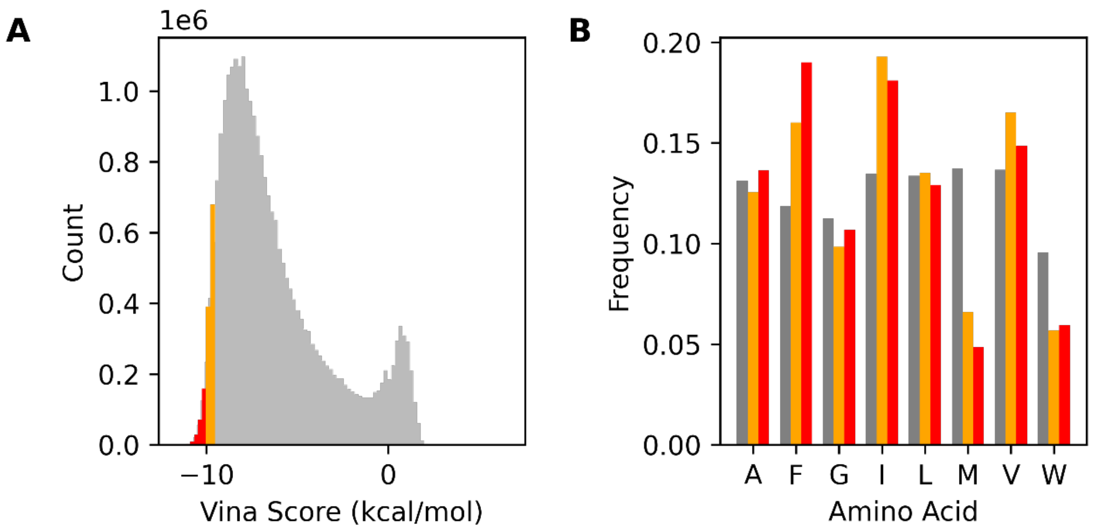


**Supplementary Figure 46. Benchmark dataset for Nile red binding to sc-apCC-6, comprising 26 million sequences.** (A) AutoDock Vina score distribution. (B) Amino acid distribution. Colour scheme: top 1% scores, red; top 5%, orange; all data, grey.

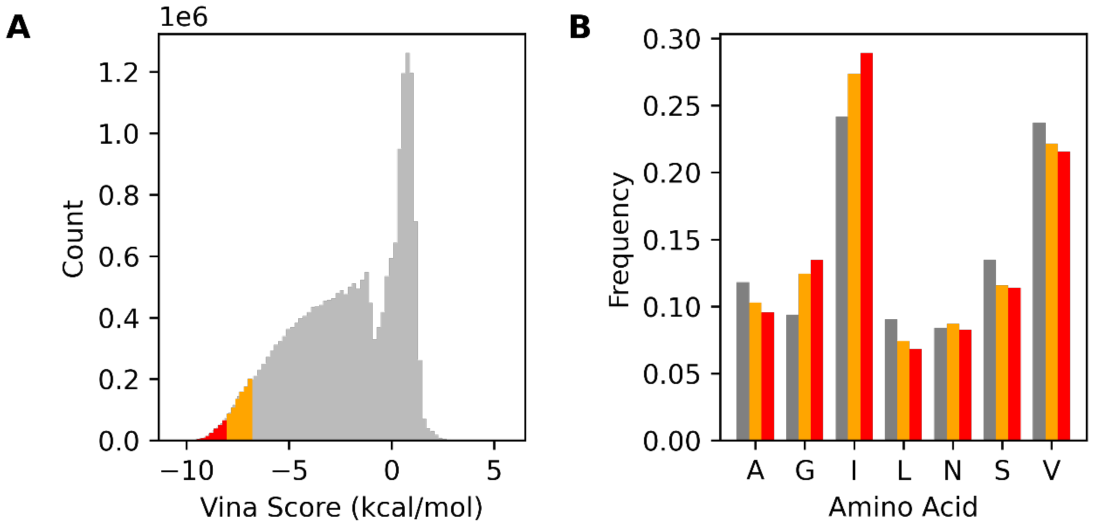


**Supplementary Figure 47. Benchmark dataset for lumiflavin binding to sc-apCC-4, comprising 21 million sequences.** (A) AutoDock Vina score distribution. (B) Amino acid distribution. Colour scheme: top 1% scores, red; top 5%, orange; all data, grey.

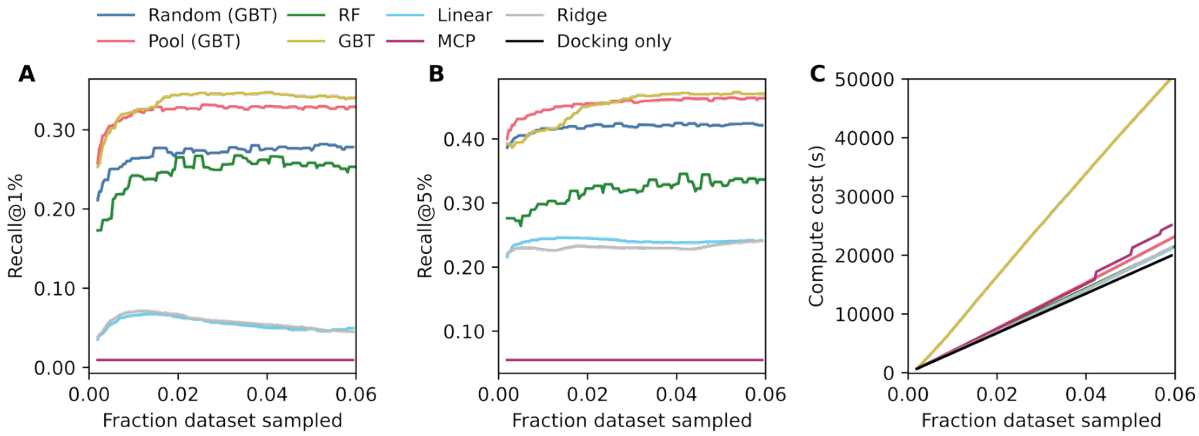


**Supplementary Figure 48. Active sampling benchmark on a sc-apCC-6 set (26 million sequences).** (A) Recall@1% and (B) recall@5% evaluated on a test set of 2.6 million sequences as a function of the fraction of sequences sampled. All models reached convergence after sampling 3% of the dataset. (C) Compute time comparison between brute-force docking (black; 54 cores) and active sampling (54 cores for docking, 4 cores for model training). Conditions: models were initialised with 50,000 randomly sampled sequences, followed by iterative active sampling with a step size of 10,000 sequences using a min–max normalised acquisition function (μ + σ). In pool-based learning, a random subset equal to 100× the step size was sampled at each iteration.

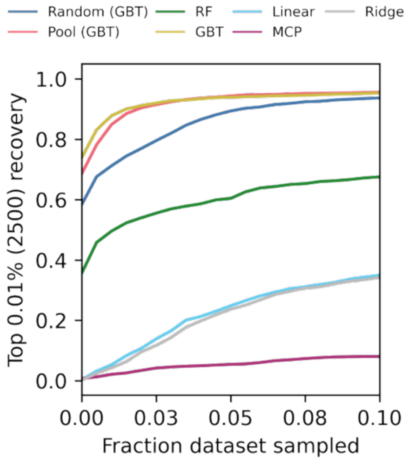


**Supplementary Figure 49.** Number of explicit docking evaluations required to recover the top 0.01% of sequences from the sc-apCC-6 set (26 million sequences). All models were trained on 3% of the dataset.

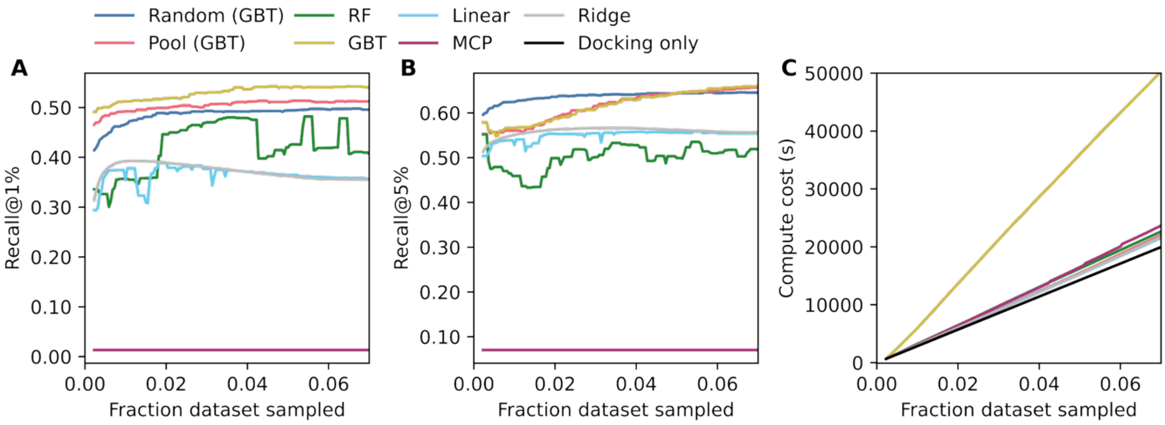


**Supplementary Figure 50. Active sampling benchmark on a sc-apCC-4 set (21 million sequences).** (A) Recall@1% and (B) recall@5% evaluated on a test set of 2.1 million sequences as a function of the fraction of sequences sampled. All models reached convergence after sampling 5% of the dataset. (C) Compute time comparison between brute-force docking (black; 54 cores) and active sampling (54 cores for docking, 4 cores for model training). Conditions: models were initialised with 50,000 randomly sampled sequences, followed by iterative active sampling with a step size of 10,000 sequences using a min–max normalised acquisition function (μ + σ). In pool-based learning, a random subset equal to 100× the step size was sampled at each iteration.

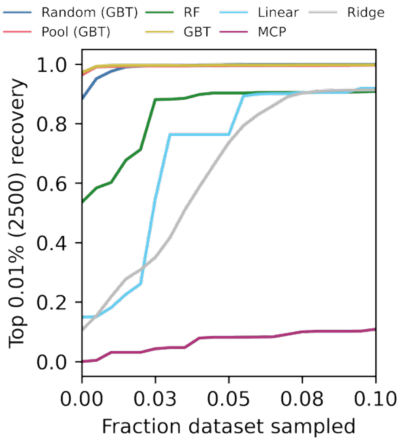


**Supplementary Figure 51.** Number of explicit docking evaluations required to recover the top 0.01% of sequences from the sc-apCC-4 set (21 million sequences). All models were trained on 5% of the dataset.

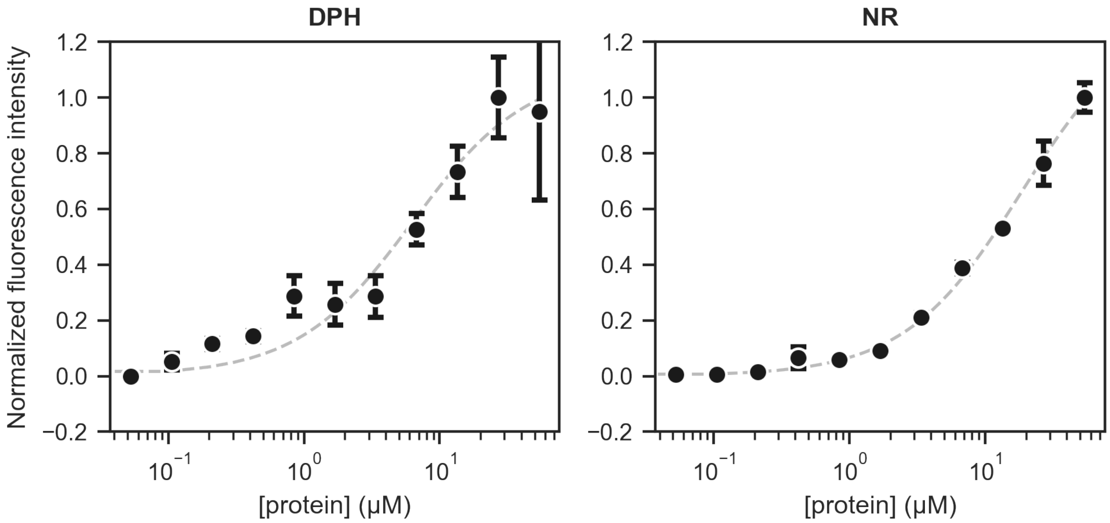


**Supplementary Figure 52. Saturation binding curves for the parent 6-helix bundle (all Leu).** Nile red, orange; DPH, teal. $K_{D(DPH)}$ = 6.1 ± 1.9 μM, $K_{D(NR)}$ > 20 μM. **Conditions**: 0.5 μM, 0 – 60 μM protein, data are the mean of three independent repeats, error bars represent the standard deviation from the mean. All data collected in PBS, 10% v/v MeCN, pH 7.4.

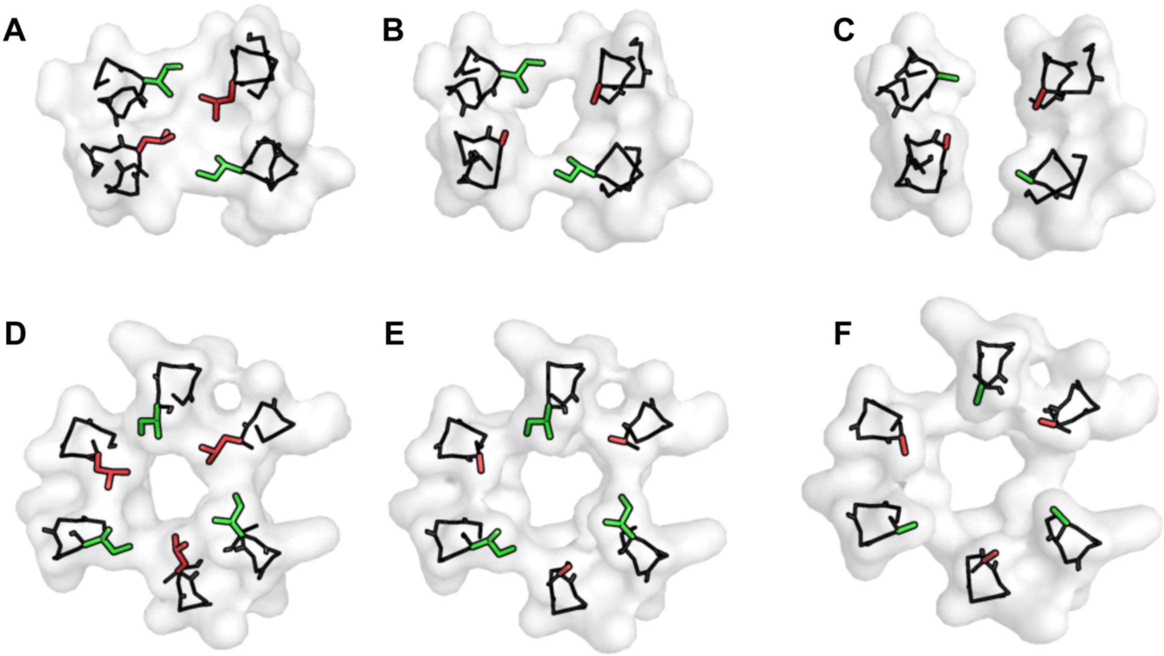


**Supplementary Figure 53. Comparison of the effect of modifying *a* (red) and *d* (green) positions in sc-apCC-4 (ABC; PDB id, 8a3k) and sc-apCC-6 (DEF; PDB id, 8qad). A&D,** original protein; **B&E,** Leu at ***a*** mutated into Ala; **C&F,** Leu at ***a*** and Ile at ***d*** mutated into Ala. In both cases, mutations at **d** create side-channels which are necessary to accommodate the ligands in the smaller sc-apCC-4 but not the sc-apCC-6.

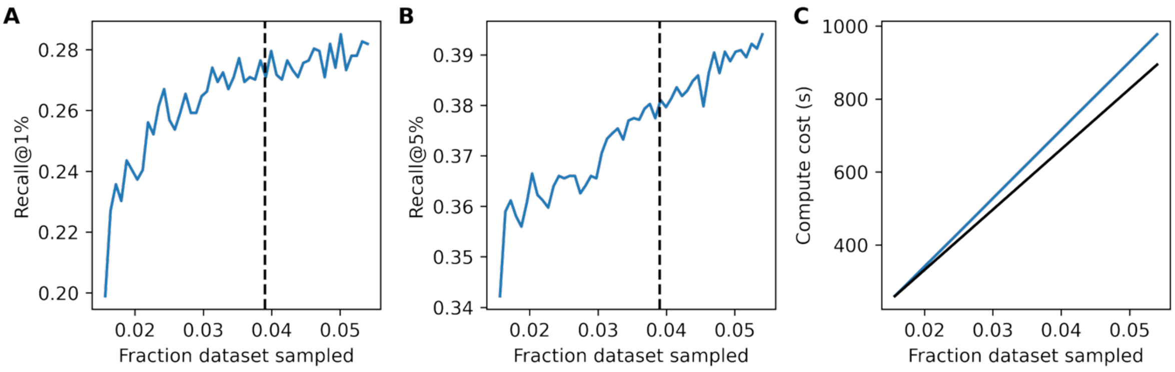


**Supplementary Figure 54. Active sampling with GBT on a small sc-apCC-6 set (1 277 192 sequences).** (A) Recall@1% and (B) recall@5% evaluated on a test set of 20,000 sequences. For experimental validation, active sampling was terminated after ~50,000 sequences (dashed black line), as recall@1% showed no substantial improvement. For benchmarking, sampling was continued beyond this point: although ranking of top sequences plateaued, overall model performance, as measured by recall@5%, continued to improve. (C) Compute time comparison between brute-force docking (black; 54 cores) and active sampling (54 cores for docking, 4 cores for model training).

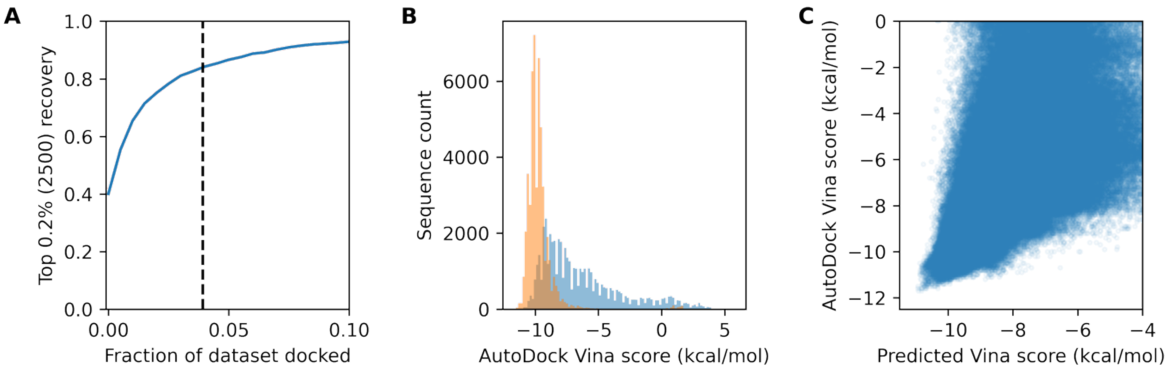


**Supplementary Figure 55. Analysis of the GBT model after active sampling on a small sc-apCC-6 set.** (A) Number of explicit docking evaluations required to recover the top 0.02% of sequences. The dashed line indicates the 50,000 sequences selected for explicit docking and experimental validation. (B) Comparison of AutoDock Vina scores for the top 50,000 sequences predicted by the GBT model versus a randomly sampled subset from the full dataset, showing strong enrichment of low-energy sequences. (C) Correlation between predicted and true AutoDock Vina scores across the full dataset after active sampling 50,000 sequences (Figure S52), with a Spearman correlation of 0.8.

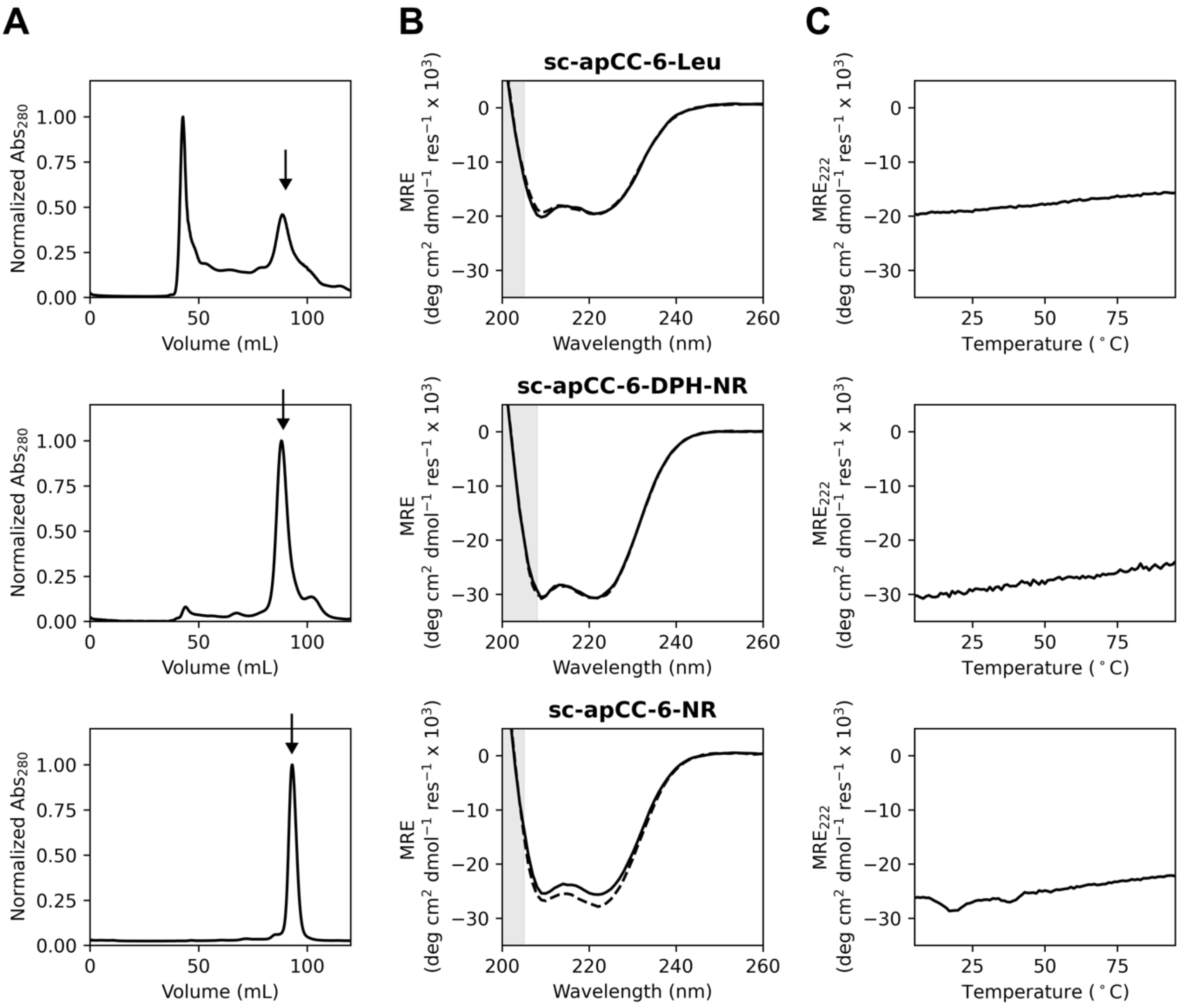


**Supplementary Figure 56. Biophysical characterization of apo sc-apCC-6 variants. (A)** Preparative size-exclusion chromatography (SEC) elution profiles. Proteins were purified on either Superdex 75 pg or Superdex 200 pg columns in 50 mM sodium phosphate, pH 7.4, with 150 mM NaCl. The arrow marks the elution fractions selected for further analysis. **(B)** Far-UV circular dichroism (CD) spectral scans collected at 5 °C (solid lines) and after heating to 95 °C followed by cooling to 5 °C (dashed lines). Filled grey regions indicate HT voltage ≥600 V. **Conditions**: 10 µM protein in 50 mM sodium phosphate, 150 mM NaCl, pH 7.4. **(C)** Temperature-dependent CD monitored at 222 nm. Thermal melts were recorded on heating (solid lines) and on cooling (dashed lines) to/from 95 °C. **Conditions**: 10 µM protein in 50 mM sodium phosphate, 150 mM NaCl, pH 7.4.

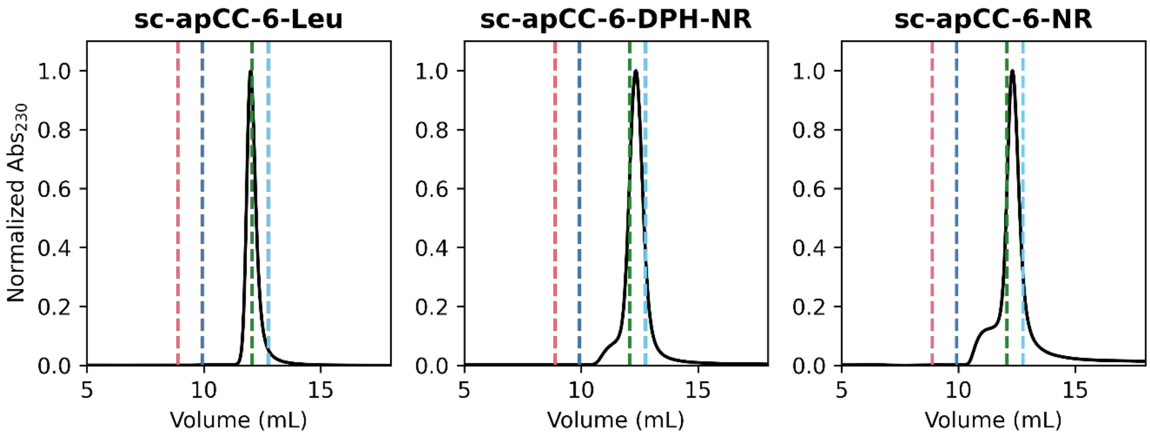


**Supplementary Figure 57. aFPLC traces of all 6-helix bundle proteins in this study.** Dashed lines represent elution volumes of calibration standards (Figure S13). **Conditions:** 50 mM sodium phosphate, pH 7.4, 150 mM NaCl. Proteins run on Superdex 75 Increase 10/300 GL at 0.5 – 2 mg/ml.

#### 2.6 Spectroscopic characterisation and simulations of the double-binder

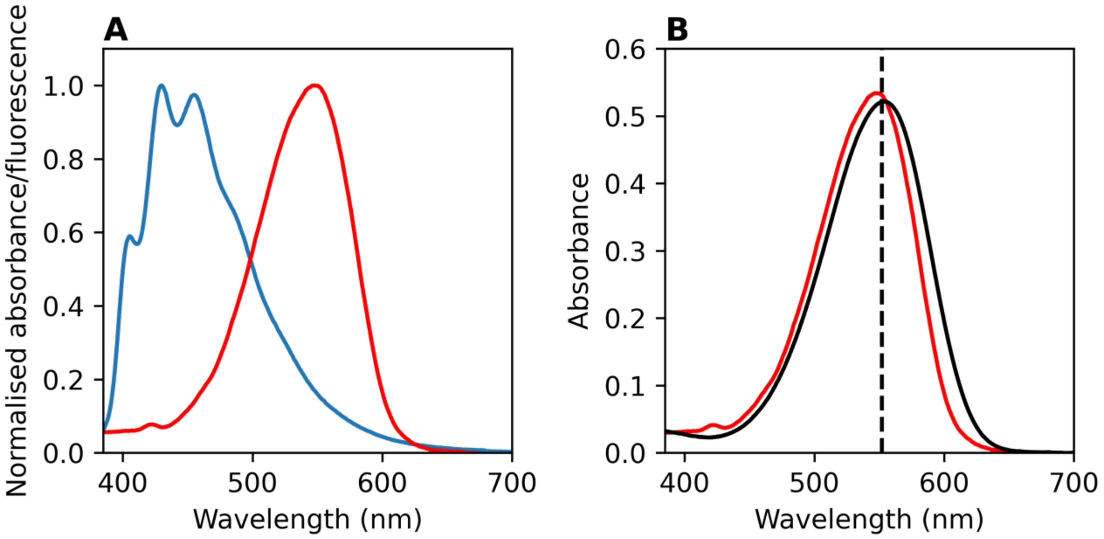


**Supplementary Figure 58. Spectral overlap integral (7.79 x $10^{14}$ nm$^4$ M$^{-1}$ cm$^{-1}$) calculation between DPH and Nile red in the double binder.** **(A)** Normalised DPH emission (blue) and Nile red absorption (red). **(B)** Absorption spectrum for Nile red in the double binder (red) and methanol (black). Molar extinction coefficient for Nile red in methanol (45 000 M$^{-1}$cm$^{-1}$ at 552 nm; dashed black) was used for the spectral overlap calculation. **Conditions:** 10 µM of either DPH or Nile red, 30 µM protein, PBS, 10% v/v MeCN, pH 7.4 or 10 µM Nile red in methanol.

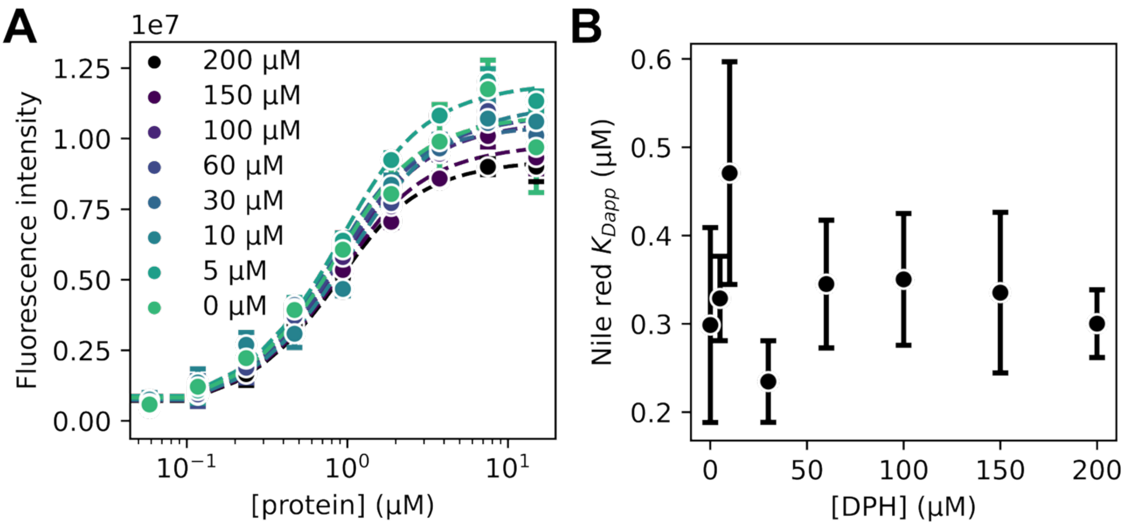


**Supplementary Figure 59. Apparent binding affinity of Nile red in the presence of varying DPH concentrations.** **(A)** Titration curves showing Nile red fluorescence intensity. **(B)** Fitted apparent $K_D$ values for Nile red in presence of DPH. **Conditions**: 1 µM Nile red, 0 – 15 µM protein, 0 – 200 µM DPH, data are the mean of two independent repeats, error bars represent the standard deviation from the mean. All data collected in PBS, 10% v/v MeCN, pH 7.4.

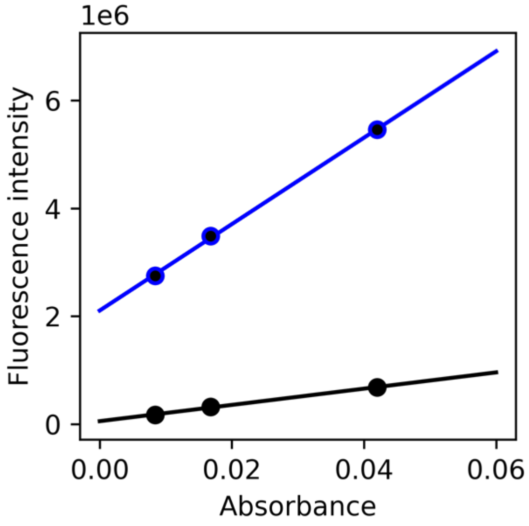


**Supplementary Figure 60. Fluorescence quantum yield estimation for DPH in the double binder (blue, Φ = 85%) by comparison to a standard – DPH in MeCN (black, Φ = 17%). Conditions**: 30

μM protein, data are the mean of three independent repeats, error bars (standard deviation from the mean) fall within the size of the markers.

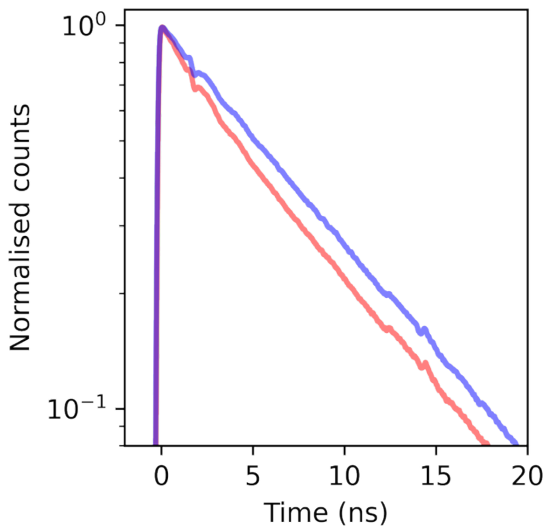


**Supplementary Figure 61. Normalised TCSPC data for DPH with (red) and without (blue) Nile red after excitation at 352 nm.** For the binder containing only DPH, a mono-exponential fit returned τ(DPH) = 7.6 ± 0.2 ns. **Conditions:** 20 μM of each ligand, 30 μM protein, PBS, 10% v/v MeCN, pH 7.4.

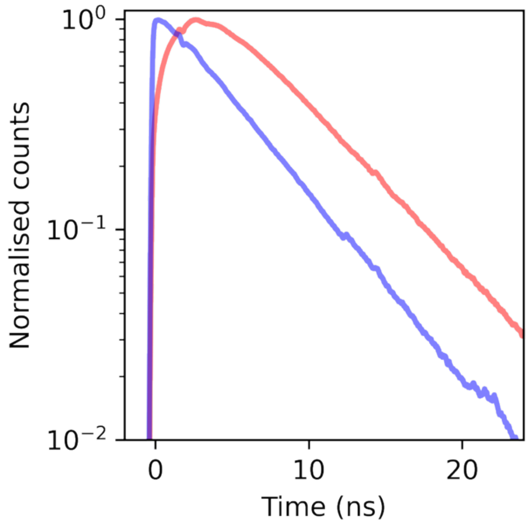


**Supplementary Figure 62. Normalised TCSPC data for Nile red with (red) and without (blue) DPH after excitation at 352 nm.** Emission from Nile red after direct excitation without DPH was accumulated for 20-times longer to account for the low Nile red absorption at this excitation wavelength. Data fit to a mono-exponential decay returned $\tau$(Nile red) = 5.2 ± 0.2 ns. **Conditions:** 20 μM of each ligand, 30 μM protein, PBS, 10% v/v MeCN, pH 7.4.

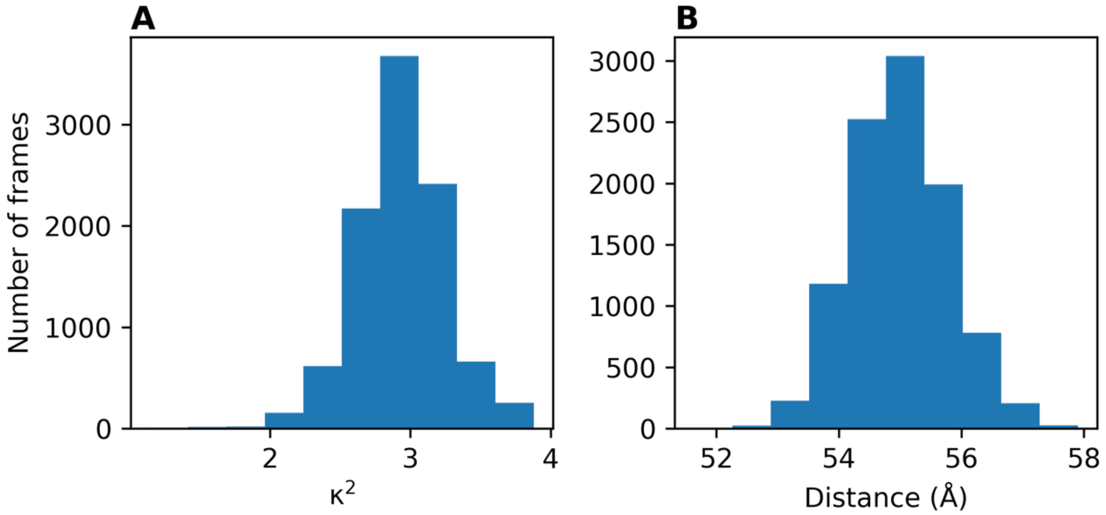


**Supplementary Figure 63. (A) Dipole orientation factor ($\kappa^2$) and (B) centre of mass DPH – Nile red distances derived from MD simulations.** The starting model was generated with AlphaFold2, ligands were docked with AutoDock Vina. 5 independent 100 ns simulations were performed. For both ligands, transition dipoles were taken as the long axes of the molecule. $\kappa^2$ = 2.9 ± 0.3 was used for ligand distance calculation from the experimental data using the Förster equation.

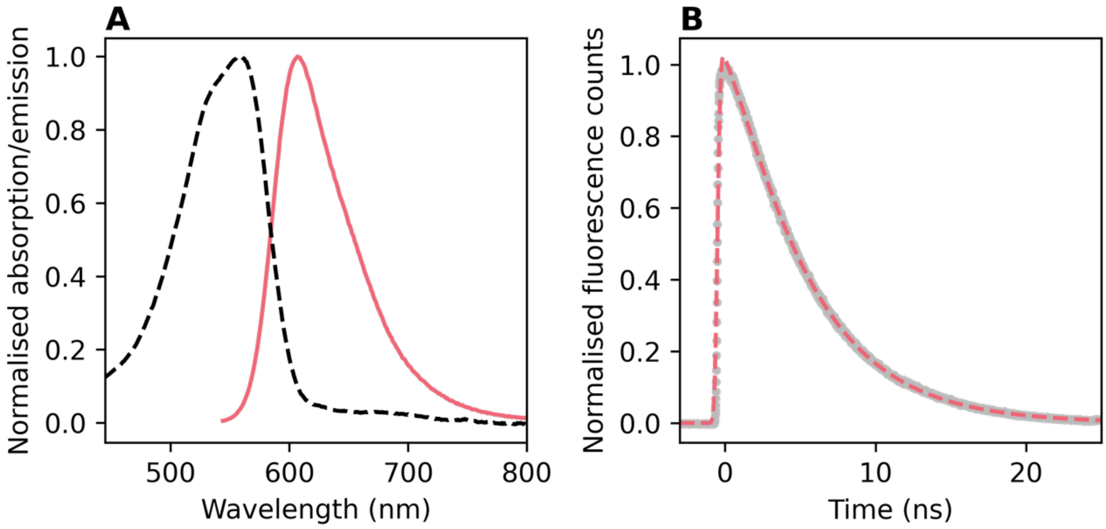


**Supplementary Figure 64. Spectral properties of Nile red bound to sc-apCC-4-SN38-1.** (A) Absorption (dashed) and emission (solid, $\lambda_{ex}$ = 510 nm) spectra. (B) Time-correlated single photon counting data showing fluorescence decay after excitation at 560 nm. The data was fitted to a mono-exponential decay with a rise component[2]. $\tau_{NR}$ = 4.8 ± 0.2 ns. **Conditions:** 5 μM Nile red, 50 μM protein, PBS, 10% v/v MeCN, pH 7.4.

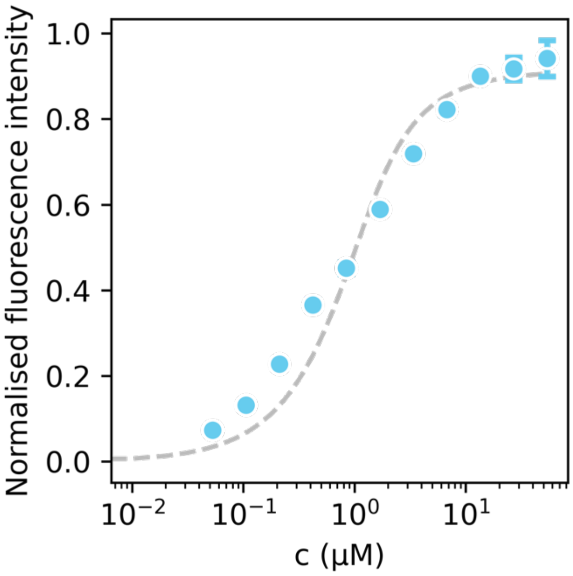


**Supplementary Figure 65. Saturation binding data and fits for Nile blue binding to sc-apCC-4-SN38-1.** Kd = 0.40 ± 0.13 μM. **Conditions**: 1 μM Nile blue, 0 – 60 μM protein, data are the mean of three independent repeats, error bars represent the standard deviation from the mean.

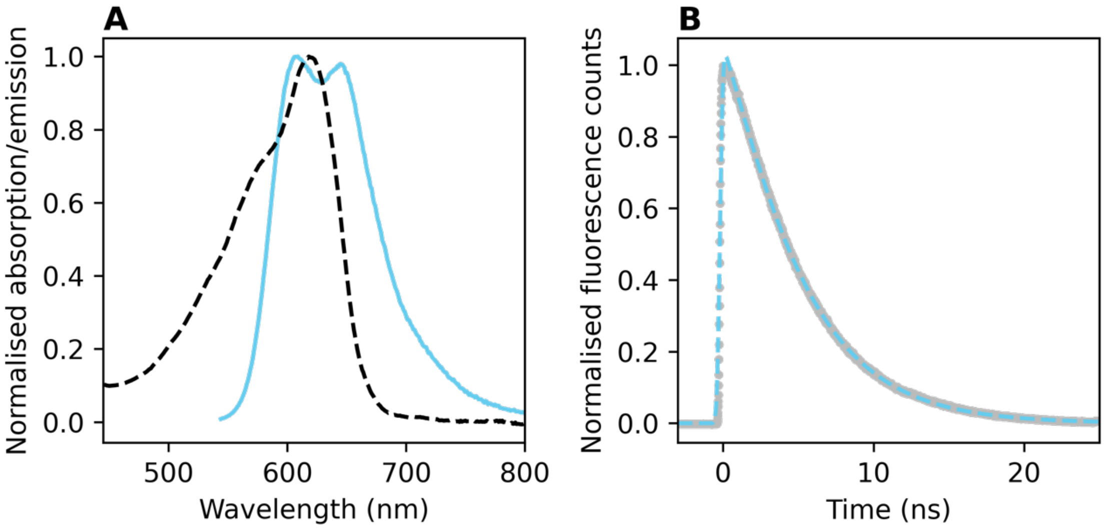


**Supplementary Figure 66. Spectral properties of Nile blue bound to sc-apCC-4-SN38-1.** (A) Absorption (dashed) and emission (solid, $\lambda_{ex}$ = 530 nm) spectra. (B) Time-correlated single photon counting data showing fluorescence decay after excitation at 560 nm. The data was fitted to a mono-exponential decay with a rise component[2]. $\tau_{NR}$ = 4.2 ± 0.2 ns. **Conditions:** 5 μM Nile blue, 50 μM protein, PBS, 10% v/v MeCN, pH 7.4.

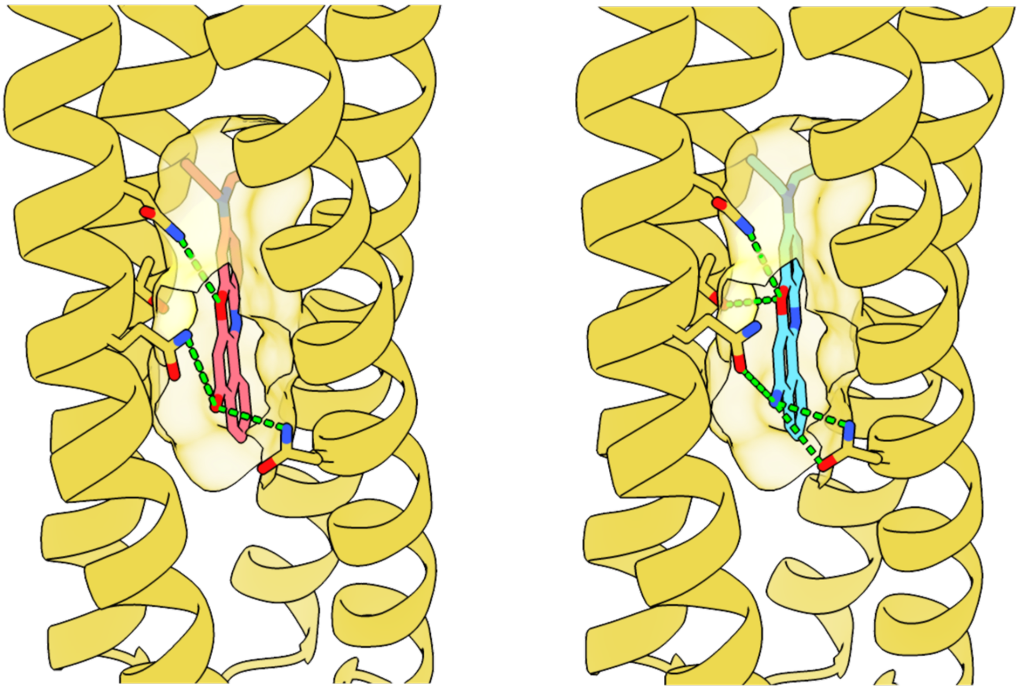

**Supplementary Figure 67. Nile red (red) and Nile blue (cyan) docked into the crystal structure of sc-apCC-4-SN38-1 (PDB id 9R1L).** Nile red Vina score: -10.4 kcal/mol, Nile blue Vina score: -10.6 kcal/mol. Sc-apCC-4-SN38-1 structure was energy-minimised using Rosetta prior to docking.

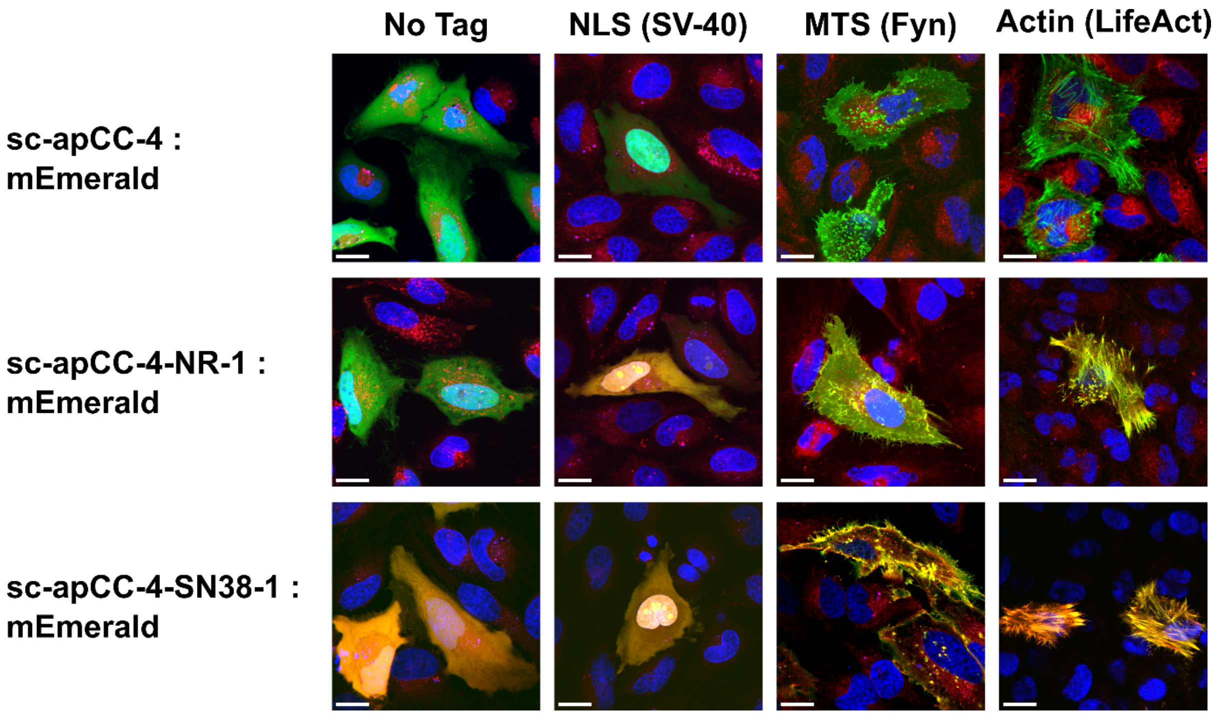


**Supplementary Figure 68.** Fluorescence confocal microphotographs of live HeLa cells expressing the non-binding sc-apCC-4 scaffold (top row) or the designer binders sc-apCC-4-NR-1 (middle row), sc-apCC-4-SN38-1 (bottom row) as fusions to mEmerald and stained with Nile red and Hoechst 33342. Cytosolic constructs as well as fusions to short sequences targeting the nucleus, plasma membrane, and actin are presented in the different columns. The composite images show mEmerald (green: ex/em 488/500-550 nm), NR (red: ex/em 561/580-654 nm), and Hoechst 33342 (blue: ex/em 405/417-477 nm) channels. Scalebar = 20 µm.

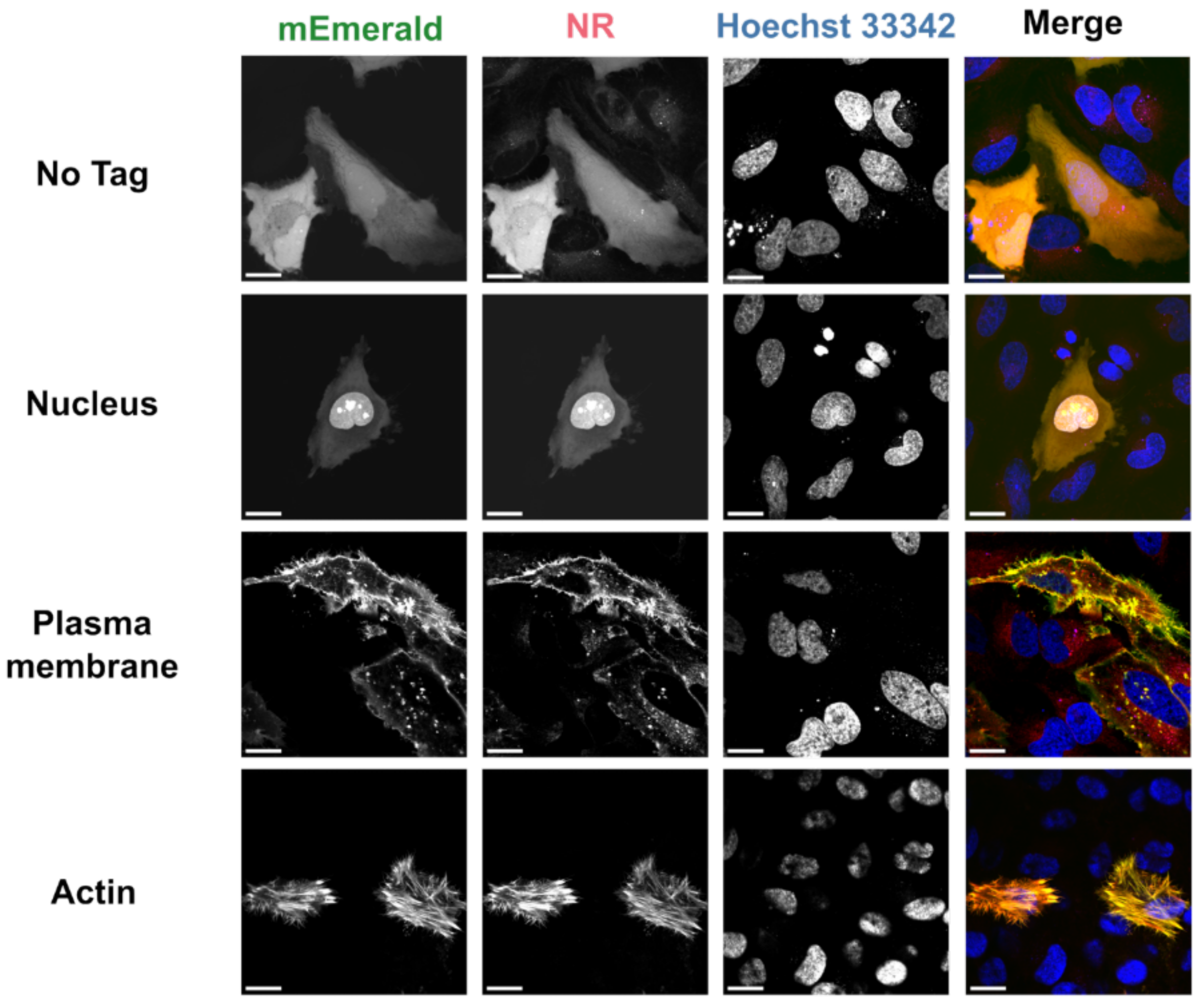


**Supplementary Figure 69.** Expanded version of Figure 68 focussing on the fusions with sc-apCC-4-SN38-1 in more detail. **(Top row**) Fluorescence confocal microphotographs of live HeLa cells transiently expressing a fusion of sc-apCC-4-SN38-1 and mEmerald, and stained by adding 0.1 μM Nile red and Hoechst 33342 to the cell media 30 minutes before imaging. (**Rows 2 – 4**) Similar images for additional fusions with *N*-terminal tags for locating the proteins to the nucleus, plasma membrane, and actin filaments. Merge (right-hand column): mEmerald (green), NR (red), and Hoechst 33342 (blue). Maximum z-projections are shown for constructs with cytosolic and nuclear localization, single-z slices are shown for actin- and membrane-targeting fusions; scalebars = 20 μm.

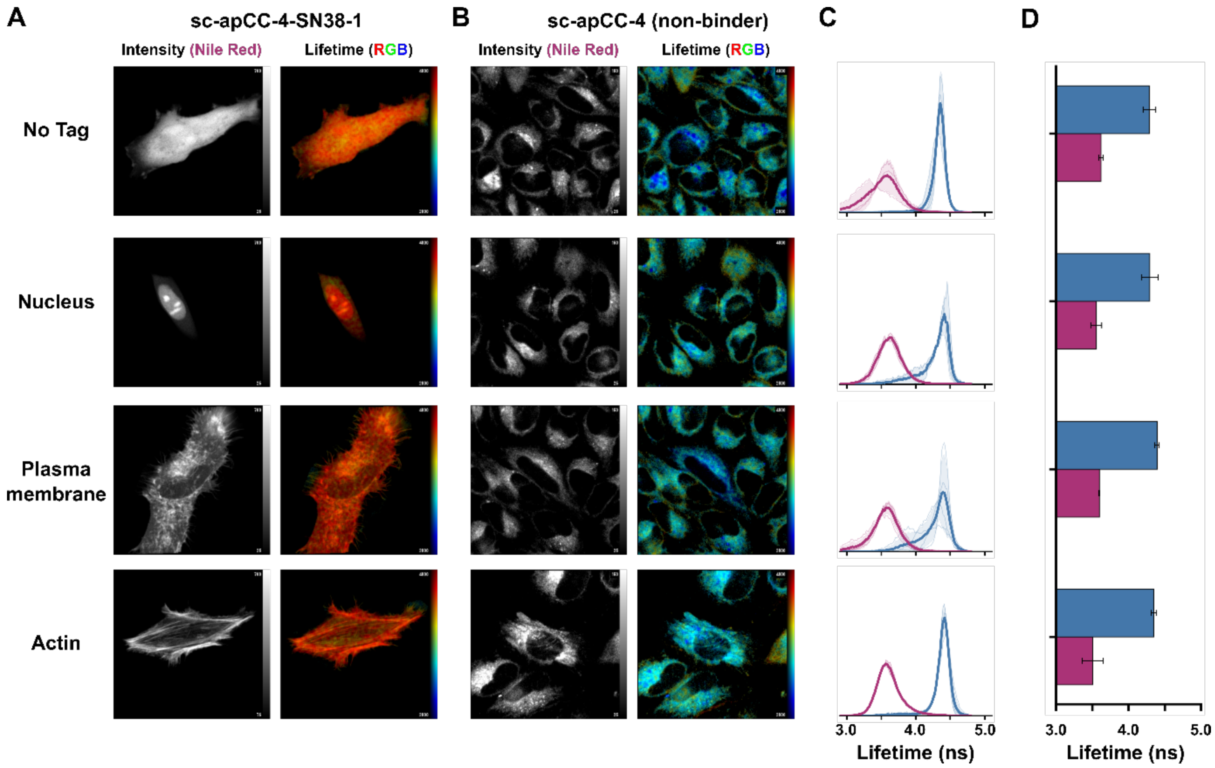


**Supplementary Figure 70. FLIM for Nile red in live HeLa cells expressing either (A) sc-apCC4-SN38-1 or (B) the non-binding parent sc-apCC4 fused to mEmerald.** (A) Fluorescence intensity (1$^{st}$ column) scaled up to 700 units and intensity weighted mean lifetime (2$^{nd}$ column) scaled from 2.8 to 4.8 ns. (B) Fluorescence intensity (1$^{st}$ column) scaled up to 160 units and intensity weighted mean lifetime (2$^{nd}$ column) scaled from 2.8 to 4.8 ns. (C) The distribution of Nile red fluorescence lifetimes across pixels above threshold for sc-apCC4-SN38-1 (blue line) and sc-apCC-4 (deep pink line). (D) Per-image weighted mean lifetime for sc-apCC4-SN38-1 (blue bars) and sc-apCC-4 (deep pink bars).

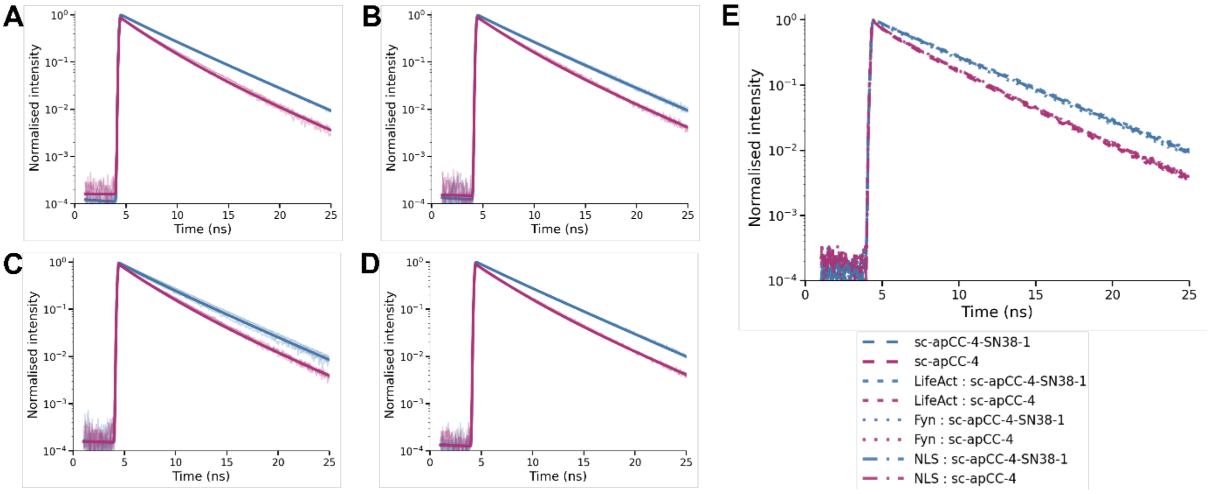


**Supplementary Figure 71. Fluorescence decays** measured for Nile red in HeLa cells expressing (A) sc-apCC-4-SN38-1:mEmerald and sc-apCC-4:mEmerald, (B) Fyn:sc-apCC-4-SN38-1:mEmerald and Fyn:sc-apCC-4:mEmerald; (C) NLS:sc-apCC-4-SN38-1:mEmerald and NLS:sc-apCC-4:mEmerald; (D) LifeAct:sc-apCC-4-SN38-1:mEmerald and LifeAct:sc-apCC-4:mEmerald; (E) (E) all curves for the c-apCC-4-SN38-1:mEmerald (blue) and sc-apCC-4:mEmerald (pink) fusions from panels (A-D) overlaid.

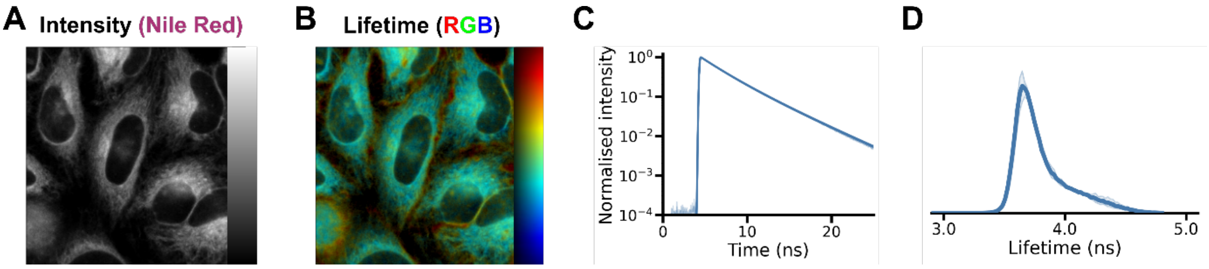


**Supplementary Figure 72. FLIM for Nile red in live non-transfected HeLa cells**. (A) Fluorescence intensity (1st column) scaled up to 160 units. (B) Intensity weighted mean lifetime (2nd column) scaled from 2.8 to 4.8 ns. (C) Fluorescence decays measured for Nile red. (D) The distribution of Nile red fluorescence lifetimes across pixels above threshold.

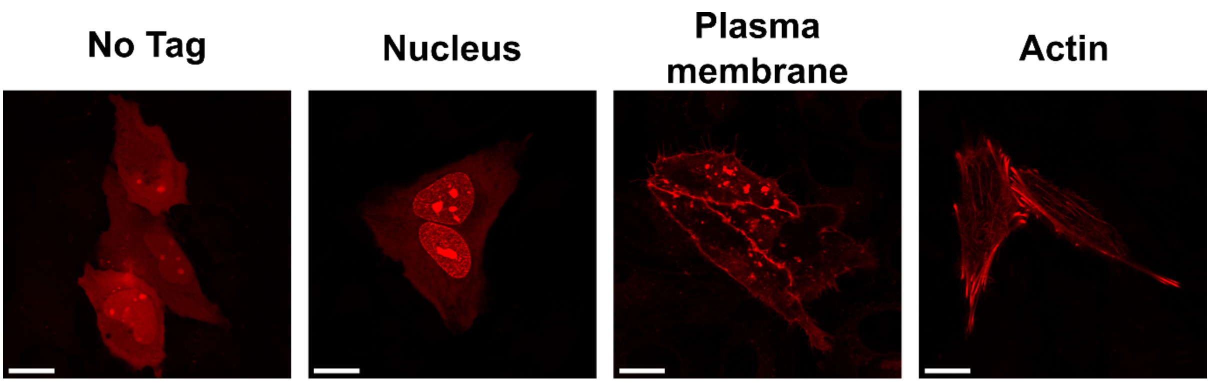


**Supplementary Figure 73.** Fluorescence confocal microphotographs of live HeLa cells expressing sc-apCC-4-SN38-1 fused to short N-terminal sequences targeting the nucleus, plasma membrane, and actin, but without the mEmerald domain. Images were obtained 30 minutes after staining with 0.1 μM Nile red added to the cell media and then observing Nile red in the red channel (ex/em 561/580-654 nm). Scalebar = 20 μm.

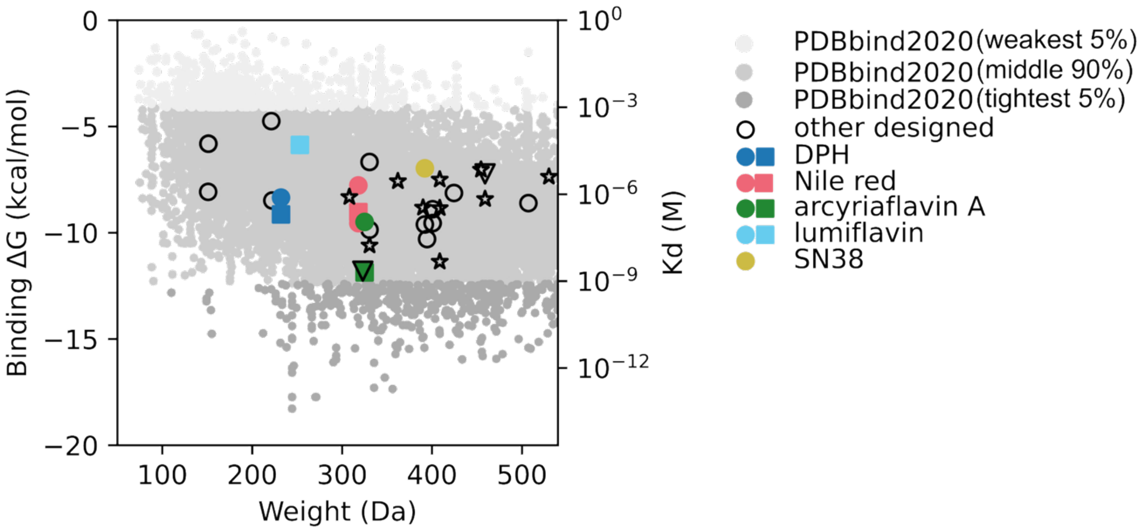


**Supplementary Figure 74. Comparison of binding affinities between RASSCCoL designed binders, natural binders from PDBbind2020[3] and *de novo* binders designed by others.** RASSCCoL designs in coloured circles, LigandMPNN optimised designs in coloured squares. 4-helix binders designed by COMBS in open triangles[4,5], LigandMPNN and diffusion-based binders in open stars[6-9]. Open circles denote older designs which were extracted by comparing PDBbind2020 with The Protein Design Archive[10]. Halogenated compounds and metal complexes excluded.
